# Reprogrammed tracrRNAs enable repurposing RNAs as crRNAs and detecting RNAs

**DOI:** 10.1101/2021.05.24.445356

**Authors:** Yang Liu, Filipe Pinto, Xinyi Wan, Shuguang Peng, Mengxi Li, Zhen Xie, Christopher E. French, Baojun Wang

**Author notes:** These authors contributed equally to this work (Filipe Pinto, Xinyi Wan). Correspondence can be addressed to BW.

## Abstract

In type II CRISPR systems, the guide RNA (gRNA) consists of a CRISPR RNA (crRNA) and a hybridized trans-acting CRISPR RNA (tracrRNA) which interacts directly with Cas9 and is essential to its guided DNA targeting function. Though tracrRNAs are diverse in sequences and structures across type II CRISPR systems, the programmability of crRNA-tracrRNA hybridization for particular Cas9 has not been studied adequately. Here, we revealed the high programmability of crRNA-tracrRNA hybridization for *Streptococcus pyogenes* Cas9. By reprogramming the crRNA-tracrRNA hybridized sequence, reprogrammed tracrRNAs can repurpose various RNAs as crRNAs to trigger CRISPR function. We showed that the engineered crRNA-tracrRNA pairs enable design of orthogonal cellular computing devices and hijacking of endogenous RNAs as crRNAs. We next designed novel RNA sensors that can monitor the transcriptional activity of specific genes on the host genome and detect SARS-CoV-2 RNA *in vitro*. The engineering potential of crRNA-tracrRNA interaction has therefore redefined the capabilities of CRISPR/Cas9 system.

## Introduction

The type II clustered regularly interspaced short palindromic repeats (CRISPR) system employs a small non-coding RNA known as trans-activating CRISPR RNA (tracrRNA) to form the mature dual-RNA structure of guide RNA (gRNA) (*1–5*). By complementary pairing with a precursor CRISPR RNA (pre-crRNA), the tracrRNA mediates RNase III-dependent RNA processing to generate mature crRNA and Cas9 ribonucleoprotein (RNP) complex (*3–8*). The pre-crRNA consists of continuous alternating spacers and repeat sequences (*9*). The unique spacers have homology to different foreign DNA sequences and guide the Cas9 RNP to recognize the target DNA by RNA-DNA pairing (*8, 10*), while the repeats of pre-crRNA can form an RNA duplex with the tracrRNA by hybridizing to an anti-repeat region near its 5’-end (*6*). Consequently, the crRNA:tracrRNA RNA duplex will be cleaved by RNA duplex-specific RNase III in the presence of Cas9, leading to fragmented pre-crRNA with one spacer for each mature crRNA (*4, 6–8*).

The tracrRNA plays a vital role in the maturation of gRNA and in the CRISPR function, and has molecular interactions with Cas9 protein (*4, 11, 12*). In many applications and studies regarding CRISPR, the crRNA and tracrRNA can be artificially fused into a single gRNA molecule called sgRNA (*13*). The 3’-end truncated crRNA and 5’-end truncated tracrRNA can be linked and still support the function of CRISPR/Cas9. The crystal structure of the Cas9 complex reveals that the sgRNA scaffold has strong interactions with Cas9 and plays an essential role in CRISPR function (*5*). Although the function of tracrRNA is conserved in the CRISPR systems of various species, its sequence and localization within the CRISPR-Cas locus are highly diverse (*8, 14*). It is also known that there is an orthogonal relationship between tracrRNAs and the corresponding Cas9 proteins from distant species (*8, 13*).

The relationship between the Cas9 and dual-RNA raises an intriguing question of whether the complementary pairing between crRNA and tracrRNA is programmable for particular Cas9 proteins. Several previous studies support the potential programmability of the crRNA-tracrRNA pairing. First, the Cas9 protein of specific bacteria can form a functional complex with tracrRNAs from closely related species (*8, 13*). In these studies, the available heterogeneous tracrRNAs have similar secondary structures but different sequences, suggesting that Cas9 proteins may be more sensitive to the secondary structure of the gRNAs than to their specific sequences. Second, the tetraloop stem of the sgRNA, equivalent to the crRNA-tracrRNA pairing fragment, is programmable for the *Streptococcus pyogenes* Cas9 (SpCas9) (*14, 15*). Assuming that the difference between sgRNA and dual-RNA will not change the Cas9-gRNA interaction model, the sequence of the crRNA-tracrRNA matching region should also be reprogrammable. Finally, previous studies reveal that the CRISPR/Cas9 system can tolerate single base-pair substitutions in the crRNA-tracRNA pairing region (*16*), suggesting multiple base-pair substitutions may also be feasible. Nonetheless, only recently has the programmability of the crRNA-tracrRNA hybridization region been explored to detect RNA molecules (*17*).

Here, we systematically explore the programmability of crRNA-tracrRNA pairing of the CRISPR/SpCas9 system, built on our initial intention of developing programmable AND logic devices. We validate the programmability of crRNA-tracrRNA and reveal a new set of principles to guide the design of CRISPR systems with reprogrammed dual-RNA. The high programmability of crRNA-tracrRNA pairing also brings new perspectives and potential for the engineering and application of the CRISPR/Cas9 tool. The orthogonal AND logic gates are developed based on this mechanism. Further, by reprogramming the crRNA-tracrRNA pairing, SpCas9 can specifically repurpose various RNAs as crRNAs for triggering CRISPR function. We develop an mRNA sensor able to hijack of endogenous RNA molecules as crRNAs and show that the inherent characteristics of mRNA and tracrRNA structures can strongly affect the mRNA-mediated CRISPR function. Notably, we successfully monitor the transcription level of endogenous genes in *E. coli* and connect the bacterial genetic network to an artificial gene circuit, which works as a whole-cell arsenic biosensor. A new type of RNA sensor is demonstrated by detecting SARS-CoV-2 target RNA *in vitro*.

This study has thus redefined the application capabilities of Cas9 proteins and the sources of crRNAs, and provides new scope for further studying the type II CRISPR systems.

## Results

### crRNA-tracrRNA hybridization is programmable and enables design of orthogonal AND gates

To study crRNA-tracrRNA pairing, we designed a crRNA-tracrRNA mediated CRISPR activation (CRISPRa) device in bacteria modified from a previously reported eukaryote-like CRISPRa system (*18, 19*).

A crRNA-tracrRNA mediated CRISPR activation (CRISPRa) device requires splitting the sgRNA into crRNA and tracrRNA. However, in the original design of our CRISPRa system, one of the two RNA aptamers occupies the tetraloop of sgRNA, equivalent to the structure where crRNA and tracrRNA are connected. Therefore, splitting the sgRNA would destroy this structure and the function of our CRISPRa device (**Figure 1a**). To overcome this issue, we redesigned the sgRNA by moving the RNA aptamers to the 3’-end of the sgRNA. As a result, we proved that the sgRNA with tail-fused aptamers is functional for the eukaryote-like CRISPRa system. Further, the 3-aptamers design provides higher efficiency than the 2-aptamers sgRNA design (**Supplementary Figure 1a, b**).

**Figure 1.**
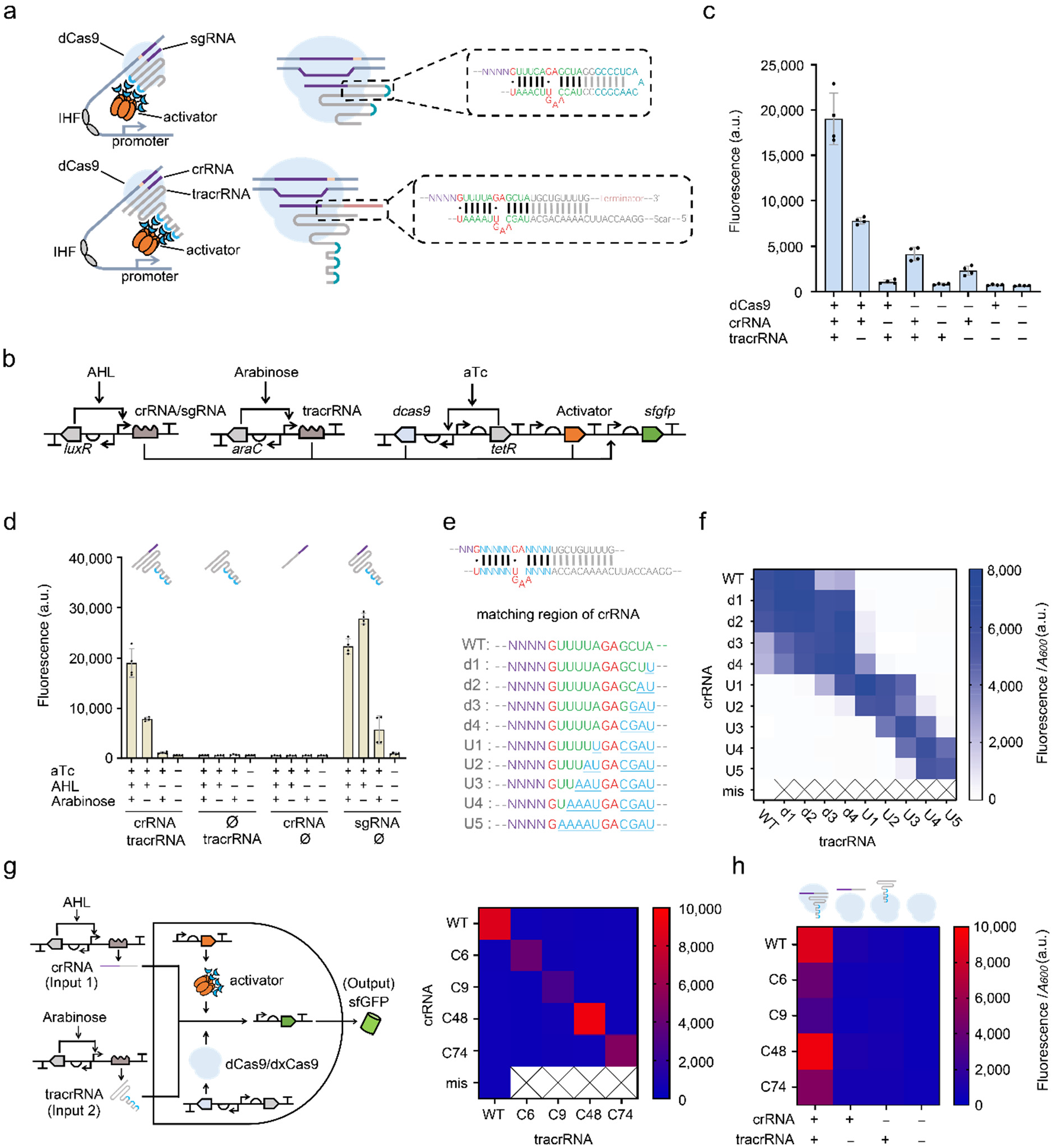
Dual-RNA mediated CRISPRa system revealed the programmability of crRNA-tracrRNA hybridization. **(a)** Schematics showing the original sgRNA mediated eukaryote-like CRISPRa system (top) and the dual-RNA mediated CRISPRa system (bottom). Green base pairs, the minimum structure required for CRISPR/Cas9 function; blue bases, RNA aptamer BoxB; red bases, the bulge structure and wobble base pairs within the crRNA-tracrRNA hybridized region. The short lines between the bases indicate the paired RNA bases. Two wobble base pairs are marked as solid black dots. **(b)** Circuit design of the crRNA-tracrRNA mediated CRISPRa system. Promoters P_*tet*_, P_*lux2*_, P_*BAD*_ drive the expression of dCas9, crRNA and tracrRNA, respectively. The activator PspFΔHTH::λN22plus is driven by constitutive promoter J23106 (*22*) (Anderson promoter collection). A σ^54^-dependent promoter with artificial UAS was employed to express the *sfgfp* reporter. **(c)** Test of the combination of three components in the crRNA-tracrRNA mediated CRISPRa. Inducer concentrations: 2.5 ng mL-1 aTc, 1.6 μM AHL, and 0.08 mM arabinose. Error bars, s.d. (*n* = 4). **(d)** Test of necessity of crRNA and tracrRNA, and comparison of the function of gRNAs before and after being split. Inducer concentrations: 2.5 ng mL^−1^ aTc, 1.6 μM AHL, and 0.08 mM arabinose. Combinations were achieved via presence (+) or absence (−) of inducers. Error bars, s.d. (*n* = 4). **(e)** Design of the library for testing reprogrammed crRNA-tracrRNA hybridized pairs. The blue letter ‘N’ indicates the substituted base pairs. The representation of motifs in the remaining sequences on the bottom is the same as those in **a. (f)** Orthogonality test of reprogrammed crRNA-tracrRNA pairs. The label ‘mis’ indicates the WT crRNA has a mismatched spacer (LEB3) with the target UAS (LEA2). All the other crRNAs have spacer LEA2 and a corresponding σ^54^-dependent promoter with UAS LEA2 was used for the reporter expression. dCas9 expression was under P_*tet*_ promoter, and the activator expression was under a constitutive promoter J23106. sgRNA was produced under P_*lux2*_ promoter. The reprogrammed tracrRNAs and crRNAs were combined in pairs. Inducer concentrations: 2.5 ng mL^−1^ aTc, 1.6 μM AHL, and 0.08 mM arabinose for dCas9, crRNA and tracrRNA induction respectively. Error bars, s.d. (*n* = 3) **(g)** Design of orthogonal AND gates based on reprogrammed crRNA-tracrRNA paring. The diagram on the left shows the design of the CRISPR-enabled AND gate circuit. The heat map on the right shows the outcome of an orthogonality test of the WT and 4 randomly generated crRNA-tracrRNA paired sequences. Inducer concentrations: 2.5 ng mL^−1^ aTc, 1.6 μM AHL, and 0.08 mM arabinose for dCas9, crRNA, tracrRNA induction respectively. Error bars, s.d. (*n* = 3) **(h)** Each orthogonal crRNA-tracrRNA mediated CRISPRa device displays the Boolean AND logic profile. The cartoon above shows the presence or absence of the components of the CRISPR complex under different induction conditions. The data in the active state and induction conditions are the same used in **g**. Error bars, s.d. (*n* = 3). a.u., arbitrary units.

Next, we split the sgRNA with 3’-end fused aptamers to tracrRNA and crRNA at the tetraloop position (**Figure 1a**). Apart from the nine essential RNA base pairs for the functional dCas9 RNP complex (*13*), additional base pairs with the same sequence as wild type (WT) crRNA and tracrRNA from *Streptococcus pyogenes* were designed to mimic the natural crRNA-tracrRNA pairing. The expression of dCas9, crRNA and tracrRNA were controlled by different inducible promoters (**Figure 1b**). An experiment permuting three induction conditions indicated that only when crRNA, tracrRNA and dCas9 were all induced did the CRISPRa system give the highest output (**Figure 1c**). When we removed the crRNA generator or tracRNA generator circuit, no activation of the CRISPRa system was observed. The activation efficiency of crRNA-tracrRNA mediated CRISPRa and sgRNA mediated CRISPRa are in the same order of magnitude (**Figure 1d**).

In the above tests, we confirmed that the dual-RNA mediated CRISPR system could also be employed for eukaryote-like CRISPRa in bacteria. Therefore, we set this crRNA-tracrRNA mediated CRISPRa device as a platform for the subsequent study of the programmability of crRNA-tracrRNA pairing in the CRISPR/Cas9 system. We first reprogrammed the base pairs of the core crRNA-tracrRNA hybridizing region. The core region is the minimal matching region of tracrRNA and crRNA for CRISPR function including 12 nt RNA at the repeat region of crRNA and the corresponding tracrRNA region (*13*). We introduced a stretch of mutations into the crRNA and made complementary mutations in tracrRNA for testing (**Figure 1e**).

Specifically, the core matching region includes nine RNA base pairs and two wobble base pairs (G-U). Two bases, GA, in crRNA and four bases, AAGU, in tracrRNA form a bulge structure, essential for CRISPR/Cas9 function (*14*) (**Figure 1e**). Previous studies have shown that the deletion or the replacement of the nucleotides in the bulge can abolish the sgRNA function (*14, 15*). Assuming that the same rule applies to crRNA-tracrRNA pairs, we reprogrammed the nine base pairs except the nucleotides of the bulge and wobble base pairs in the first step.

The library of the above reprogrammed crRNA-tracrRNA pairs was tested using our CRISPRa system. The results show that all the reprogrammed crRNA-tracrRNA pairs enabled efficient CRISPRa output. Further, all the outputs were in the same order of magnitude as that of the wildtype (WT) crRNA-tracrRNA hybridizing region (**Supplementary Figure 1c**). We then explored the orthogonality of mutated crRNA-tracrRNA pairs by recombining the mutated crRNAs and tracrRNAs. Interestingly, the bulge structure is like a dividing line. The upstream region near the bulge is less able to tolerate mismatches than the immediately downstream (**Figure 1f**). For SpdCas9, more than two adjacent mismatches in the upstream region of the bulge inhibit the CRISPRa function. By contrast, in the downstream region, even four mismatches could be tolerated for CRISPRa function. Surprisingly, in some cases, the mismatches in the downstream region enable higher CRISPRa output than that of the WT crRNA-tracrRNA pair (**Figure 1f**).

Regarding the original purpose of this work, the dynamic behavior of the dual-RNA mediated CRISPRa system shows the Boolean logic profile of AND gate, providing an output only when both the inputs are present (*20*). Although AND-gate circuits have various applications in synthetic biology (*21*), building complex artificial genetic networks requires a large number of highly orthogonal AND gates. CRISPRa based on the programmability of crRNA-tracrRNA pairing can efficiently address this problem.

In this case, the presence or absence states of the crRNA or tracrRNA represent states 1 and 0 of the inputs, respectively. The maximum and minimal expression levels of the sfGFP represent the states 1 and 0 of the output (**Figure 1g**). Our experiments have exhibited a typical AND-gate function of this system in *E. coli* (**Figure 1d**).

We then randomly generated four sequences for the crRNA-tracrRNA matching region and tested the orthogonality between these AND gates. As a result, we proved that orthogonality between the AND-gate circuits could be achieved based on the principle of complementary base pairing (**Figure 1g**).

As inducible systems are more desired to control logic gate devices (*18*), we employed the induced or non-induced states of the crRNA or tracrRNA to represent states 1 and 0, respectively. For the inducible system, this AND gate became less efficient due to its high sensitivity to tracrRNA leakiness from the inducible promoter (**Figure 1c**). To overcome this issue, we truncated the crRNA-tracrRNA pairing region to 14 bp to weaken their pairing affinity. As a result, this optimization improves the AND gates with good symmetry in response to the two inputs (**Figure 1h**).

In addition, inspired by our previous experience with dxCas9 3.7, and in how it could optimize the sensitivity to the sgRNA leakiness of our CRISPRa system (*18*), we speculated that the same strategy might also be applied to the crRNA-tracrRNA mediated CRISPRa. We first verified that dxCas9 could work with our newly designed sgRNA with tail-fused aptamers and functioned better than Cas9 (**Supplementary Figure 2**). Then, we proved that the dxCas9 could make the AND gates more symmetrical in response to the two inputs (**Supplementary Figure 3**).

### CRISPR activity with reprogrammed crRNA-tracrRNA pairing affected by multiple factors

For any engineering purpose based on reprogrammed tracrRNA-mediated CRISPR, the sequence preference of SpCas9 for the crRNA-tracrRNA hybridizing region is an important guide principle. Notably, previous research has showed that SpCas9 has direct intermolecular interaction with the dual-RNA hybridized fragments (*5*).

Since many related factors may affect the CRISPR function, we first investigated by varying the length of the RNA hybridization segment. We truncated the WT crRNA and WT tracrRNA simultaneously, making the length of the RNA hybrid segment gradually approach the minimum length for SpCas9 (**Figure 2a**). By characterizing them with CRISPRa, we noticed a dramatic decrease of the CRISPRa output when the paired length is shorter than 14 bp (including the two wobble base pairs). Conversely, the 14 bp length could support a similar output level to that from the WT version (**Figure 2a, Supplementary Figure 4**).

**Figure 2.**
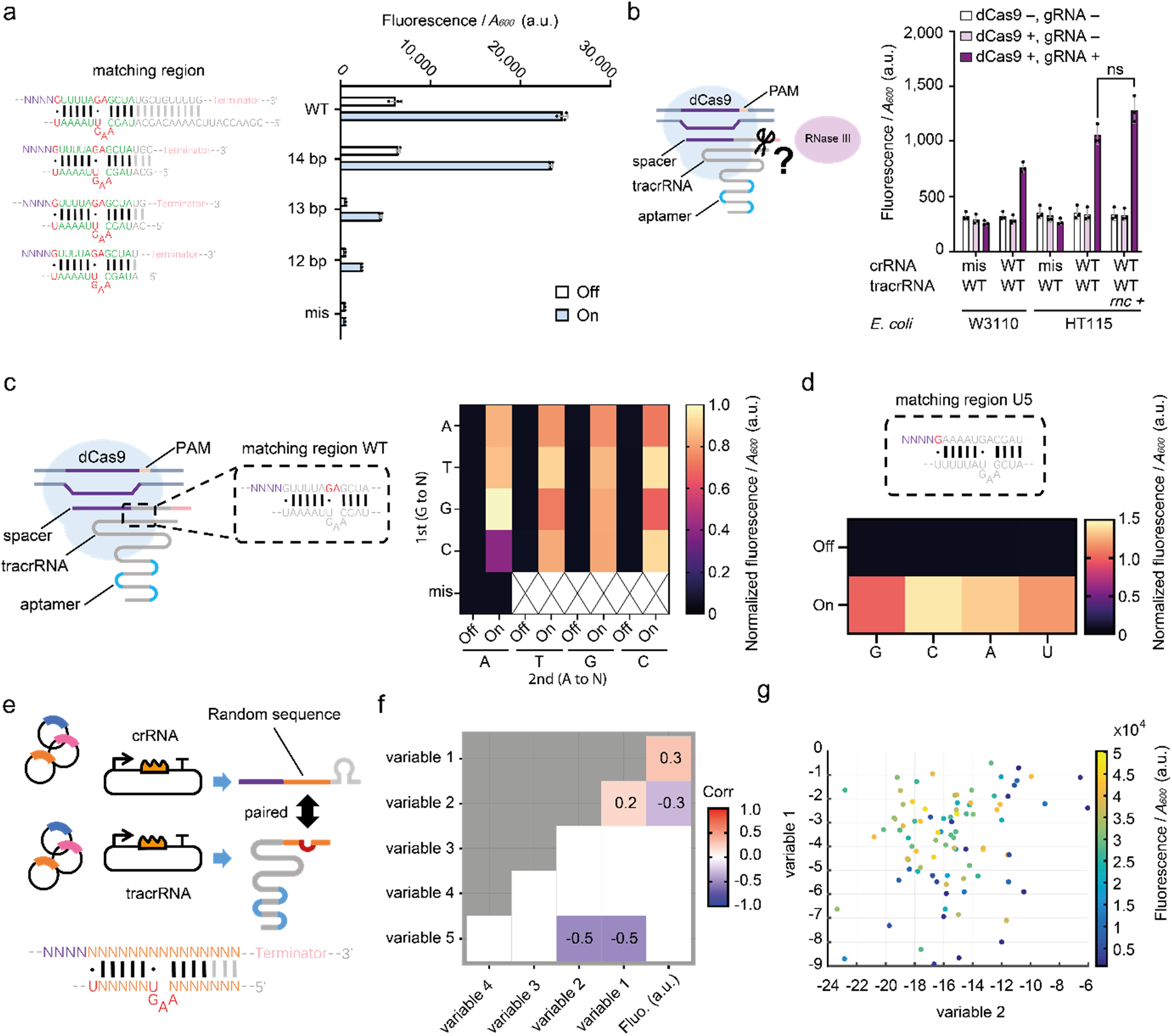
Mode of action of reprogrammed crRNA-tracrRNA pairs. **(a)** The length of the hybridized region affects the dual-RNA mediated CRISPRa function. The diagram on the left depicts the sequences used and the bar chart on the right exhibits the CRISPRa output. dCas9 was induced under P_*tet*_ promoter with 2.5 ng mL^−1^ aTc. The activator, crRNA and tracrRNA are all driven by the constitutive promoter J23106. The label ‘mis’ indicates the WT crRNA has a mismatched spacer (LEB3) with the target UAS (LEA2). All other crRNAs have spacer LEA2, and a corresponding σ^54^-dependent promoter with UAS LEA2 was used for reporter expression. Error bars, s.d. (*n* = 3). **(b)** RNase III is unnecessary for dual-RNA mediated CRISPRa. dCas9 was controlled by P_*tet*_ promoter, and *rnc*, crRNA and tracrRNA expression was driven by promoters P_*rhaB*_, P_*lux2*_ and P_*BAD*_ respectively. Inducer concentrations: 2.5 ng mL^−1^ aTc, 1.6 μM AHL, 0.33 mM arabinose and 0.4 mM rhamnose. The label ‘mis’ is the same as in **a**. Statistical difference was determined by a twotailed *t* test: *p* = 0.0577, *t* = 2.638. Error bars, s.d. (*n* = 3). **(c)** Saturated mutations of the GA site. The diagram on the left shows location of the GA site for saturated mutations. The heat map on the right shows the experimental results. Promoters P_*tet*_, P_*lux2*_ and P_*BAD*_ were used to drive the expression of dCas9, crRNA and tracrRNA respectively. The activator PspFΔHTH::λN22plus is driven by the constitutive promoter J23106. Inducer concentrations: 2.5 ng mL^−1^ aTc, 1.6 μM AHL and 0.08 mM arabinose for ‘On’ state, no induction for the ‘Off’ state. Data normalized by the output level from the ‘GA’ at ‘On’ state. *n* = 4. **(d)** The cartoon on the top shows the position of the base guanine in the core hybridized region between crRNA and tracrRNA. The U5 crRNA-tracrRNA pair was used as an original crRNA version, which had been shown as a functional crRNA-tracrRNA pair (see in **Figure 1f**). The G was replaced by C, A, and U. The heat map on the bottom shows the results of saturated mutations. The random sequence LEA2 was used here as the UAS and spacer. Induction conditions are the same as in **c**, and the data was normalized by the output level from the sample with the WT U5 ‘G’ at ‘On’ state. *n* = 4. **(e)** Schematic showing the randomized sequence library of crRNA-tracrRNA pairing. The orange bases show the paired randomized sequences, and purple ‘N’s stand for the spacer region, a fixed sequence (LEA2) among all variants. The red bases in tracrRNA are not changed. **(f)** Pearson correlation coefficient between individual features (variables) and the output fluorescence level. Blank indicates no significant correlation (*p* > 0.05). Numbers represent correlation coefficients. **(g)** Reprogrammed tracrRNA-crRNA library represented by variable 2 for crRNA-tracrRNA heterodimer binding and variable 1 for crRNA folding. Each dot represents a sample. Samples are coloured according to their fluorescence level. *n* = 3; a.u., arbitrary units.

Another question is whether RNase III is necessary for dual-RNA-mediated CRISPRa. For our design, the 3’-end of the crRNA has a terminator sequence, and the 5’-end of the tracrRNA retains a wild-type residual fragment. Both the crRNA and tracrRNA have ends longer than mature dual-RNA processed by RNase III. In previous studies, RNase III was shown to be indispensable for the maturation of crRNA and the immune function of Cas9 (*6*). Considering that RNase III has a preference for target double-strand RNA sequence, if RNase III plays a crucial function in our system, then the reprogramming of the crRNA-tracrRNA hybrid sequence may indirectly affect CRISPR function via RNase III. To verify this issue, we introduced our dual-RNA-mediated CRISPRa device into *E. coli* strains W3110 and its RNase III gene (*rnc*) knockout strain HT115. Our results show that RNase III is not required for our CRISPRa system. Furthermore, we complemented strain HT115 with a copy of the *rnc* gene expressed from a plasmid. However, this did not significantly improve the function of CRISPRa (**Figure 2b**). This result implies that RNase III will not be a limiting factor for crRNA-tracrRNA reprogramming.

Next, we tested the importance of the three bases in the wobble base pairs (G - U) and the bulge in crRNA via two additional experiments. By introducing saturation mutation at the GA site of the WT hybridization sequence, and at the first wobble base pair (G - U) of a functional hybridization sequence U5, we confirmed that changing these compositions will not completely disrupt CRISPRa function, although a few mutations can lead to changes in CRISPR activity (**Figure c, d**).

Since the results above provide a conveniently simplified crRNA-tracrRNA model for further research, we built a library with 90 paired candidates, including random RNA sequences in the crRNA-tracrRNA matching region (**Figure 2e**). For the library design, we ruled out the presence of NGG immediately downstream of the spacer to prevent a CRISPRi effect on the crRNA generator. The CRISPRa output from these paired crRNA-tracrRNA candidates was collected and combined with the other selected variables for statistical analysis.

Surprisingly, about 69 % of candidates show higher than 50 % of the activity of the WT sequence, and about 50 % of candidates show higher than 75 % of the activity of the WT sequence. About 22 % of candidates even show higher activity than the WT sequence (**Supplementary Figure 5**).

To explore potential influencing factors that may affect the activity of these reprogrammed tracrRNA-crRNA pairs, we first manually extracted the following features for each tracrRNA-crRNA sequence: the minimum free energy (MFE) of the crRNA optimal secondary structure (variable 1), change in Gibbs free energy (ΔG) of crRNA-tracrRNA heterodimer binding (variable 2), homology of the crRNA-tracrRNA matching region (variable 3), sequence similarity between the crRNA and the target DNA including the sequence downstream of the PAM site (variable 4) and GC content of the matching region (variable 5). Then we calculated the Pearson correlation coefficients between these features and output fluorescence.

**Figure 2f** shows that only variable 2 and variable 1 have a significant correlation with output fluorescence (*p* < 0.05). We next built a linear regression model using these two variables to reveal their relationship. Although the R^2^ value of the linear model is only 19%, the available data still suggests a trend, which is in line with our understanding of the CRISPR/Cas9 complex formation. The function of CRISPRa tends to be promoted by high affinity between crRNA and tracrRNA (low ΔG for heterodimer binding between crRNA and tracrRNA) and weak secondary structure of crRNA itself (high free energy of the thermodynamic ensemble for crRNA folding) (**Figure 2g**).

### Reprogrammed tracrRNA able to repurpose mRNA as crRNA

The programmability of the crRNA-tracrRNA pairing demonstrated above indicates the exciting potential of hijacking various RNAs as crRNAs. The natural crRNA mainly includes two sequence motifs: the spacer matched with target DNA and the downstream repeat matched with tracrRNA. Although pre-crRNA processing is necessary for CRISPR function (*6, 23*), the engineering of sgRNA proves that the 3’-end of crRNA near Cas9 is unnecessary for Cas9 function (*4*). In addition, the extended 5’-end of sgRNA and extended 3’-end of crRNA do not disrupt the activity of CRISPR/Cas9 (*22, 24, 25*).

Combining the facts that the spacer sequence itself can be reprogrammed to target different DNA sequences, and that the downstream repeat region is also programmable, then the entire crRNA sequence should be programmable. Conversely, any RNA sequence may become crRNA through dual recognition by programmed tracrRNA and target DNA.

To verify the above hypothesis, we randomly selected three GA sites on the mRNA encoding red fluorescent protein (RFP), since GA provided the highest function in our saturation mutation test (**Figure 2d**). The NGG adjacent to the assumed spacers was deliberately avoided. According to the context sequence adjacent to the GA sites, we designed three corresponding tracrRNAs and three cognate σ^54^-dependent promoters. Each of them has the UAS matching the assumed spacer sequence, which is the upstream mRNA region of the predicted mRNA-tracrRNA hybridization position. If the mRNA of RFP can be hijacked as crRNA, the mRNA should activate the CRISPRa device with dCas9 and corresponding tracrRNAs (**Figure 3a**).

**Figure 3.**
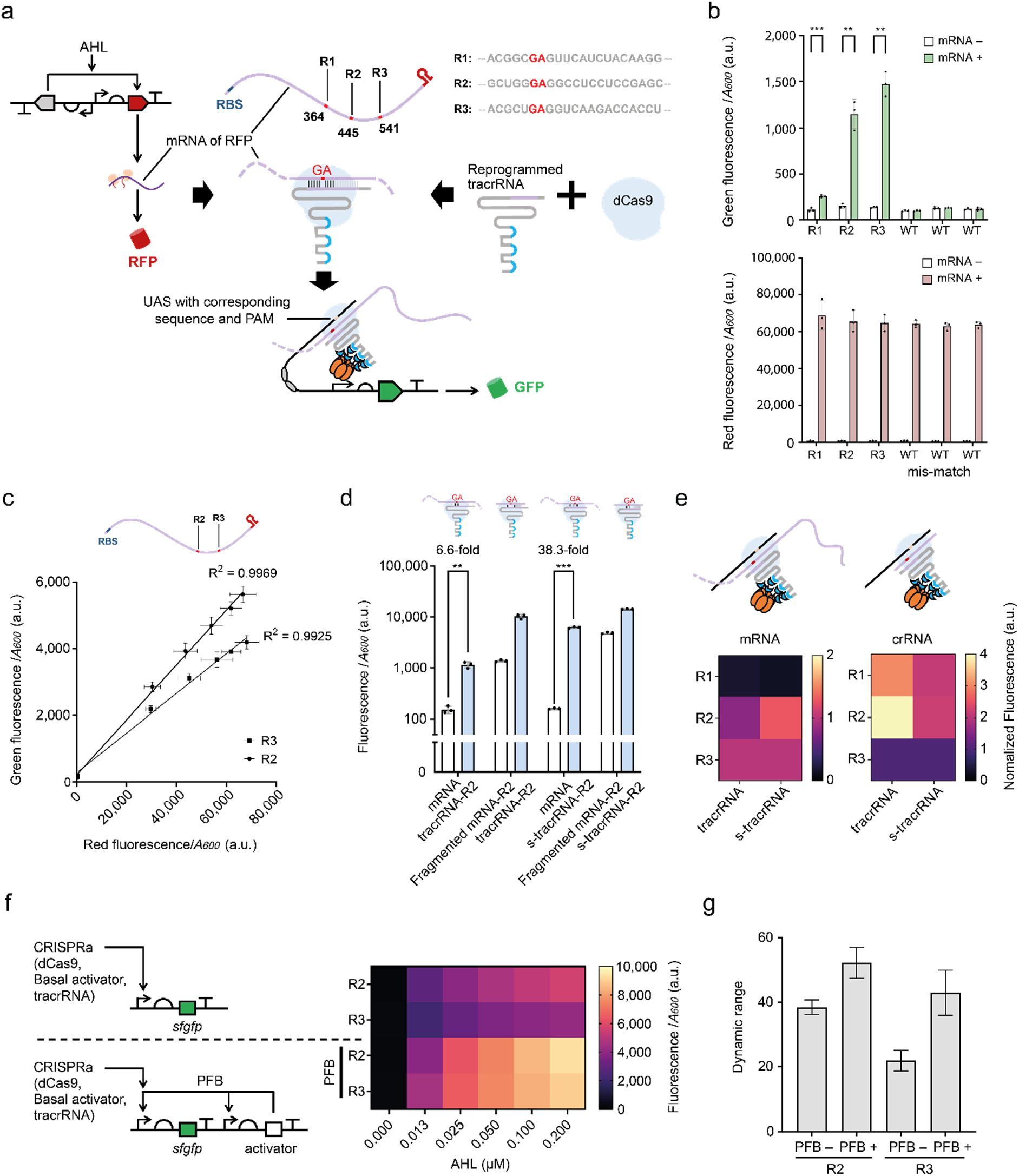
Engineered tracrRNAs repurposed mRNAs into crRNAs and activated CRISPRa. (a) Schematic showing the design of the mRNA sensor by hijacking mRNA as crRNA. Three randomly selected RNA sites with a GA motif (red symbol) are represented on the mRNA. The sequences of three target hybridizing sites are shown in grey letters with GA site highlighted in red. The upstream RNA sequences immediately adjacent to the tracrRNA target hybridizing site were used as the design templates for the UASs. Accordingly, the three synthetic promoters can be activated by the three cognate CRISPRa complexes comprising the mRNA, with one example shown (bottom). (b) Green (top) and red fluorescent outputs (bottom) of the mRNA-mediated CRISPRa. WT tracrRNA was used as a control. dCas9 and activator expression were driven by the P_*tet*_ and P_*rhaB*_ promoters. The mRNA of RFP and the tracrRNA were transcribed from P_*lux2*_ and P_*BAD*_, respectively. Inducer concentrations: 2.5 ng mL^−1^ aTc, 0.2 mM rhamnose, and 0.08 mM arabinose; 100 nM AHL used for mRNA expression. Statistical difference was determined by a two-tailed *t* test: R1, *p* = 0.0004, *t* = 11.01, or by a Welch’s *t* test: R2, *p* = 0.0070, *t* = 10.91; R3, *p* = 0.0032, *t* = 17.13. Error bars, s.d. (*n* = 3) (c) Scatter plot shows the linear relationship between the expression levels of RFP and sfGFP under varying AHL inductions by fitting to a linear model. For R2, Equation: *GF* = 0.08230×*RF* + 216.5 (*GF*: green fluorescence, *RF:* red fluorescence); The coefficient of determination (R^2^) is 0.9969; *p*-value <0.0001. For R3, Equation: *GF* = 0.05937×*RF* + 282.8 (*GF*: green fluorescence, *RF*: red fluorescence); The coefficient of determination (R^2^) is 0.9925; *p*-value <0.0001. Error bars, s.d. (*n* = 3) (d) 5’ end truncation on tracrRNA improved mRNA-mediated CRISPR function. The cartoon shows the test of component combinations: mRNA fragment or whole mRNA as crRNA with 5’-end extended tracrRNA or 5’-end truncated tracrRNA (s-tracrRNA). The mRNA fragment used has the same sequences as that in the mRNA target site. R2 used as controls. For the combination of mRNA and tracrRNA-R2, the same data were used as the data in b. Inductions are the same as in b. Statistical difference was determined by a two-tailed Welch’s *t* test: mRNA + s-tracrRNA-R2, *p* = 0.0005, t = 43.63. Error bars, s.d. (*n* = 3) (e) The same sequence from mRNA fragment and whole mRNA resulted in altered CRISPR functions. The raw data obtained from R2 site are the same as that in d, and inductions are the same as in b and d. In each group, the data are normalized by the CRISPRa output level of the sample with R3 site. (f) Diagram shows the configuration of the original reporter circuit and the version with where a positive feedback loop (PFB) added, with results shown on the right. The input components were controlled by the same promoters used in b. An AHL gradient (0.2, 0.1, 0.05, 0.03, 0.01, 0 μM) was used for mRNA expression. Inducer concentrations: 2.5 ng mL^−1^ aTc, 0.2 mM rhamnose, and 0.08 mM arabinose. (g) Dynamic range calculated from f. Error bars, s.d. (*n* = 3). a.u., arbitrary units.

For each tracrRNA target site on the mRNA, we used the WT tracrRNA, which should not match the RFP mRNA, as a control. Our result shows that, when employing mRNA-paired tracrRNAs, there was a positive linear correlation between the expression level of RFP and GFP for the R2 and R3 sites. This correlation disappeared when using WT tracrRNA (**Figure 3b, c**). The three randomly selected tracrRNA target sites gave different CRISPRa efficiencies. The output from site R1 is weak, and those from sites R2 and R3 are relatively stronger (**Figure 3b**). For all the sites, when using the corresponding sequences in isolation as a short crRNA, the CRISPRa output intensities obtained were greater than using the whole mRNA (**Figure 3d, Supplementary Figure 6**). We also found that the relationship of CRISPRa strength between different target sites is dependent on the inherent characteristics of mRNA. When the target sequences were fragmented as crRNAs, this strength relationship changes dramatically.

It is tempting to speculate that the translation process might cause the above output discrepancy between mRNA and isolated fragments. However, surprisingly, the binding of the CRISPR complex to mRNA did not seem to interfere with RFP translation (**Figure 3b**). Conversely, to assess whether the translation process could interfere with the CRISPRa system, a version of mRNA lacking the ribosome binding site (RBS) was used to test the impact of translation on CRISPRa. The experimental results show that the level of RFP was indeed reduced due to the lack of RBS, but the output of mRNA-mediated CRISPRa did not change obviously (**Supplementary Figure 7**). In addition, we also found that the length of the mRNA-tracrRNA pairing region can affect the CRISPRa efficiency, and for different targets on mRNA, the optimal length is also different (**Supplementary Figure 8**).

Although the above results show that, for hijacking mRNA, the CRISPR function of a particular target site is unpredictable, we nevertheless explored some design principles based on two exciting facts we discovered. First, a simplified tracrRNA design can improve the performance of this mRNA monitor on all the sites. Our result shows that when we deleted the redundant 5’-end of tracrRNA, the strength and dynamic range of CRISPRa are greatly improved (**Figure 3d, Supplementary Figure 6**). Second, the optimization of tracrRNA did not change the strength pattern of a set of sites on the mRNA. We compared the patterns when CRISPRa is combined with mRNA or fragmented mRNA, with or without tracrRNA optimization. We found that the patterns based on fragmented mRNA and complete mRNA are not similar, however, the patterns based on presence or absence of optimization maintain the similarity when from the mRNA or fragmented mRNA, respectively. This suggests that the inherent characteristics of mRNA such as secondary structure may robustly affect the function of mRNA-mediated CRISPRa (**Figure 3e**), and also suggests that site selection and tracrRNA optimization may be independent optimization methods.

Finally, we combined the design strategies developed to optimize the mRNA sensor *in vivo*. To amplify the output signal, we designed a positive feedback loop (PFB) (Figure 3f). One more σ^54^-dependent promoter activated by the mRNA was utilized to express additional engineered activator protein for enhancing CRISPRa. The result shows that the PFB circuits could amplify the CRISPRa output and dynamic range based on different mRNA-tracrRNA matching sites (Figure 3g). The dynamic range increases from 9.6-fold of the original device to 52.2-fold with PFB design through the above engineering methods.

### Monitoring endogenous RNAs via reprogrammed tracrRNA-mediated CRISPRa

Successful hijacking of the RFP mRNA as crRNA raises an interesting question: is the same strategy available for endogenous RNAs transcribed from the genome? To explore this possibility, we chose the *ars* operon of *E. coli* as a target, since the promoter P_*arsR*_ can be induced by sodium arsenite (NaAsO_2_), a toxic environmental pollutant (*26, 27*).

Having previously shown that different sites on the mRNA may have different availabilities, we tested some candidate sites on the *ars* transcripts on artificial circuits first. The arsenic responsive gene cluster (*arsRBC*) was isolated and cloned into a vector under inducible promoter P_*lux2*_. Four candidate sites were chosen and made into short fragments of mRNA (**Figure 4a**).

**Figure 4.**
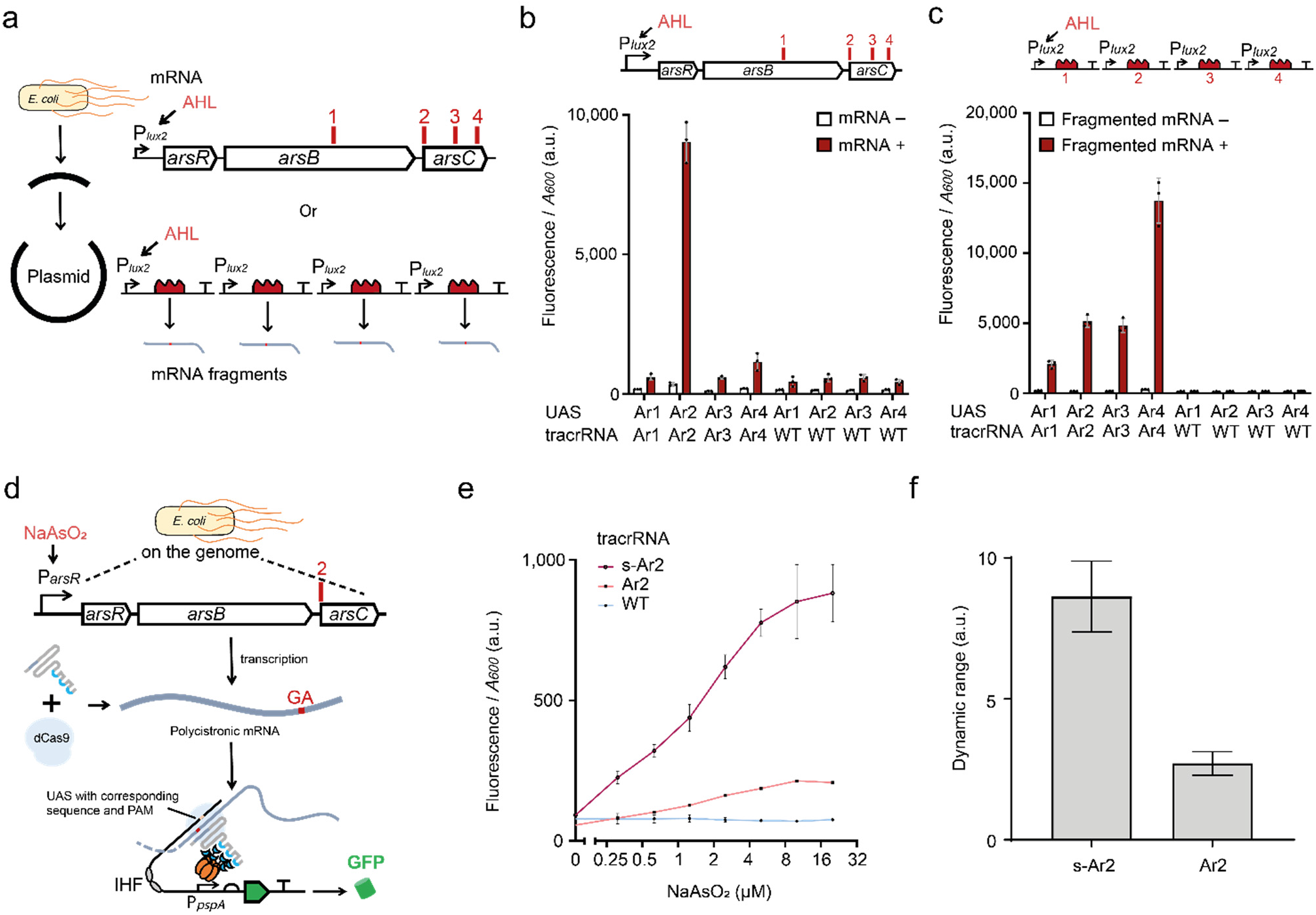
Hijacking of endogenous mRNA as crRNA to detect arsenic induced transcription. **(a)** Schematic showing the genetic sites selected for crRNA targeting. The DNA encoding the endogenous arsenic responsive pathway in *E. coli* was isolated and inserted into a plasmid under P_*lux2*_ promoter. In parallel, the four candidate sites were also extracted separately and inserted into the same vector backbone. The red numbers and short lines show positions of the four candidate sites (Ar1-Ar4) **(b)** Test of the availability of the candidate sites on the plasmid transcribed mRNA. The mRNA expressed by the P_*lux2*_ promoter was then bound to different tracrRNA and promoters. Expression of dCas9 and activator were driven by P_*tet*_ and P_*rhaB*_ promoters. The tracrRNA were transcribed from P_*BAD*_. Inducer concentrations: 2.5 ng mL^−1^ aTc, 0.4 mM rhamnose, 0.1μM AHL and 0.08 mM arabinose. Error bars, s.d. (*n* = 3) **(c)** Test of the availability of candidate sites on the fragmented mRNAs as crRNAs. crRNAs expressed by the P_*lux2*_ promoter were bound to corresponding tracrRNAs and promoters. Inductions are the same as in **b**. Error bars, s.d. (*n* = 3) **(d)** Schematic showing mechanism of hijacking the endogenous mRNA of the arsenic responsive operon of *E. coli* to activate reporter expression by CRISPRa. Red symbols indicate the Ar2 targeting site in the polycistronic mRNA of the arsenic-related gene cluster. The upstream RNA sequence immediately adjacent to the tracrRNA target was used as the design template for the upstream activating sequence (UAS) for CRISPRa activation. **(e)** CRISPRa outputs of hijacking endogenous mRNA as crRNA. ‘Ar2’ stands for the sensor with originally designed tracrRNA. The ‘s-Ar2’ stands for the sensor with 5’-end truncated tracrRNA. ‘WT’ indicates that a WT tracrRNA which cannot match the mRNA was used here. Expression of the dCas9 and activator were driven by P_*tet*_ and P_*rhaB*_ promoters. The tracrRNA were transcribed from P_*BAD*_. Inducer concentrations: 2.5 ng mL^−1^ aTc, 0.4 mM rhamnose, and 0.08 mM arabinose.A gradient of sodium arsenite (0, 0.31, 0.63, 1.3, 2.5, 5, 10, 20 μM) was used for inducing transcription of the arsenic responsive gene cluster. Error bars, s.d. (*n* = 3). **(f)** Dynamic range calculated from **e**. Error bars, s.d. (*n* = 3). a.u., arbitrary units.

We tested the availability of these sites by using programmed corresponding tracrRNAs and promoters. The results showed that only Ar2 is an available target on the entire *ars* mRNA. Yet, for the fragmented mRNA, different availabilities of the mRNA sites were revealed, similar to what we have observed in hijacking RFP mRNA. Again, this implies that the characteristics of mRNA itself can strongly affect the function of mRNA-mediated CRISPR (**Figure 4b, c**).

Next, we used the selected Ar2 site to monitor the activity of the *ars* operon on the *E. coli* genome. The reporter circuit and a tracrRNA-Ar2 generator were transformed into *E. coli* (**Figure 4d**). We constitutively induced the expression of dCas9, tracrRNA, and activator in *E. coli*. When we set a gradient of NaAsO2 for a group of *E. coli* cultures, as expected, for tracrRNA-Ar2, an output increase was detected with increasing NaAsO_2_ concentration (**Figure 4e**). In this case, the output signal is much weaker than that from the artificially expressed mRNA, which may be due to the relatively low concentration of endogenous RNA and the different spatial locations of the transcription in the cell. Finally, we tried to optimize this sensor by simplifying the 5’-end of tracrRNA, and proved that it can improve the performance of the mRNA monitor (**Figure 4e, f**).

Overall, we confirm that endogenous mRNA can be hijacked by Cas9, and in this way, it allows us to monitor genomic transcriptional activity and connect the cellular gene regulatory network to an artificial actuating or reporting gene circuit.

### CRISPR/Cas9-operated RNA detection *in vitro*

An engineering method that can convert non-crRNA into crRNA undoubtedly has great application potential for nucleic acid sensing. Here, we developed a novel programmable RNA sensor with unique dual recognition characteristics, which theoretically has a higher specificity than the DNA recognition of the CRISPR/Cas9 complex. Compared with other well-known CRISPR RNA sensors such as SHERLOCK, HOLMES and DETECTR that can continuously cleave non-specific singlestranded nucleic acid (*28–30*), Cas9 cleavage lacks such signal amplification effect. In contrast to Cas12a and Cas13, Cas9 can only cleave specific target DNA and then remains bound to it, resulting in insufficient repetitive cleavage of the same target DNA (*31*).

In order to overcome the above shortcomings of Cas9-directed RNA sensing, we designed an *in vitro* transcription-based reporting system with a signal amplification effect. We named it the <u>C</u>RISPR-<u>o</u>perated <u>N</u>ucleic acid <u>A</u>mputation <u>N</u>otification (CONAN) system. It consists of a reporter DNA named CONAN DNA, fluorogen DFHBI, purified Cas9, and reprogrammed tracrRNA for sensing target RNA in an *in vitro* T7 expression system (*32*) (**Figure 5a**). The CONAN DNA can express an inactive Broccoli RNA aptamer which is blocked by a 3’ end secondary structure. When it senses the target RNA, the Cas9 will be activated and destroy the 3’ end secondary structure on the CONAN DNA, which will then continuously transcribe a functional aptamer that can bind to DFHBI, leading to an amplified fluorescent output signal (*33*). Due to the strong transcriptional activity of the T7 promoter, once a cleavage event occurs, Broccoli RNA can continue to accumulate in the cell-free system until it reaches a balance with the RNA degradation rate (**Figure 5a**).

**Figure 5.**
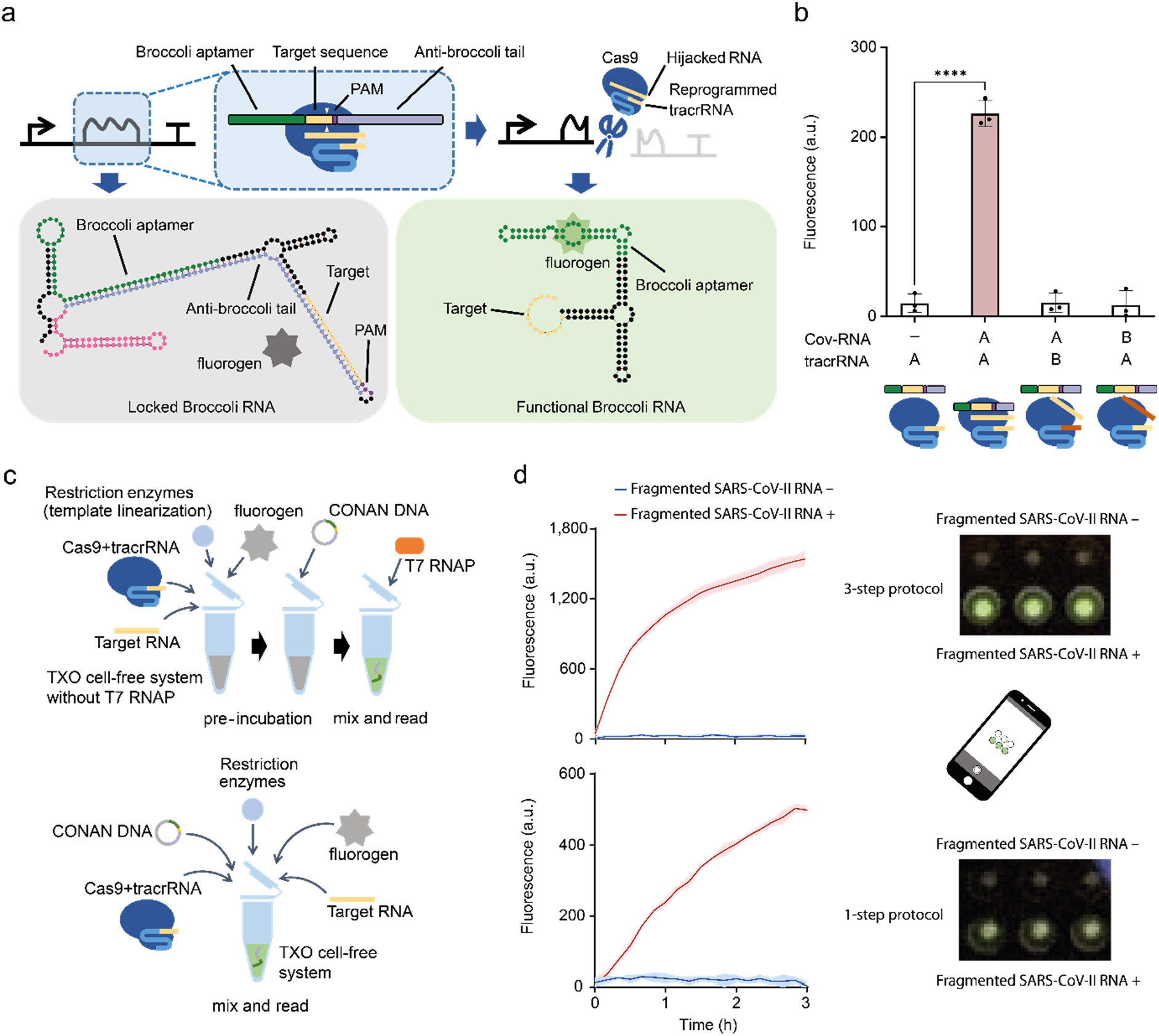
CRISPR-operated nuclear acid amputation notification (CONAN) system. **(a)** Schematic showing the design and mechanism of CONAN sensor. In the box surrounded by the dashed line, the green bar indicates the Broccoli RNA sequence, and the yellow bar indicates the CRISPR target DNA site paired with the target RNA. The dark purple bar indicates PAM. The light purple bar indicates an anti-Broccoli tail with an inverted repeat sequence of the Broccoli. The two white triangles mark the position where CRISPR/Cas9 cleaves the CONAN reporter DNA. The cartoon below shows the secondary structure of the RNA aptamer predicted by mfold Web Server (*34*). Polygonal star indicates the fluorogen which is activated when it binds to the Broccoli aptamer. The colours in the RNA structure diagram are consistent to those in the box above, and the pink segment indicates the T7 terminator. **(b)** Specificity test of CONAN. Two different genomic RNA fragments (fragments A and B) from the SARS-CoV-2 virus and the corresponding reprogrammed tracrRNAs were used to test the specificity of CONAN. The T7 *in vitro* transcription system was used with addition of 5 ng μL^−1^ CONAN DNA, 50 nM synthetic CoV-SARS-2 RNA fragment, 50 nM synthetic tracrRNA and 50 nM SpCas9. Data were collected 3h following incubation. Statistical difference was determined by a two-tailed *t* test: R1, *p* < 0.0001, *t* = 20.57. **(c)** Schematic showing the procedure of CONAN sensor to detect target coronavirus (SARS-CoV-2) RNA fragment. The reaction is carried out either with (top) or without (bottom) pre-incubation (see **METHODS**). **(d)** Dynamic output responses (left) and mobile phone images (right) of the CONAN sensor results responding to SARS-CoV-2 RNA fragment with (top) or without (bottom) pre-incubation. Phone images were acquired at the end of the reactions (overnight, see **METHODS**). The T7 *in vitro* transcription system is the same as in **b**. Error bars, s.d. (*n* = 3); a.u., arbitrary units.

We tested the CONAN sensor using synthetic coronavirus SARS-CoV-2 RNA fragments with corresponding reprogrammed tracrRNAs and Cas9. In this case, we designed a Cas9 cleavage site A between the Broccoli RNA coding sequence and the anti-Broccoli tail in the CONAN DNA. Only when the SARS-CoV-2 RNA fragment A matched the CONAN DNA and corresponding tracrRNA A, did CONAN give a significant report signal (**Figure 5b**). Otherwise, even if the CONAN DNA matched the target RNA, there was no false-positive signal. This dual recognition mechanism ensures the high specificity of CONAN detection.

Two different protocols, one-step and three-step protocol were designed to further test and optimize the CONAN system (**Figure 5c**). Upon sensing the SARS-CoV-2 RNA, we observed fluorescence reported by the CONAN sensor within 10 min (**Figure 5d**). The results also indicate that pre-incubation of CONAN DNA with CRISPR/Cas9 could improve the CONAN sensor’s output dynamic range. After incubation in the plate reader, we directly photographed the reaction vessel with a cell phone in the fluorescence imaging box. The broccoli RNA concentration in the microwells had accumulated enough to be distinguished by the naked eye and the cell phone camera (**Figure 5d**), which indicates that CONAN has the potential to be developed into a portable rapid diagnostic tool.

## Discussion

In this work, we investigated the programmability of crRNA-tracrRNA pairing in the CRISPR/Cas9 system based on our previously engineered eukaryote-like CRISPRa device. We systematically revealed the tolerance of SpCas9 protein for reprogrammed sequences of the crRNA-tracrRNA hybridizing region and found that all the nine base pairs necessary for the CRISPR function can be replaced without affecting the CRISPR function significantly. We showed that mismatches can confer orthogonality to crRNA-tracrRNA mediated CRISPRa and design of orthogonal AND logic devices. Interestingly, the mismatch tolerance of different segments of the crRNA-tracrRNA hybridizing region is uneven. The segment close to the spacer is more sensitive to mismatches than the part far away from it. For the RNA duplex between the spacer and bulge, two continuous mismatches or more are sufficient to disrupt the function of CRISPRa.

We used a paired crRNA-tracrRNA hybridization library to study the sequence preference of the SpCas9 crRNA-tracrRNA hybridization region. Two key factors, the affinity between crRNA and tracrRNA and the crRNA secondary structure, were identified. It is worth noting that different Cas9 versions may have different tolerance and preference to the reprogrammed crRNA-tracrRNA hybridizing sequence and mismatches. During our previous research, compared with dCas9, dxCas9 shows a lower tolerance for continuous mismatches at specific sites in the crRNA-tracrRNA pairing region (*18*). The sequence preference of different Cas9 proteins for the crRNA-tracrRNA hybridizing region may need to be assessed by combining machine learning and high-throughput methods case by case.

However, according to various practical scenarios we have covered here (i.e., endogenous RNA hijacking or for the AND gate devices), there are other factors than the aforementioned sequence preferences which may affect the CRISPRa function. Hence it is not easy to predictably select the best RNA target sequence. This reinforces the significance of further research on sequence preference by combining other high-throughput methods and machine learning with only a standardized library. Further studies are still required for further understanding of this issue.

Since CRISPR/Cas9 can tolerate extensions of the 5’-end and 3’-end, with enough length, any RNA may become a crRNA by binding to its complementary tracrRNA. Our experiments confirmed this possibility by hijacking of plasmid-transcribed mRNA and endogenous mRNA molecules as crRNAs to trigger CRISPRa function. This is consistent with another recent study (*17*).

Using the CRISPR/Cas9 system to recognize RNA molecules is an engineering field worth further exploring. However, there are some questions remained to be answered. Different targets on the same mRNA and different tracrRNA lengths on the same target have led to different CRISPRa efficiencies in our experiments. It is unclear whether this was due to RNA secondary structure, translation interference, RNase III, or other unknown factors. In our research, the RBS deletion of the plasmid-transcribed RFP mRNA did not significantly affect CRISPRa output, reflecting the counter-intuitive weak effect of translation on mRNA-mediated CRISPR function. Additionally, whether RNase III can process the mRNA-tracrRNA remains unknown, affecting our assessment about whether the secondary structure of the downstream sequence of the target RNA will affect CRISPR function.

We realized that the same system might work in eukaryotes. However, in eukaryotic cells, the localization of endogenous RNA in different regions is an unfavorable factor, affecting the function of endogenous RNA-based CRISPR tools. The tolerance of dCas9 for gRNA sequence diversity also raises an interesting question. Considering that Cas9 binding to non-canonical RNA has recently been reported in bacteria (*17*), would it be possible for endogenous RNA in eukaryotic cells to coincidentally form a functional CRISPR complex with Cas9, and then cause an off-target effect? In short, exploring the interaction between endogenous RNA and Cas9 in eukaryotes will lead to some new topics that are worth exploring.

Finally, we have developed a new cell-free reporter system CONAN with a signal amplification effect on our RNA sensor based on the programmability of crRNA-tracrRNA hybridization. Compared to the recently developed CRISPRi-based *in vitro* sensing system, CONAN is transcription only and therefore the response time is much shorter upon sensing (*17*). Moreover, it only amplifies the output signal when it senses target RNA and has much lower background signal. Thus it will be preferable for broad biosensing applications. This system can be used for CRISPR/Cas9-mediated RNA sensors and for enhancing or amplifying the signal of other types of CRISPR-based RNA sensors, as well as for many other research and application scenarios related to DNA fragmentation.

In summary, the programmability of crRNA-tracrRNA hybridization and the RNA recognition capability of the CRISPR/Cas9 system opens up a significantly broader picture of CRISPR/Cas9 engineering. For example, this programmable system may be utilized to sense RNA *in vivo* or *in vitro*, to build RNA editors, to reconstruct the topology of the genetic regulatory network, to mark transcripts in situ, or to build complex cellular computing circuits. Our findings have elevated our knowledge of CRISPR systems and will expand current CRISPR technologies in the future.

## Methods

### Strains and growth conditions

Unless otherwise specified, all the assays were performed in *E. coli* strain MC1061Δ*pspF*. The *E. coli* MC1061Δ*pspF* was generated through P1 phage transduction, using *E. coli* strain BW25113Δ*pspF*739::kan from the Keio collection as the donor strain (*35*). For circuit construction, *E. coli* TOP10 and *E. coli* DH5α were both employed in this study.

For bacterial culture, we cultured *E. coli* in Lennox’s Lysogeny Broth (LB-Lennox) medium (10 g L^−1^ peptone (EMD Millipore), 5 g L^−1^ yeast extract (EMD Millipore), 5 g L^−1^ NaCl (Fisher Scientific)) with appropriate antibiotics for all the experiments. Antibiotics were applied at final concentrations of: 50 μg mL^−1^ ampicillin (Sigma-Aldrich), 25 μg mL^−1^ kanamycin (Sigma-Aldrich), and 12.5 μg mL^−1^ chloramphenicol (Sigma-Aldrich).

Before gene expression assays unless otherwise mentioned, we co-transformed the desired circuits into chemically competent cells of *E. coli* MC1061Δ*pspF* using a general chemical transformation protocol. Then, we picked a single colony carrying test circuits from the agar plate and suspended the cells in 30 μL LB-Lennox medium with appropriate antibiotics. 5 μL cell suspension was added into 195 μL LB-Lennox medium with appropriate antibiotics in a transparent flat-bottom 96-well plate (CytoOne). To avoid abnormal growth at the edges of a plate, the outermost wells were not used (but were filled with 200 μL of medium). The plates were cultured at 37 °C, 1000 rpm on a plate shaker (MB100-4A) for 18 h to 23 h overnight. For induction, we added 2 μL overnight culture into 198 μL LB-Lennox medium with appropriate antibiotics and inducers on a flat clear bottom 96-well plate with black walls (Greiner Bio-one). The inducers employed in this study include (N-(3-oxohexanoyl)-L-homoserine lactone, AHL (Sigma-Aldrich), aTc (CAYMAN Chemical), rhamnose (Alfa Aesar), arabinose (Acros Organics). All stock inducer solutions have 40 × concentrations, which were diluted to 1 × final concentrations by adding 5 μL of stock solutions into a final volume of 200 μL culture system.

All reagents were purchased as powder and were prepared by dissolving in double distilled water or ethanol (chloramphenicol) followed by filtration through 0.22 μM syringe filters (Millipore).

### Plasmid circuit construction

We used standard molecular biology protocols in this study for circuit construction. All plasmids and their structures and sequences are listed in **Supplementary Table 3, 5-9**. The typical composition of the plasmids used in this study is illustrated in **Supplementary Figure 9**. Key primers and oligonucleotides used in this study are listed in **Supplementary Table 4**. The dCas9 generators / dxCas9 generators, activator generators, and reporter circuits were carried by the BioBrick vector pSB4A3 (*36*). The sgRNA or crRNA generators were hosted on a separate and compatible vector, p15AC. A third compatible vector, pSAVE221, carried the tracrRNA generators with kanamycin resistance (GenBank: JX560327). Variants of each genetic part were chosen according to the specific needs and details of a particular experiment. Qiaspin Miniprep Kit (Qiagen) and Monarch PCR & DNA Cleanup Kit (NEB) were used in this study for DNA purification.

All the genetic parts of the dCas9 generator, dxCas9 generator, activator generator, inducible promoters, and sgRNA with aptamers at the tetraloop come from our previous published study (*18*). The new versions of sgRNA were synthesized by annealing of oligonucleotides (j5 protocol (*37*)). PCR was used to split sgRNA to crRNA and tracrRNA, introduce mutations into gRNAs, and change the spacer sequence of crRNAs.

All the plasmids have been sequenced to confirm the sequence (Source Bioscience), except for the CONAN DNA, which cannot be sequenced because it contains a long reverse palindrome sequence, and it has been checked by HpyAV restriction mapping.

### *In vivo* gene expression assay

For the experiments hijacking mRNA as crRNA, after 7 h induction culture, the cultures on 96-well plate were read by a plate reader (BMG FLUOstar fluorometry) equipped with a 485 nm excitation laser and a 520 nm emission filter for green fluorescence measurements (Gain 700), and a 584 nm excitation laser and a 620–10 nm emission filter for red fluorescence measurements (Gain 2000). The *A_600_* data was collected at the same time for calculating fluorescence/*A_600_*.

For assays of **Figure 1c, 1d, 1f, 2c**, to improve accuracy, after 6 h induction culture, the cultures were fixed by adding 2 μL culture into 198 μL 1 × phosphate-buffered saline (K813-500ML, VWR) in a 96-well U-bottom plate (Thermo Fisher Scientific) with 1 mg mL^−1^ kanamycin and the samples were stored at 4 °C for at least 1 hr. The fixed samples on 96-well U-bottom plate were read by the Attune NxT Flow Cytometer (equipped with Attune NxT Autosampler), equipped with 488 nm excitation laser and 530/30 nm emission filter for green fluorescence measurements.

For other assays, after 6 h induction culture, the cultures on 96-well plate were read by a plate reader (BMG FLUOstar fluorometry) equipped with a 485 nm excitation laser and a 520 nm emission filter for green fluorescence measurements (Gain 700), the *A_600_* data was collected at the same time for calculating fluorescence/A_*600*_.

### *In vitro* CONAN assay

The *in vitro* CONAN assays were performed in a 10 μL reaction volume using a previously developed *in vitro* transcription only cell-free system (TXO) (*32*). Each reaction contained 5 ng μL^−1^ DNA, 0.75 × optimized transcription/detection buffer (OTDB) (10× stock solution includes 400 mM Tris base pH 7.5 adjusted with HCl (Sigma-Aldrich), 60 mM MgCl_2_·6H_2_O (Sigma-Aldrich), 100 mM DTT (Melford) and 20 mM Spermidine (Alfa Aesar), 1.5 mM NTPs (Thermo Scientific™ NTP Set, Tris buffered). 1.4 units μL^−1^ T7 RNA polymerase (RNAP, NEB 2 units μL^−1^ RNase inhibitor (NEB), 50 nM Cas9 (NEB), 50 nM synthesized tracrRNA (IDT, **Supplementary Table 2**), 50 nM synthesized SARS-CoV-2 RNA fragment (IDT, **Supplementary Table 2**), 0.5 units μL^−1^ PstI (NEB) and 7.5 μM DFHBI (Tocris). SARS-CoV-2 RNA was replaced with nuclease free water (NEB) for the assay without the target RNA. For the assay with pre-incubation, everything except T7 RNAP and CONAN DNA was mixed and first incubated at 37°C for 10 min for allow the formation of CRISPR/Cas9 complex, then the CONAN DNA was added and the mixture was incubated further at 37°C for 1 h to allow the cleavage of the DNA, after which the T7 RNAP was added to the mixture to activate the Broccoli transcription. The CONAN DNA was purified using QIAprep Spin Miniprep Kit (Qiagen) following the manufacturer’s protocol.

For each assay, 35 uL reaction were made, then three replicates with 10 uL of each reaction were loaded into a black 384-well microplate with clear bottom (Greiner Bio-One). The plate was sealed with a transparent EASYseal plate sealer (Greiner Bio-One) and was incubated and measured continuously by BMG FLUOstar plate reader for 8 h at 37 °C with 1s shaking (200 rpm, double orbital) before each measurement. The settings for measuring the fluorescence were the same as for the *in vivo* characterization. The plate reader data were processed using Omega MARS 3.20 R2, Microsoft Excel 2013 and GraphPad Prism v9.1.1. The background of output signals was subtracted from each *in vitro* reaction by using its triplicate-averaged counterpart of the negative control (reporter-free) at the same time. All the data shown are mean values with standard deviation as error bars.

To prepare for cell phone imaging at the end of CONAN assays, the microplate was placed onto the surface of a Safe Imager (S37102, Invitrogen) blue-light trans-illuminator and was covered with an amber filter in a dark environment. A cell phone (iPhone 11) was used to acquire the images with the build-in night mode and 3 s auto exposure. No additional adjustments were made to the images.

### Calculation of fluorescence intensities

All attempts at replication were successful. For data collected by flow cytometer, the geometric mean of fluorescence of a population was calculated in FlowJo v10.7.1, then corrected by the geometric mean of background fluorescence of cells with suitable empty plasmids grown in identical conditions but without inducers. Corrected fluorescence values were then averaged in GraphPad Prism v9.1.1 to give the means with standard deviations (s.d.), which were used for plotting graphs.

For data collected by the plate reader, the fluorescence values and OD values were corrected by a blank negative control (with corresponding volumes of medium and with water replacing inducers). Then, the fluorescence/*A_600_* for each well was then calculated. The fluorescence/A*600* value for each well was corrected by that of a fluorescence-free strain. All the above steps were done in Microsoft Excel 2016. Corrected fluorescence/*A_600_* values were then averaged in GraphPad Prism v9.1.1 to give the means with standard deviations (SD), which were used for plotting graphs.

For the dynamic range calculation, the mean and standard deviations (SD) of the corrected fluorescence values of replicates were calculated in Microsoft Excel 2016. We calculated the increase in fluorescence value *ΔFluo* (corrected fluorescence *Fluo_ON_* of the ON state minus the corrected fluorescence *Fluo_OFF_* of the OFF state) and the corresponding standard deviations (SD) *SD_ΔFluo_*. The dynamic range *DR* was calculated by dividing *ΔFluo* by *Fluo_OFF_*. The standard deviations *SD_DR_* of the dynamic range *DR* was calculated by the uncertainty propagation formula:

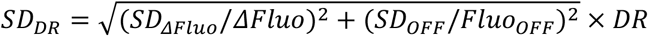

For the CONAN sensor, the cell-free system without adding any DNA template was employed as the blank sample, and the fluorescence values were corrected by the blank control.

### Characterization of orthogonality and the AND logic gates

For the characterization of orthogonality and AND logic gates (**Figure 1f, 1g**), we performed high-throughput co-transformation and subsequent cell culturing steps without plating the cells on agar plates. 1.3 μL of each plasmid (miniprep of 10 mL cell culture) was added into 96-well PCR plates, and each well included a particular combination of three plasmids. 25 μL chemically competent cells of *E. coli* MC1061*ΔpspF* was added into each well on ice, and then the 96-well PCR plates were kept on ice for 20 min. A 60 s heat shock was done by a PCR machine (ABI Veriti 96 well PCR Thermal Cycler) at 42°C, then the plates were placed back on ice for 5 min.

15 - 25 μL transformation product of each well was inoculated into a flat-bottom 96-well plate (CytoOne) with 150 μL fresh LB-Lennox medium. The plates were cultured at 37°C, 800 rpm on a plate shaker (MB100-4A) for 2 h to recover. Then, the recovered sample of each well was diluted 4-fold by adding 50 μL culture into 150 μL LB-Lennox medium with appropriate antibiotics in another flat-bottom 96-well plate (CytoOne) for overnight culture at 37°C, 1000 rpm on a plate shaker (MB100-4A). Then the samples were diluted 100-fold to final volume 200 μL for the second overnight culture at 37°C, 1000 rpm. The overnight culture was diluted for 6 h induction culture followed by plate reader reading.

### Library of crRNA-tracrRNA pairs with randomized hybridizing regions

Random sequences were generated by online tool (*38*), then the sense strand and antisense strand sequences were used for the paired segments of crRNA and tracrRNA, respectively. Among them, tracrRNA retains the bulge structure and the G for the wobble base pairs. We use Excel 2016 to check whether there is an NGG in crRNA next to the spacer and exclude these sequences. Sticky ends compatible with the expression vector were designed to ensure that the DNA annealing products can be directly used for ligation.

The randomized sequences for every reprogramed crRNA and tracrRNA were ordered as complementary DNA oligonucleotides (Merck), annealed and cloned into the respective backbone plasmids. The annealing was carried out in 96-well PCR plates by mixing 2 μL of each oligonucleotide at 100 μM with 2 μL of 10x T4 DNA Ligase Reaction Buffer (NEB) and 14 μL of water. The plate was then incubated at 95 °C for 5 min in a thermocycler and allowed to cool down within the machine until reaching room temperature (c.a. 20 min). The annealed oligonucleotides produce overhangs compatible with the digested destination plasmids. The crRNA duplexes were cloned into the plasmid pLY257, previously digested with BbsI-HF (NEB), and the tracrRNA duplexes were cloned into the plasmid pLY258, previously digested with BsaI-HFv2 (NEB). Backbone plasmids were purified from gel after digestion using the Monarch DNA Gel Extraction Kit (NEB). Ligations were carried out in 96-well PCR plates by mixing 0.2 μL of digested plasmid (3-9 ng μL^−1^) and 0.3 μL of annealed oligonucleotides with 0.5 μL of 2x Instant Sticky-end Ligase Master Mix (NEB) and immediately placing the plate on ice. 5 μL of chemically competent TOP10 cells were added to each well and the plate was incubated on ice for 30 min. The entire plate was heat shocked for 45s at 42 °C in a thermocycler and incubated on ice for 2 min before adding 194 μL of LB medium to the wells. The cells were transferred to a flat-bottom 96-well plate and incubated for 1 h at 37 °C and 1,000 rpm on a plate shaker (Allsheng). After this recovery time, 10 μL of cells were spotted on LB agar containing the appropriate antibiotics on an OneWell Plate (Greiner) and incubated overnight at 37 °C. Individual colonies were picked from the plates and inoculated into flat-bottom 96-well plate containing 200 μL of LB medium, grown for 5 h at 37 °C and 1,000 rpm on a plate shaker. 2 μL of cells were used to inoculate 96-deepwell plates (Starlab) containing 1.25 mL of TB medium and incubated overnight at 37 °C and 700 rpm on a plate shaker. Cells were collected by centrifuging the plates for 10 min at 1,200 x *g* and the plasmids’ DNA was purified using the QIAGEN Plasmid Plus 96 Miniprep Kit (Qiagen), following the manufacturers’ instructions. All plasmids were sequenced to confirm the sequence (Source Bioscience). Complementary plasmids were co-transformed into competent MC1061Δ*pspF* cells carrying the reporter plasmid pLY54 (prepared using the Mix & Go *E. coli* Transformation Kit from Zymo Research, following the manufacturer’s instructions). Briefly, 2 μL of each plasmid were added to 22.5 μL of the MC1061Δ*pspF* competent cells in a 96-well PCR plate, incubated on ice for 30 min and the entire plate was heat shocked for 45s at 42 °C in a thermocycler. After additional incubation on ice for 2 min, 175 μL of SOC medium was added to the wells, transferred to a flat-bottom 96-well plate and incubated for 1 h at 37 °C and 1,000 rpm on a plate shaker. After recovery, 30 μL were transferred to a new plate containing 170 μL of SOC medium supplemented with the appropriate antibiotics (ampicillin, kanamycin and tetracycline) and incubated overnight at 37 °C and 1,000 rpm on a plate shaker. For each experiment performed on three different days, 2 μL of the cultures were used to inoculate 198 μL of fresh LB medium supplemented with the appropriate antibiotics and grown overnight at 37 °C and 1,000 rpm on a plate shaker. After overnight growth, 2 μL of cultures were used to inoculate 198 μL of fresh LB medium supplemented with the appropriate antibiotics and 2.5 ng mL^−1^ aTc to induce dCas9 expression, on a 96-well μclear black flat-bottom plate covered with a lid (GBO) to prevent evaporation. The plate was incubated inside a FLUOstar Omega microplate reader (BMG labtech) at 37 °C and 700 rpm, for a period of 23 h. End-point fluorescence for GFP (excitation filter: 485 nm; emission filter: 520–10 nm; gain = 800) and culture optical density at 600 nm (OD600) were measured every 20 min using the Omega Control v5.11 (BMG Labtech) and data was analysed using Omega MARS Software v3.32 (BMG Labtech). The LB medium background fluorescence and absorbance were subtracted from the readings of sample wells and fluorescence/OD600 (Fluo./OD600) was calculated for all samples. Normalized data were produced by subtracting the negative control Fluo./OD600 from the other samples and the 6 h data was exported to GraphPad Prism v9.1.1 for plotting the figures.

The variables considered were: the average Fluo./OD600 calculated for the three biological replicates at 6 h after inoculation; the minimum free energy (MFE) of the crRNA optimal secondary structure, predicted using the RNAfold web server (http://rna.tbi.univie.ac.at/cgi-bin/RNAWebSuite/RNAfold.cgi) (Variable 1); the ΔG for crRNA and tracrRNA matching region heterodimer binding, predicted using the RNAcofold web server (http://rna.tbi.univie.ac.at/cgi-bin/RNAWebSuite/RNAcofold.cgi) (Variable 2); the homology between the crRNA, without the terminator sequence, and the target DNA sequence (35 nucleotides including the spacer region and the 15 nucleotides downstream) (Variable 3); the alignment scores of the crRNA and tracrRNA matching region calculated using CLUSTALW (https://www.genome.jp/tools-bin/clustalw) (Variable 4) and the GC content of the crRNA and tracrRNA matching region (Variable 5).

The Pearson correlation analysis was conducted with the R package “ggcorrplot”, and the regression model was built with the MATLAB 2015b function “regress”.

## Data availability

All data in the main text and the supplementary materials are available upon reasonable request.

## Acknowledgements

We thank Dr Jennifer Tullet (University of Kent) for supplying the W3110 and HT115 *E. coli* strains used in this study. This work was supported by the UK Research and Innovation Future Leaders Fellowship [MR/S018875/1], Leverhulme Trust grant [RPG-2020-241] and US Office of Naval Research Global grant [N62909-20-1-2036]. Z.X. was supported by the Natural Science Foundation of China (31771483, 61721003).

## Contributions

Y.L. and B.W. conceived the study and designed the experiments. Y.L. performed the majority of the experiments. F.P. performed the experiments related to the library of crRNA-tracrRNA pairs with randomized matching regions. X.W. and M.L. performed the experiments for the CONAN sensor. Y.L. F.P. and X.W. performed data analysis. S.P. and Z.X. analysed the data from the random paired sequence library. C.F. provided support and materials for the development of the *in vitro* RNA sensor. All authors took part in the interpretation of results and preparation of materials for the manuscript. B.W. and Y.L. wrote the manuscript. B.W. supervised and acquired the funding of the study.

## Competing Interests

The authors declare no competing interests.

## Notes

### Competing Interest Statement

The authors have declared no competing interest.

