## Supplementary Information for "Reprogrammed tracrRNAs enable repurposing RNAs as crRNAs and detecting RNAs"

**Supplementary Figures 1−8**

**Supplementary Tables 1−9**

**References 1−2**

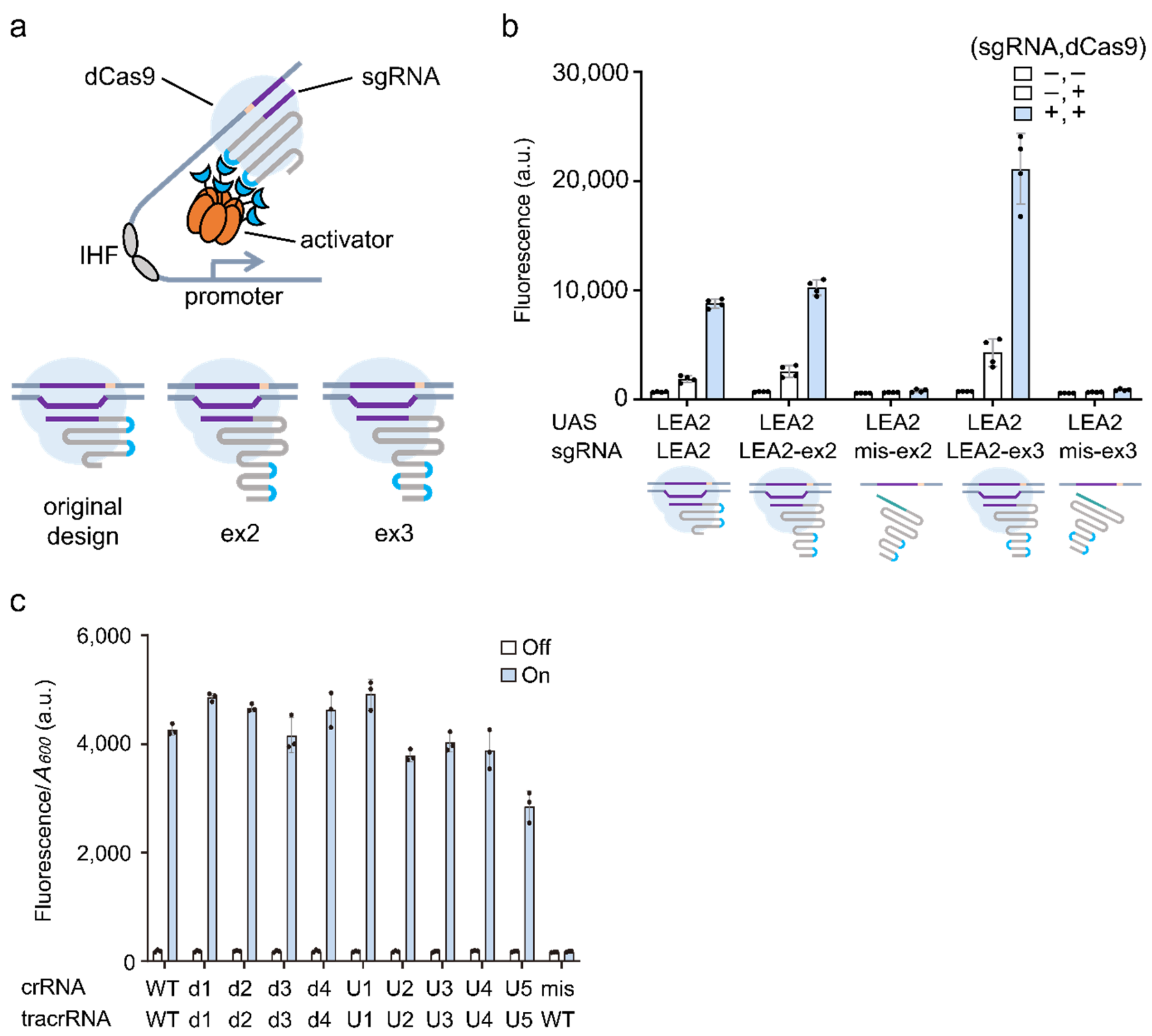


**Supplementary Figure 1. CRISPRa function with tail-fused aptamers and CRISPRa output from reprogrammed crRNA-tracrRNA pairs (a)** Schematic of the CRISPRa devices showing three different designs of sgRNA. The light blue U shape line segments in the sgRNA scaffold indicate BoxB RNA aptamers. **(b)** CRISPRa function with three different kinds of sgRNA design. The random sequence LEA2 was used and UAS and sgRNA spacer. For each of the two new designs, a sgRNA with the same sgRNA scaffold but with different spacer (random sequence LEB3) was employed as mismatched control. dCas9 expression was controlled by $P_{tet}$ promoter, and activator expression was driven by a constitutive promoter J23106 (Anderson promoter). A $P_{lux2}$ promoter was used to transcribe sgRNA. 2.5 ng $mL^{-1}$ aTc and 1.6 μM AHL were used for dCas9 and sgRNA induction respectively. Error bars, s.d. ($n$ = 4). **(c)** The CRISPRa function of the various crRNA-tracrRNA pairs in the library shown in **Figure 1e**. The label 'mis' means the WT crRNA has a mismatched spacer (LEB3) with the target UAS (LEA2). All the other crRNAs in this test have spacer LEA2, and a corresponding $\sigma^{54}$-dependent promoter with UAS LEA2 was used for the reporter. Expression of dCas9 and activator expression was driven by the $P_{tet}$ and constitutive promoter J23106. A $P_{lux2}$ promoter was used to transcribe sgRNA. No inducer was added for the OFF state. 2.5 ng $mL^{-1}$ aTc, 1.6 μM AHL, and 0.08 mM arabinose were used for dCas9, crRNA, tracrRNA induction (ON state), respectively. Error bars, s.d. ($n$ = 3); a.u., arbitrary units.

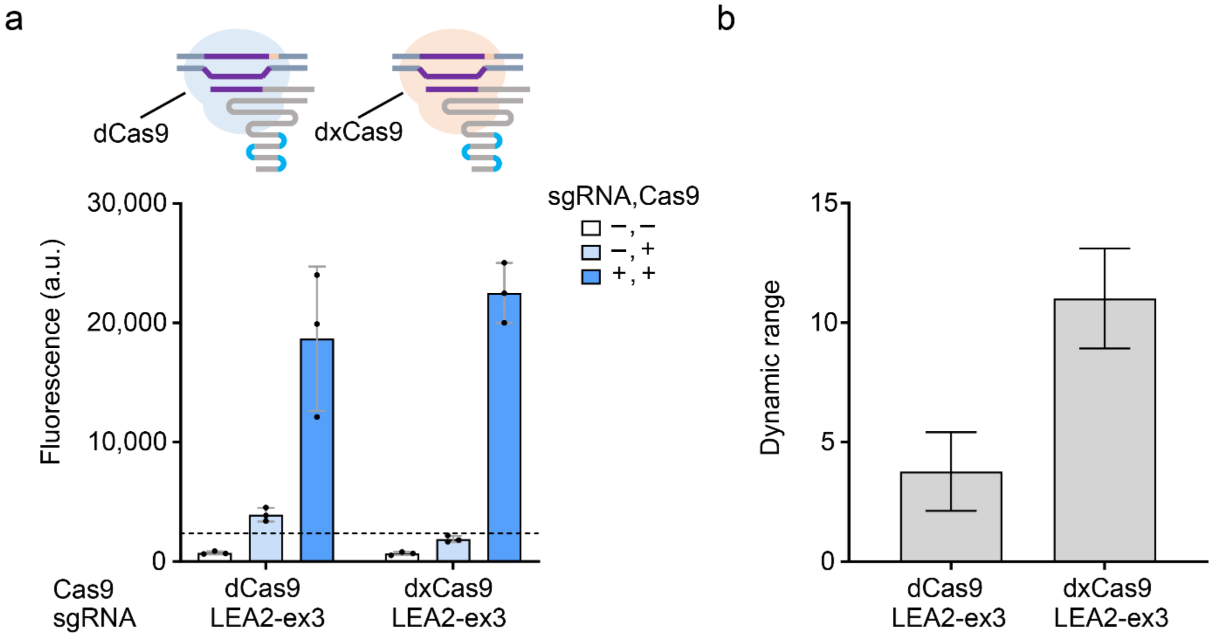


**Supplementary Figure 2. sgRNA tuning-based output dynamic range can be improved by employing dxCas9 instead of dCas9. (a)** A sgRNA with three BoxB aptamers added to its 3' end tail (LEA2-ex3) was used for CRISPRa. The UAS of promoter and spacer of sgRNA had the LEA2 sequence. $P_{tet}$ controlled dCas9 or dxCas9 generator, and $P_{rhaB}$ drove activator expression, $P_{lux2}$ drove sgRNA transcription. Inducer concentrations: 2.5 ng mL$^{-1}$ aTc, 0.4 mM rhamnose, and 1.6 μM AHL. The plus and minus signs represent the presence or absence of inducers, respectively. **(b)** The sgRNA tuning based dynamic range was calculated from the two sets of data with dCas9 or dxCas9 in the left bar chart. Error bars, s.d. (*n* = 3); a.u., arbitrary units.

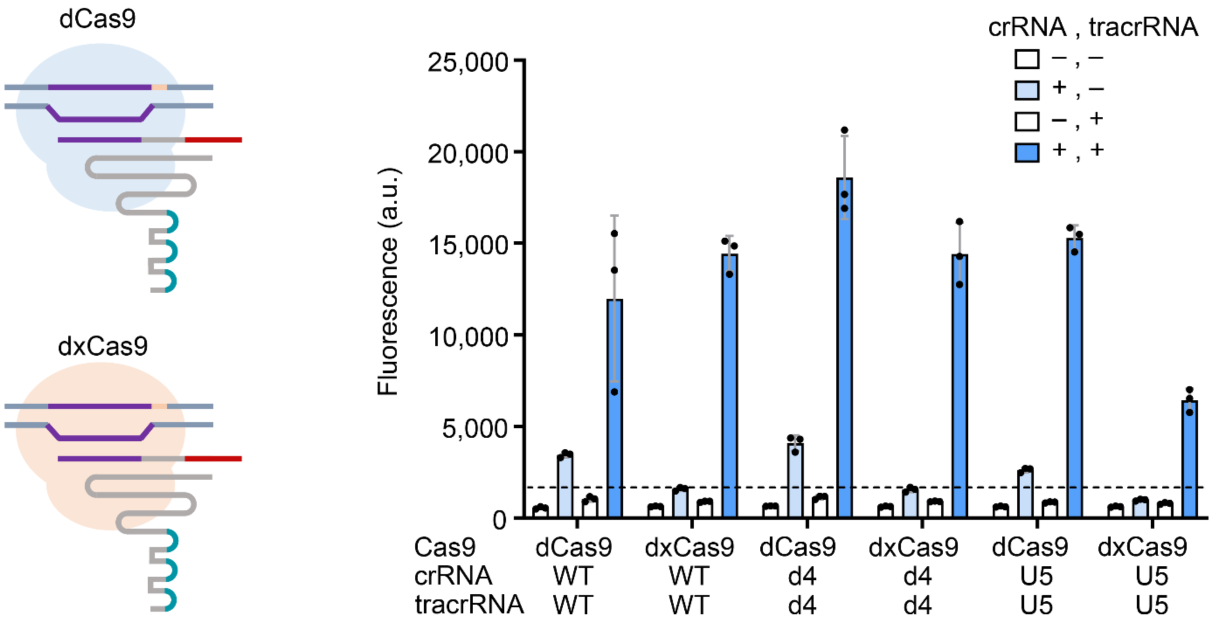


**Supplementary Figure 3. Optimizing asymmetry sensitivity to inputs of crRNA-tracrRNA mediated CRISPRa by utilizing dxCas9.** The cartoon on the left shows the two CRISPR complexes used in the experiment. The purple line segment represents DNA and its complementary paired spacer sequence of crRNA. The red line indicates the terminator sequence on crRNA. The blue U shaped segment indicates BoxB aptamers at the tail of tracrRNA. The bar chart on the right shows the output of three CRISPRa devices with different crRNA-tracrRNA matching sequences (WT, d4, U5). Random sequence LEA2 was used for UAS and spacer in these circuits. Expression of dCas9 or dxCas9 was driven by $P_{tet}$, and expression of activator was driven by $P_{rhaB}$. crRNA and tracrRNA transcription was driven by $P_{lux2}$ and $P_{BAD,}$ respectively. Inducer concentrations: 2.5 ng $mL^{-1}$ aTc, 0.4 mM rhamnose, 1.6 μM AHL, and 0.08 mM arabinose. The plus and minus signs represent the presence or absence of inducers. Error bars, s.d. (*n* = 3); a.u., arbitrary units.

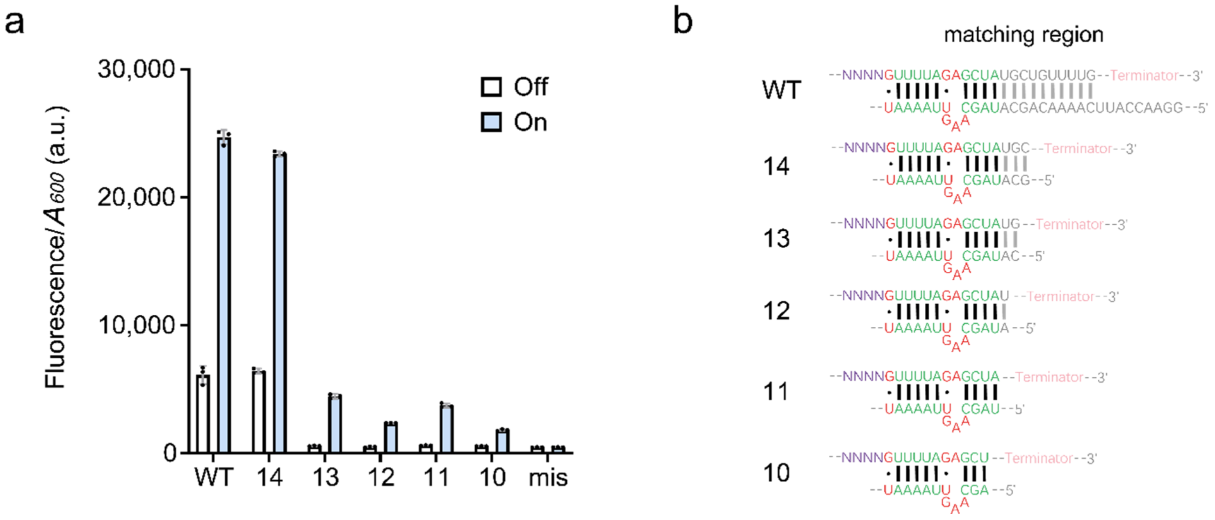


**Supplementary Figure 4. Minimal length of the hybridizing sequences required for functional crRNA-tracrRNA pair. (a)** CRISPRa output from different lengths of the hybridizing region. The bar chart shows the results from crRNA-tracrRNA pairs with 10 –14 bp length of the matching region. The dCas9 was expressed by $P_{tet}$ promoter with 2.5 ng $mL^{-1}$ aTc. The activator, crRNA and tracrRNA are all driven by the constitutive promoter J23106. The label ‘mis’ means the WT crRNA has a mismatched spacer (LEB3) with the target UAS (LEA2). All the other crRNAs in this test have spacer LEA2, and a corresponding $\sigma^{54}$-dependent promoter with UAS LEA2 was used for the reporter. The data of crRNA-tracrRNA pairs WT, 14 –12 bp, and mis is equal to that in **Figure 2a**. Error bars, s.d. ($n$ = 3); a.u., arbitrary units. **(b)** The structure and sequences of the crRNA-tracrRNA pairs we employed in this experiment.

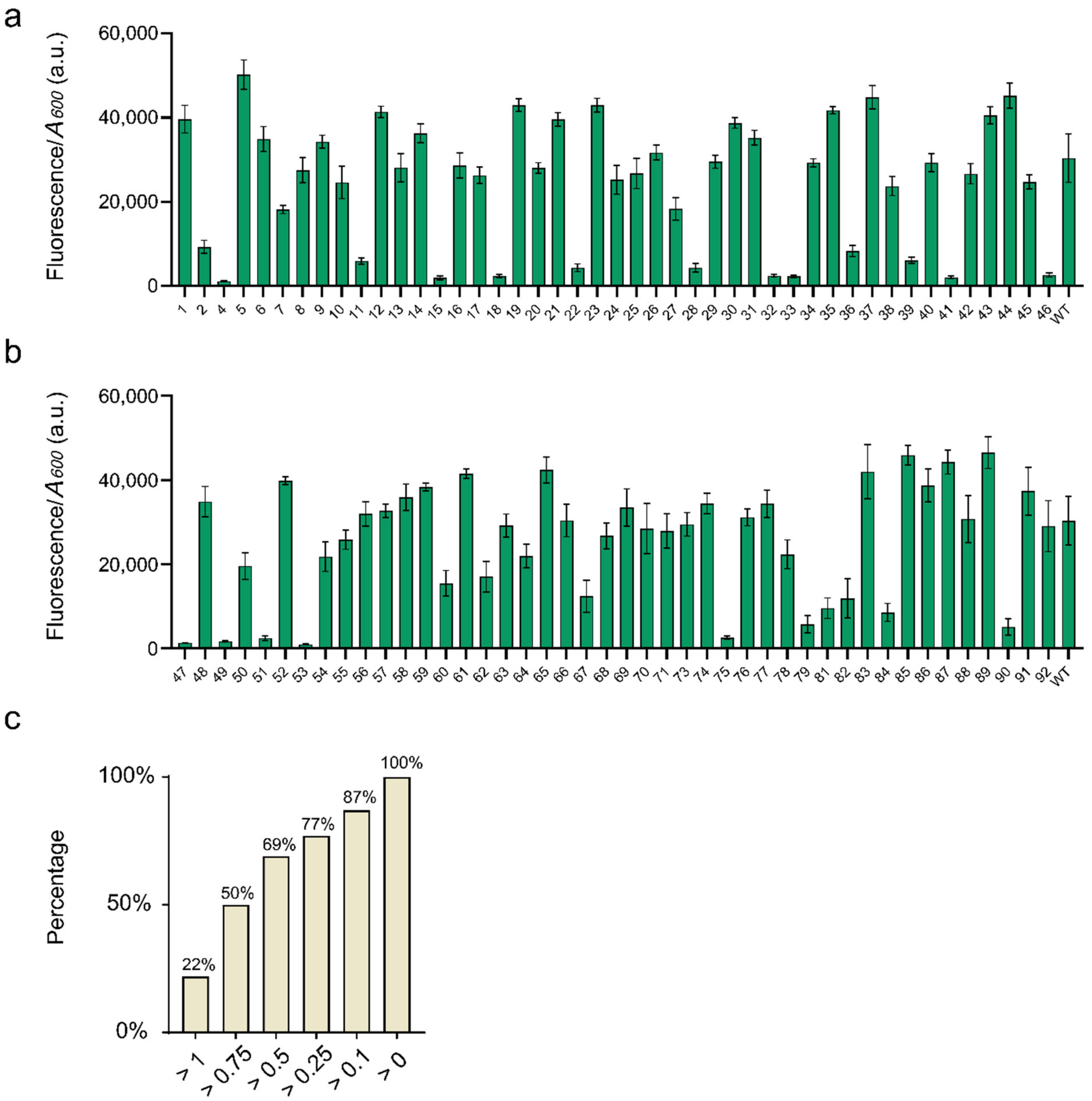


**Supplementary Figure 5. CRISPRa output from the paired crRNA-tracrRNA randomized-sequence library. (a)** The CRISPRa output from strains 1 – 46 and a positive control strain with the WT crRNA-tracrRNA matching region sequence. The dCas9 generator was controlled by $P_{tet}$. The crRNA and tracrRNA transcription and activator expression was driven by Promoter J23106. 2.5 ng $mL^{-1}$ aTc was used in this test. Error bars, s.d. (n = 3); a.u., arbitrary units. **(b)** The CRISPRa output from strains 47 – 92 and a positive control strain with the WT crRNA-tracrRNA matching region sequence. The induction condition is the same as in **a**. Error bars, s.d. ($n$ = 3); a.u., arbitrary units. **(c)** The percentage of the candidates whose output level is higher than a certain value. The abscissa defines the output range for each group, the values are the different multipliers for output from the positive control strain. The corresponding output lower limit is the product of these multipliers and the output value from the positive control strain.

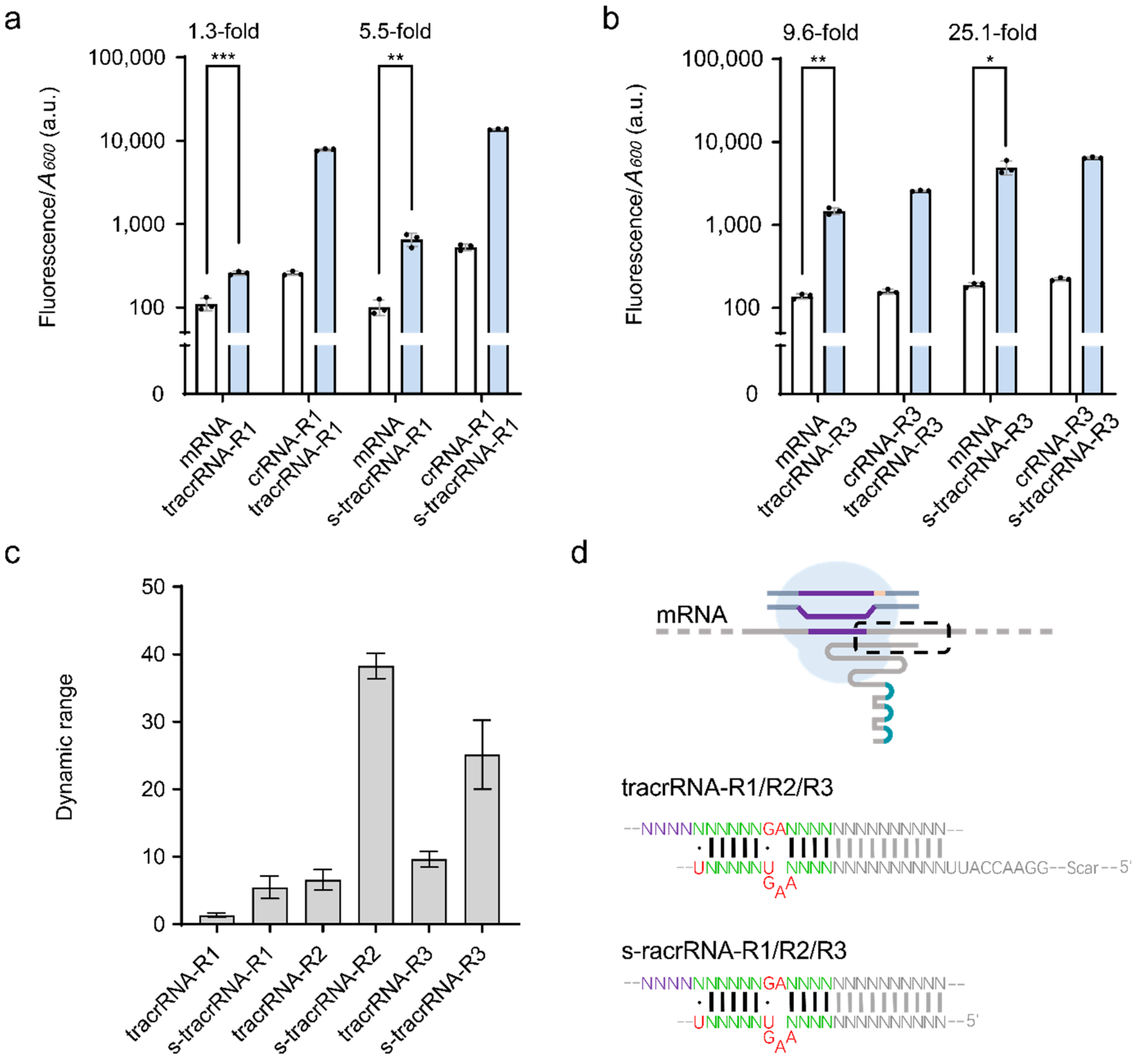


**Supplementary Figure 6. Engineered tracrRNA repurposes mRNA into crRNA. (a)** 5’ end truncation of tracrRNA improves the mRNA-mediated CRISPR function. The mRNA fragment with the same sequences as that in the mRNA target site R1 was expressed as controls. For the bar of mRNA + tracrRNA-R1, the data is the same as in **Figure 3b**. Expression of dCas9 and activator was driven by the $P_{tet}$ and $P_{rhaB}$ promoters, respectively. The mRNA of RFP and the tracrRNA were transcribed from $P_{lux2}$ and $P_{BAD,}$ respectively. Inducer concentrations: 2.5 ng $mL^{-1}$ aTc, 0.2 mM rhamnose, and 0.08 mM arabinose. 0.1 μM AHL was used for mRNA expression. Statistical difference was determined by a two-tailed *t* test: mRNA + s-tracrRNA-R1, $p = 0.0014$, t = 7.904. Error bars, s.d. ($n = 3$) **(b)** The content is the same as in **a**, except that the RNA target is R3. Statistical difference was determined by a Welch’s *t* test: mRNA + s-tracrRNA-R3, $p = 0.0121$, t = 8.994. Error bars, s.d. ($n = 3$). **(c)** The dynamic range of different mRNA-mediated CRISPRa devices. All the induction condition is the same as in **a**. Error bars, s.d. (n=3). **(d)** Schematic diagram shows the structures of the hybridizing regions for the original tracrRNA version and the 5' end truncated tracrRNA. a.u., arbitrary units.

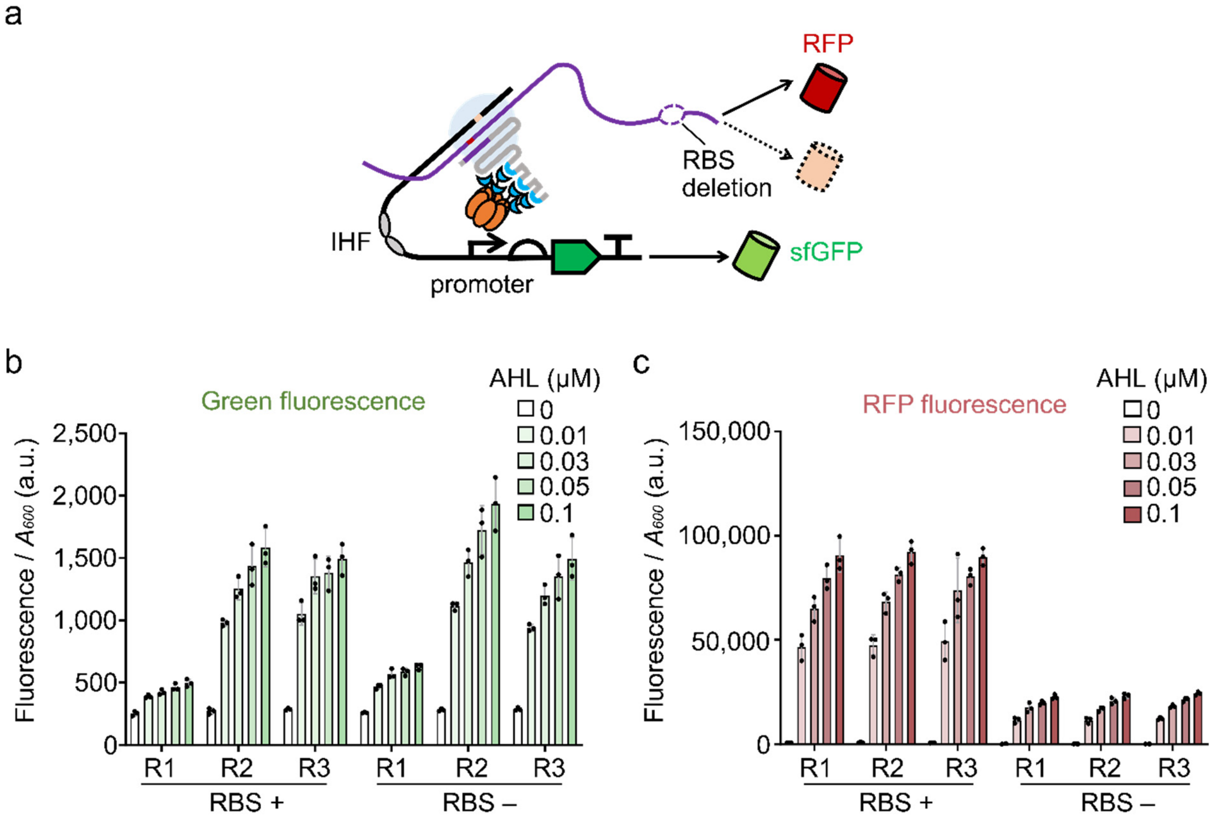


**Supplementary Figure 7. Test of ribosome binding site (RBS) deletion in the RFP mRNA hijacking experiment. (a)** Schematic showing the circuit design for the RBS deletion experiment. The oval purple dotted line represents the RBS, and the red cylindrical structure drawn by the dotted line indicates that the translation of red fluorescent protein is greatly reduced after the RBS is deleted. **(b)** The green fluorescence output from the mRNA mediated CRISPRa with or without an RBS on the mRNA of RFP. Expression of dCas9 and activator was driven by $P_{tet}$ and $P_{rhaB}$ promoters. mRNA and tracrRNA transcription were driven by $P_{lux2}$ and $P_{BAD}$, respectively. Inducer concentrations: 2.5 ng mL$^{-1}$ aTc, 0.2 mM rhamnose, and 0.08 mM arabinose. AHL gradian (0, 0.01, 0.03, 0.05, 0.1 μM) was used for mRNA induction. **(c)** The red fluorescence output from the mRNA mediated CRISPRa with or without an RBS on the mRNA of RFP. The data was from the same experiment in **b**. Error bars, s.d. (*n* = 3). a.u., arbitrary units.

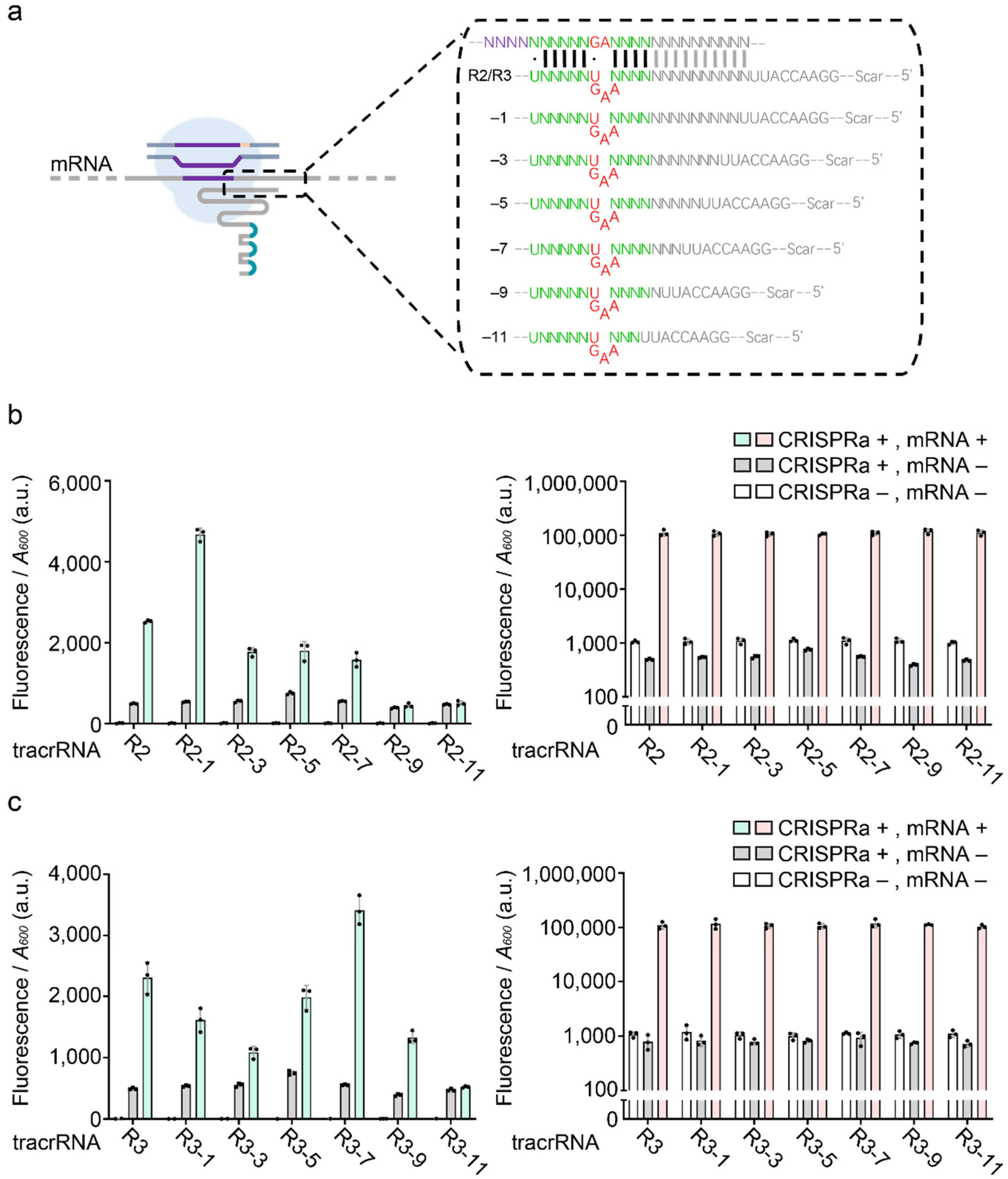


**Supplementary Figure 8. Length of the crRNA-tracrRNA hybridizing region affects the CRISPRa function. (a)** Schematic showing the design of the library, including truncated tracrRNA anti-coding regions. In this test, tracrRNA-R2 and tracrRNA-R3 were employed and coupled with mRNA of RFP for CRISPRa. The initial pairing length is 21 bp. **(b)** The result of the anti-coding region truncation test on tracrRNA-R2. Green fluorescence detection results on the left and red fluorescence detection results on the right. Expression of dCas9 generator and activator were driven by $P_{tet}$ and $P_{rhaB}$. mRNA and tracrRNA transcription were driven by $P_{lux2}$ and $P_{BAD}$, respectively. Inducer concentrations: 2.5 ng mL$^{-1}$ aTc, 1.6 μM AHL, 0.4 mM rhamnose, and 0.08 mM arabinose. CRISPR + or CRISPR – indicates that all the inducers were added or not except AHL (for mRNA induction). **(c)** The result of the anti-coding region truncation test on tracrRNA-R3. Green fluorescence detection results are on the left, and red fluorescence detection results are on the right. All the conditions were the same as that in **b.** Error bars, s.d. (*n* = 3); a.u., arbitrary units.

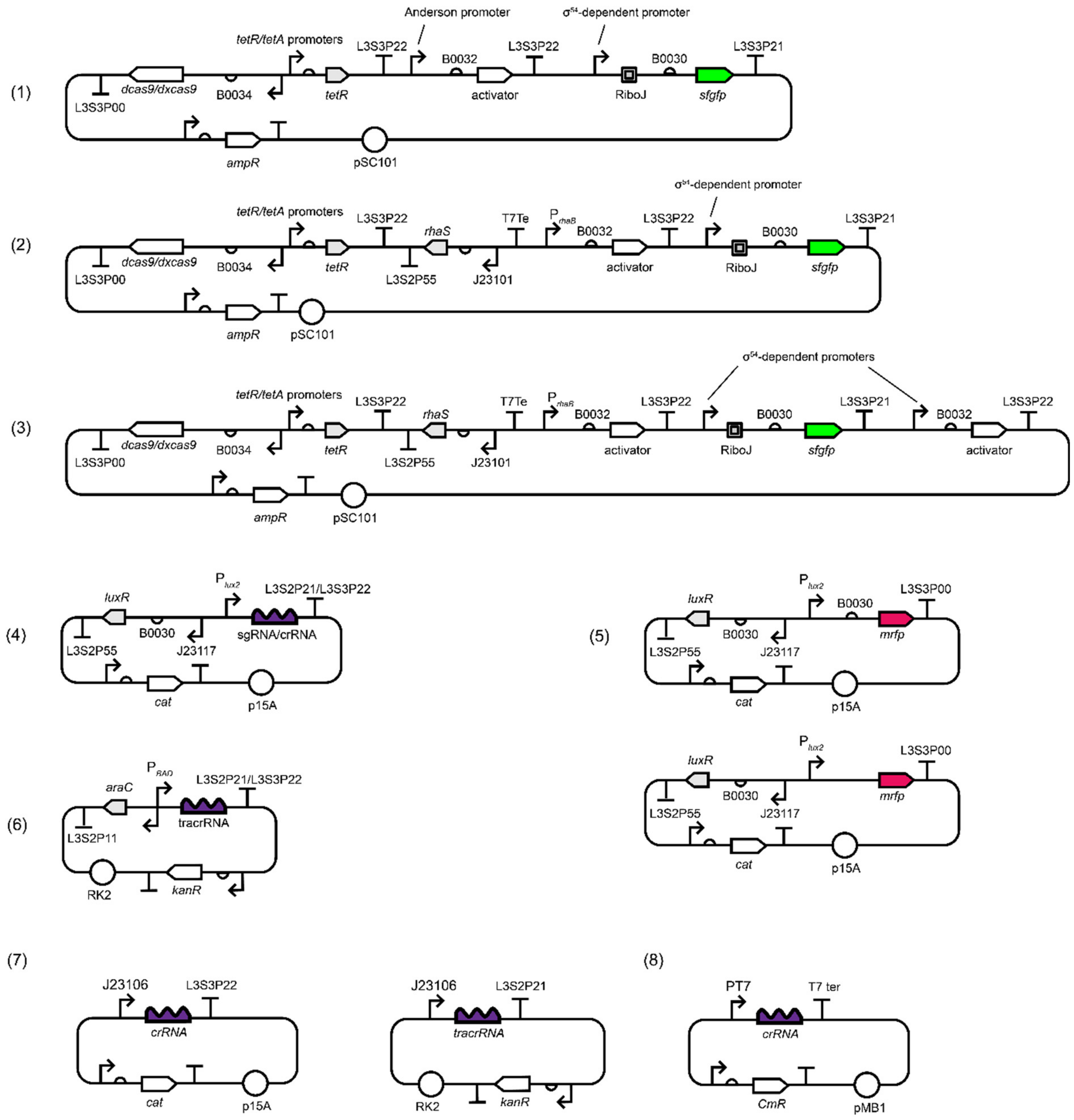


**Supplementary Figure 9. Representative plasmid maps for key circuit constructs used in this study. (1)** The plasmid map of pLY54. pLY54 was used in the experiments in **Figure 1,2**, and also was used in tests of **Supplementary Figure 1−5**. **(2)** The plasmid map of pLY162−pLY164, and pLY167-pLY169, pLY255, pLY256. pLY162−pLY164 were used in the RFP mRNA hijacking experiment (**Figure 3**, **Supplementary Figure 6,7,8**); pLY167, pLY168, pLY255, pLY256 are reporter circuits for the hijacking of endogenous mRNA of the arsenic-related gene cluster in *E. coli* (**Figure 4**); pLY169 is the CRISPRa circuit with dxCas9, which was used in experiments of **Supplementary Figure 2,3**. **(3)** The plasmid map of pLY165 and pLY166, they are positive feedback circuits used in **Figure 3**. **(4)** The plasmid map of pLY76, pLY170−pLY184, pLY187−pLY189, pLY218−pLY231 and pLY246. They are all the crRNA generators in this study. **(5)** The plasmid map of pLY185 and pLY186. They are two mRFP generators with or without RBS, which were used in **Supplementary Figure 7**. **(6)** The plasmid map of pLY190−pLY217, pLY232−pLY239, pLY251−pLY254. They are all the tracrRNA generators in this study. **(7)** The plasmid map of pLY241−pLY250, and the 180 crRNA/tracrRNA generators in our library **(Supplementary Table 1)**. **(8)** The plasmid map of pIY240, which is a CONAN circuit **(Figure 5)**.

### Supplementary tables

**Supplementary Table 1: Sequences from the library of randomized hybridizing region of the paired crRNA-tracrRNA**

| Name | Sequence in hybridized region (crRNA 5' – 3') | Sequence in hybridized region (tracrRNA 5' – 3') |
|---|---|---|
| c(+) | GUUUUAGAGCUAUGC | GCAUAGCAAGUUAAAAU |
| c1 | AGCAGGGAACAGACU | AGUCUGUAAGUCCUGCU |
| c2 | GAUCGGUAAAAGUUA | UAACUUUUAAGUCCGAUU |
| c4 | AAAUGUAUUUAAAGA | UCUUUAAAAGUACAUUU |
| c5 | AUUCACAUGAAUCCU | AGGAUUCAAGUGUGAAU |
| c6 | UACGUACCGGGUGAA | UUCACCCAAGUUACGUU |
| c7 | CAGGCUCGGAUGUCC | GGACAUCAAGUAGCCUU |
| c8 | UUCAACCAGCCCUUG | CAAGGGCAAGUGUUGAU |
| c9 | AGAAUCCUCGCUCAG | CUGAGCGAAGUGAUUCU |
| c10 | GCCUGGAACACCUAU | AUAGGUGAAGUCCAGGU |
| c11 | CCCUGAUACCCUGUU | AACAGGGAAGUUCAGGU |
| c12 | GUUCGCGCACGCGCG | CGCGCGUAAGUGCGAAU |
| c13 | CACCUCCGAGCGCUG | CAGCGCUAAGUGAGGUU |
| c14 | GGUCUCACCCUUCAA | UUGAAGGAAGUGAGACU |
| c15 | AAAGAAAUUUCUGAU | AUCAGAAAAGUUUCUUU |
| c16 | UCGCGUUUUGUAACU | AGUUACAAAGUACGCGU |
| c17 | GACUCAUGGGUAUUU | AAAUACCAAGUUGAGUU |
| c18 | CAAAUGUCGUAUUUG | CAAAUACAAGUCAUUUU |
| c19 | GGCCUCGAAUCCAUG | CAUGGAUAAGUGAGGCU |
| c20 | GACCCAUAUGGCGGG | CCCGCCAAAGUUGGGUU |
| c21 | CCAGUGUCUCCCAAC | GUUGGGAAAGUCACUGU |
| c22 | UCGAGAUGACUUUCU | AGAAAGUAAGUUCUCGU |
| c23 | GGAGAAAUAGUUGCG | CGCAACUAAGUUUCUCU |
| c24 | CCAUAGAUGAUCAUU | AAUGAUCAAGUCUAUGU |
| c25 | UCCAACAAUCCACAC | GUGUGGAAAGUGUUGGU |

| Name | Sequence in hybridized region (crRNA 5' – 3') | Sequence in hybridized region (tracrRNA 5' – 3') |
|---|---|---|
| c26 | ACUAGUUUCCCCAAG | CUUGGGGAAGUACUAGU |
| c27 | CGACAAAAGAGUUAA | UUAACUCAAGUUUGUCU |
| c28 | GCUGGGGCAGCCCUA | UAGGGCUAAGUCCCAGU |
| c29 | CGAGAUGUGUUGGUU | AACCAACAAGUAUCUCU |
| c30 | CCAGUAAGACACAGC | GCUGUGUAAGUUACUGU |
| c31 | ACGGUCCACUCUUUU | AAAAGAGAAGUGACCGU |
| c32 | GCGCUAAAAUAGCUG | CAGCUAUAAGUUAGCGU |
| c33 | CGAUUUGAGGCUGAG | CUCAGCCAAGUAAAUCU |
| c34 | GUGUUAAAAUCCUUG | CAAGGAUAAGUUAACAU |
| c35 | AAAUUGGUCUUAACC | GGUUAAGAAGUCAAUUU |
| c36 | UAGGAGCUUGACAGA | UCUGUCAAAGUCUCCUU |
| c37 | CUGCCGUGUGUCAGU | ACUGACAAAGUCGGCAU |
| c38 | UUCAGGCGCCUCUGC | GCAGAGGAAGUCCUGAU |
| c39 | AGCCGGACCACCGAG | CUCGGUGAAGUCCGGCU |
| c40 | UUGGAAGUCCUCCGG | CCGGAGGAAGUUUCCAU |
| c41 | GUCUAAUCGACCUAG | CUAGGUCAAGUUUAGAU |
| c42 | UUCGGGUUCAGCGCA | UGCGCUGAAGUCCCGAU |
| c43 | UACUCCUUUACGAAA | UUUCGUAAAGUGGAGUU |
| c44 | CAUUGAGCCGUUGCC | GGCAACGAAGUUCAAUU |
| c45 | ACCUGGUGCUGAUCA | UGAUCAGAAGUCCAGGU |
| c46 | UAGACCCAGGCCGGC | GCCGGCCAAGUGGUCUU |
| c47 | UACUUUGGAAAUAUA | UAUAUUUAAGUAAAGUU |
| c48 | AUUAUUGCCGACACA | UGUGUCGAAGUAAUAAU |
| c49 | GCUGUUUAGAGGCUG | CAGCCUCAAGUAACAGU |
| c50 | CGAGUGUCGACCCUG | CAGGGUCAAGUCACUCU |
| c51 | GUAAUGAUCCGUAAU | AUUACGGAAGUCAUUAU |
| c52 | CUUAUCCGGCAACGC | GCGUUGCAAGUGAUAAU |
| c53 | UCCGCGACCGGCGGG | CCCGCCGAAGUCGCGGU |
| c54 | UCGAACGAGCAGAUG | CAUCUGCAAGUGUUCGU |

| Name | Sequence in hybridized region (crRNA 5' – 3') | Sequence in hybridized region (tracrRNA 5' – 3') |
|---|---|---|
| c55 | AGUCAAGACGGAUAA | UUAUCCGAAGUUUGACU |
| c56 | UGUAGGUCACGGAGU | ACUCCGUAAGUCCUACU |
| c57 | AGUACGAUGAGCGGG | CCCGCUCAAGUCGUACU |
| c58 | UAUCUCUAGGAGGGU | ACCCUCCAAGUGAGAUU |
| c59 | CAUAUCCCGGUCCCA | UGGGACCAAGUGAUAUU |
| c60 | AUCCGAGCGGCGGCC | GGCCGCCAAGUUCGGAU |
| c61 | GUCCGUAUACUCCUG | CAGGAGUAAGUACGGAU |
| c62 | CGUCUGCACCGUUGA | UCAACGGAAGUCAGACU |
| c63 | UUGACGGCAUGUAGG | CCUACAUAAGUCGUCAU |
| c64 | UCUAGGCACCCCUAC | GUAGGGGAAGUCCUAGU |
| c65 | GCAUAUCCCUUAAGC | GCUUAAGAAGUAUAUGU |
| c66 | UUAUACAGGGAGUAC | GUACUCCAAGUGUAUAU |
| c67 | CCCACCCGAAUCAUC | GAUGAUUAAGUGGUGGU |
| c68 | ACCCUGCACGACGUU | AACGUCGAAGUCAGGGU |
| c69 | AAGGUCUGACGCCGG | CCGGCGUAAGUGACCUU |
| c70 | UAUGAGACACACGCA | UGCGUGUAAGUCUCAUU |
| c71 | CCUUGGCGUCUGCUU | AAGCAGAAAGUCCAAGU |
| c73 | CAUGAGGUCGAUAUC | GAUAUCGAAGUCUCAUU |
| c74 | AAUCGGGAGCUGCCU | AGGCAGCAAGUCCGAUU |
| c75 | ACUUUGGUACAGUGA | UCACUGUAAGUCAAAGU |
| c76 | GUCAGUCUGCUGCGC | GCGCAGCAAGUACUGAU |
| c77 | AUAGAUGUCCUGCGG | CCGCAGGAAGUAUCUAU |
| c78 | GAAUCACAAGAGGCG | CGCCUCUAAGUUGAUUU |
| c79 | AACCAUAGAAACAUU | AAUGUUUAAGUAUGGUU |
| c81 | UCUCAGAGAACCUGA | UCAGGUUAAGUCUGAGU |
| c82 | GUUAACCCCUUCUAA | UUAGAAGAAGUGUUAAU |
| c83 | UUUCACGAUAGUCUU | AAGACUAAAGUGUGAAU |
| c84 | GCACACCCGGUCGAA | UUCGACCAAGUGUGUGU |
| c85 | AAGUGAGAUUCCGGU | ACCGGAAAAGUUCACUU |

| Name | Sequence in hybridized region (crRNA 5' – 3') | Sequence in hybridized region (tracrRNA 5' – 3') |
|---|---|---|
| c86 | GACCGAUCAGCCCUG | CAGGGCUAAGUUCGGUU |
| c87 | CAGUGUUUGCAUCGC | GCGAUGCAAGUACACUU |
| c88 | CGCAGAUCCAUGUAC | GUACAUGAAGUUCUGCU |
| c89 | GGUAGGAAGGCAUCC | GGAUGCCAAGUCCUACU |
| c90 | CCGUCGCGAGCCUAU | AUAGGCUAAGUCGACGU |
| c91 | GCAUUCGAGUGGCAA | UUGCCACAAGUGAAUGU |
| c92 | UAGUCAACUAGGGGU | ACCCCUAAAGUUGACUU |

**Supplementary Table 2: Synthetic RNAs used in CONAN sensor assays**

| Name | Sequence in hybridized region |
|---|---|
| SARS-CoV-II RNA 1 | GCCACUUCUGCUGCUCUUCAACCUGAAGAAGAGCAAGAAGAAGAUUGG |
| tracrRNA CoV1 | GGAACCAUUAAUCUUCUUCUUGCAAGUUUCUUUAAGGCUAGUCCGUUAUCAACUUG AAAAAGUGGCACCGAGUCGGUGCUUUUUUU |
| SARS-CoV-II RNA 2 | CAACUUGUAUGAUGUGUUACAAACGUAAUAGAGCAACAAGAGUCGAAUGU |
| tracrRNA CoV2 | GGAACCAUUUCGACUCUUGUUGCAAGUUAUUAUAAGGCUAGUCCGUUAUCAACUUG AAAAAGUGGCACCGAGUCGGUGCUUUUUUU |

**Supplementary Table 3: Plasmids used in this study**

'dCas9' in this table refers 'dCas9 generator'

'dxCas9' in this table refers 'dxCas9 generator'

'Reporter' in this table refers 'Reporter with sfGFP::ASV tag'

Notes:

- Scars between the parts coming from construction are not shown.
- Circuits in the randomized sequence library are not included in this table.

| Name | Structure | Vector | Reference |
|---|---|---|---|
| pLY54 | dCas9-J23106-PSPFΔHTH::λN22plus-$P_{pspA}$-LEA2B2-Reporter | pSB4A3* | Previous study(*1*) |
| pLY162 | dCas9-$P_{rhaB}$-PSPFΔHTH::λN22plus-$P_{pspA}$-R1-Reporter | pSB4A3* | This study |
| pLY163 | dCas9-$P_{rhaB}$-PSPFΔHTH::λN22plus-$P_{pspA}$-R2-Reporter | pSB4A3* | This study |
| pLY164 | dCas9-$P_{rhaB}$-PSPFΔHTH::λN22plus-$P_{pspA}$-R3-Reporter | pSB4A3* | This study |
| pLY165 | dCas9-$P_{rhaB}$-PSPFΔHTH::λN22plus-$P_{pspA}$-R2-Reporter-$P_{pspA}$-R2-Reporter | pSB4A3* | This study |
| pLY166 | dCas9-$P_{rhaB}$-PSPFΔHTH::λN22plus-$P_{pspA}$-R3-Reporter-$P_{pspA}$-R3-Reporter | pSB4A3* | This study |
| pLY167 | dCas9-$P_{rhaB}$-PSPFΔHTH::λN22plus-$P_{pspA}$-Ar1-Reporter | pSB4A3* | This study |
| pLY168 | dCas9-$P_{rhaB}$-PSPFΔHTH::λN22plus-$P_{pspA}$-Ar2-Reporter | pSB4A3* | This study |
| pLY169 | dxCas9-$P_{rhaB}$-PSPFΔHTH::λN22plus-$P_{pspA}$-LEA2B2-Reporter | pSB4A3* | This study |
| pLY76 | $P_{lux2}$-sgRNA-LEA2 | p15AC | Previous study(*1*) |
| pLY170 | $P_{lux2}$-sgRNA-LEA2-ex2 | p15AC | This study |
| pLY171 | $P_{lux2}$-sgRNA-LEA2-ex3 | p15AC | This study |
| pLY172 | $P_{lux2}$-sgRNA-LEB3-ex2 | p15AC | This study |
| pLY173 | $P_{lux2}$-sgRNA-LEB3-ex3 | p15AC | This study |
| pLY174 | $P_{lux2}$-crRNA-LEA2-WT | p15AC | This study |
| pLY176 | $P_{lux2}$-crRNA-LEA2-d1 | p15AC | This study |

| Name | Structure | Vector | Reference |
|---|---|---|---|
| pLY177 | $P_{lux2}$-crRNA-LEA2-d2 | p15AC | This study |
| pLY178 | $P_{lux2}$-crRNA-LEA2-d3 | p15AC | This study |
| pLY179 | $P_{lux2}$-crRNA-LEA2-d4 | p15AC | This study |
| pLY180 | $P_{lux2}$-crRNA-LEA2-U1 | p15AC | This study |
| pLY181 | $P_{lux2}$-crRNA-LEA2-U2 | p15AC | This study |
| pLY182 | $P_{lux2}$-crRNA-LEA2-U3 | p15AC | This study |
| pLY183 | $P_{lux2}$-crRNA-LEA2-U4 | p15AC | This study |
| pLY184 | $P_{lux2}$-crRNA-LEA2-U5 | p15AC | This study |
| pLY185 | $P_{lux2}$-mRFP | p15AC | This study |
| pLY186 | $P_{lux2}$-mRFP-ΔRBS | p15AC | This study |
| pLY187 | $P_{lux2}$-crRNA-LEA2-U5A | p15AC | This study |
| pLY188 | $P_{lux2}$-crRNA-LEA2-U5C | p15AC | This study |
| pLY189 | $P_{lux2}$-crRNA-LEA2-U5T | p15AC | This study |
| pLY190 | $P_{BAD}$-tracrRNA-WT | pSEVA221 (JX560327)(*2*) | This study |
| pLY192 | $P_{BAD}$-tracrRNA-d1 | pSEVA221 (JX560327) | This study |
| pLY193 | $P_{BAD}$-tracrRNA-d2 | pSEVA221 (JX560327) | This study |
| pLY194 | $P_{BAD}$-tracrRNA-d3 | pSEVA221 (JX560327) | This study |
| pLY195 | $P_{BAD}$-tracrRNA-d4 | pSEVA221 (JX560327) | This study |
| pLY196 | $P_{BAD}$-tracrRNA-U1 | pSEVA221 (JX560327) | This study |
| pLY197 | $P_{BAD}$-tracrRNA-U2 | pSEVA221 (JX560327) | This study |
| pLY198 | $P_{BAD}$-tracrRNA-U3 | pSEVA221 (JX560327) | This study |
| pLY199 | $P_{BAD}$-tracrRNA-U4 | pSEVA221 | This study |

| Name | Structure | Vector | Reference |
|---|---|---|---|
| | | (JX560327) | |
| pLY200 | $P_{BAD}$-tracrRNA-U5 | pSEVA221 (JX560327) | This study |
| pLY201 | $P_{BAD}$-tracrRNA-R1 | pSEVA221 (JX560327) | This study |
| pLY202 | $P_{BAD}$-tracrRNA-R2 | pSEVA221 (JX560327) | This study |
| pLY203 | $P_{BAD}$-tracrRNA-R3 | pSEVA221 (JX560327) | This study |
| pLY204 | $P_{BAD}$-tracrRNA-Ar1 | pSEVA221 (JX560327) | This study |
| pLY205 | $P_{BAD}$-tracrRNA-Ar2 | pSEVA221 (JX560327) | This study |
| pLY206 | $P_{BAD}$-tracrRNA-R2L1 | pSEVA221 (JX560327) | This study |
| pLY207 | $P_{BAD}$-tracrRNA-R2L3 | pSEVA221 (JX560327) | This study |
| pLY208 | $P_{BAD}$-tracrRNA-R2L5 | pSEVA221 (JX560327) | This study |
| pLY209 | $P_{BAD}$-tracrRNA-R2L7 | pSEVA221 (JX560327) | This study |
| pLY210 | $P_{BAD}$-tracrRNA-R2L9 | pSEVA221 (JX560327) | This study |
| pLY211 | $P_{BAD}$-tracrRNA-R2L11 | pSEVA221 (JX560327) | This study |
| pLY212 | $P_{BAD}$-tracrRNA-R3L1 | pSEVA221 (JX560327) | This study |
| pLY213 | $P_{BAD}$-tracrRNA-R3L3 | pSEVA221 (JX560327) | This study |
| pLY214 | $P_{BAD}$-tracrRNA-R3L5 | pSEVA221 | This study |

| Name | Structure | Vector | Reference |
|---|---|---|---|
| | | (JX560327) | |
| pLY215 | $P_{BAD}$-tracrRNA-R3L7 | pSEVA221 (JX560327) | This study |
| pLY216 | $P_{BAD}$-tracrRNA-R3L9 | pSEVA221 (JX560327) | This study |
| pLY217 | $P_{BAD}$-tracrRNA-R3L11 | pSEVA221 (JX560327) | This study |
| pLY218 | $P_{lux2}$-crRNA-R1 | p15AC | This study |
| pLY219 | $P_{lux2}$-crRNA-R2 | p15AC | This study |
| pLY220 | $P_{lux2}$-crRNA-R3 | p15AC | This study |
| pLY221 | $P_{lux2}$-mRNA-Ar | p15AC | This study |
| pLY222 | $P_{lux2}$-crRNA-Ar1 | p15AC | This study |
| pLY223 | $P_{lux2}$-crRNA-Ar2 | p15AC | This study |
| pLY224 | $P_{lux2}$-crRNA-Ar3 | p15AC | This study |
| pLY225 | $P_{lux2}$-crRNA-Ar4 | p15AC | This study |
| pLY226 | $P_{lux2}$-s-crRNA-LEB3 | p15AC | This study |
| pLY227 | $P_{lux2}$-s-crRNA-LEA2-WT | p15AC | This study |
| pLY228 | $P_{lux2}$-s-crRNA-LEA2-C6 | p15AC | This study |
| pLY229 | $P_{lux2}$-s-crRNA-LEA2-C9 | p15AC | This study |
| pLY230 | $P_{lux2}$-s-crRNA-LEA2-C48 | p15AC | This study |
| pLY231 | $P_{lux2}$-s-crRNA-LEA2-C74 | p15AC | This study |
| pLY232 | $P_{BAD}$-s-tracrRNA- WT | pSEVA221 (JX560327) | This study |
| pLY233 | $P_{BAD}$-s-tracrRNA- C6 | pSEVA221 (JX560327) | This study |
| pLY234 | $P_{BAD}$-s-tracrRNA- C9 | pSEVA221 (JX560327) | This study |
| pLY235 | $P_{BAD}$-s-tracrRNA- C48 | pSEVA221 (JX560327) | This study |
| pLY236 | $P_{BAD}$-s-tracrRNA- C74 | pSEVA221 | This study |

| Name | Structure | Vector | Reference |
|---|---|---|---|
| | | (JX560327) | |
| pLY237 | $P_{BAD}$-s-tracrRNA- R1 | pSEVA221 (JX560327) | This study |
| pLY238 | $P_{BAD}$-s-tracrRNA- R2 | pSEVA221 (JX560327) | This study |
| pLY239 | $P_{BAD}$-s-tracrRNA- R3 | pSEVA221 (JX560327) | This study |
| pLY240 | $P_{T7}$-CONAN | pSB1C3** | This study |
| pLY241 | J23106-s-crRNA-13 | p15AC | This study |
| pLY242 | J23106-s-crRNA-12 | p15AC | This study |
| pLY243 | J23106-s-crRNA-11 | p15AC | This study |
| pLY244 | J23106-s-crRNA-10 | p15AC | This study |
| pLY247 | J23106-s-tracrRNA-13 | pSEVA221 (JX560327) | This study |
| pLY248 | J23106-s-tracrRNA-12 | pSEVA221 (JX560327) | This study |
| pLY249 | J23106-s-tracrRNA-11 | pSEVA221 (JX560327) | This study |
| pLY250 | J23106-s-tracrRNA-10 | pSEVA221 (JX560327) | This study |
| pLY251 | $P_{BAD}$-s-tracrRNA- Ar2 | pSEVA221 (JX560327) | This study |
| pLY245 | $P_{BAD}$-tracrRNA-ESI | pSEVA221 (JX560327) | This study |
| pLY246 | $P_{lux2}$-crRNA-LEB3-WT | p15AC | This study |
| pLY252 | $P_{BAD}$-tracrRNA-Ar3 | pSEVA221 (JX560327) | This study |
| pLY253 | $P_{BAD}$-tracrRNA-Ar4 | pSEVA221 (JX560327) | This study |
| pLY254 | $P_{BAD}$-tracrRNA-WT-21T | pSEVA221 | This study |

| Name | Structure | Vector | Reference |
|---|---|---|---|
| | | (JX560327) | |
| pLY255 | dCas9- $P_{rhaB}$-PSPFΔHTH::λN22plus-$P_{pspA}$-Ar3-Reporter | pSB4A3* | This study |
| pLY256 | dCas9- $P_{rhaB}$-PSPFΔHTH::λN22plus-$P_{pspA}$-Ar4-Reporter | pSB4A3* | This study |
| pLY257 | J23106-crRNA-ESI | p15AC | This study |
| pLY258 | J23106-tracrRNA-ESI | pSEVA221 (JX560327) | This study |

*http://parts.igem.org/Part:pSB4A3 (There are a few point mutations)

**http://parts.igem.org/Part:pSB1C3

#### Supplementary Table 4: Key primers and oligos used in this study

| Name | Sequence | Purpose |
|---|---|---|
| 23-gRNA-F1 | CTAGAAGTTATTATATAGTTCGGTCGTTTGAGAGCTAGAAATAGCAAGTTCAAATAAGGCTAGTCCGTTATCAACTTGAAAAAGTGGCACCGAGTCG | For sgRNA-LEA2-ex2/ex3 synthesis |
| 23-gRNA-R1 | GCACCGACTCGGTGCCACTTTTTCAAGTTGATAACGGACTAGCCTTATTTGAACTTGCTATTTCTAGCTCTCAAACGACCGAACTATATAATAACTT | For sgRNA-LEA2-ex2/ex3 synthesis |
| exBoxB-F2 | GTGCGGGCCCTGAAGAAGGGCCCTAGCAAGTTCAAATAAGGCTAGTCCGTTAT | For sgRNA-LEA2-ex2 synthesis |
| exBoxB-R2 | GTTGATAACGGACTAGCCTTATTTGAACTTGCTAGGGCCCTTCTTCAGGGCCC | For sgRNA-LEA2-ex2 synthesis |
| exBoxB-R3 | GCGGCCGCTACTAGTAAAAAAAAGCACCGACTCGGTGCCACTTGGGCCCTTCTTCAGGGCCCAA | For sgRNA-LEA2-ex2 synthesis |
| exBoxB-F3 | CAACTTGGGCCCTGAAGAAGGGCCCAAGTGGCACCGAGTCGGTGCTTTTTTTTACTAGTAGCGGCCGCTGCA | For sgRNA-LEA2-ex2 synthesis |
| ex3BoxB-F2 | GTGCGGGCCCTGAAGAAGGGCCCAAGGCTAGGGCCCTGAAGAAGGGCCCGCAGGGCCCTGAAGAAGGGCCCTTTTTTTTACTAGTAGCGGCCGCTGCA | For sgRNA-LEA2-ex3 synthesis |
| ex3BoxB-R2 | GCGGCCGCTACTAGTAAAAAAAAGGGCCCTTCTTCAGGGCCCTGCGGGCCCTTCTTCAGGGCCCTAGCCTTGGGCCCTTCTTCAGGGCCC | For sgRNA-LEA2-ex3 synthesis |
| R2L1-F | CTAGAGGAACCATTCTCGGAGGAGGCCAAGTAGCGTTAAGGCTAGTC | For tracrRNA-R2L1 synthesis |
| R2L1-R | AACGGACTAGCCTTAACGCTACTTGGCCTCCTCCGAGAATGGTTCCT | For tracrRNA-R2L1 synthesis |
| R2L3-F | CTAGAGGAACCATTCGGAGGAGGCCAAGTAGCGTTAAGGCTAGTC | For tracrRNA-R2L3 synthesis |
| R2L3-R | AACGGACTAGCCTTAACGCTACTTGGCCTCCTCCGAATGGTTCCT | For tracrRNA-R2L3 synthesis |
| R2L5-F | CTAGAGGAACCATTGAGGAGGCCAAGTAGCGTTAAGGCTAGTC | For tracrRNA-R2L5 synthesis |
| R2L5-R | AACGGACTAGCCTTAACGCTACTTGGCCTCCTCAATGGTTCCT | For tracrRNA-R2L5 synthesis |
| R2L7-F | CTAGAGGAACCATTGGAGGCCAAGTAGCGTTAAGGCTAGTC | For tracrRNA-R2L7 synthesis |
| R2L7-R | AACGGACTAGCCTTAACGCTACTTGGCCTCCAATGGTTCCT | For tracrRNA-R2L7 synthesis |
| R2L9-F | CTAGAGGAACCATTAGGCCAAGTAGCGTTAAGGCTAGTC | For tracrRNA-R2L9 synthesis |
| R2L9-R | AACGGACTAGCCTTAACGCTACTTGGCCTAATGGTTCCT | For tracrRNA-R2L9 synthesis |
| R2L11-F | CTAGAGGAACCATTGCCAAGTAGCGTTAAGGCTAGTC | For tracrRNA-R2L11 synthesis |
| R2L11-R | AACGGACTAGCCTTAACGCTACTTGGCAATGGTTCCT | For tracrRNA-R2L11 synthesis |
| R0L1-F | CTAGAGGAACCATTCTCGGAGGAGGCCAAGTCCAGCTAAGGCTAGTC | For tracrRNA-R3L1 synthesis |
| R0L1-R | AACGGACTAGCCTTAGCTGGACTTGGCCTCCTCCGAGAATGGTTCCT | For tracrRNA-R3L1 synthesis |
| R0L3-F | CTAGAGGAACCATTCGGAGGAGGCCAAGTCCAGCTAAGGCTAGTC | For tracrRNA-R3L3 synthesis |
| R0L3-R | AACGGACTAGCCTTAGCTGGACTTGGCCTCCTCCGAATGGTTCCT | For tracrRNA-R3L3 synthesis |
| R0L5-F | CTAGAGGAACCATTGAGGAGGCCAAGTCCAGCTAAGGCTAGTC | For tracrRNA-R3L5 synthesis |
| R0L5-R | AACGGACTAGCCTTAGCTGGACTTGGCCTCCTCAATGGTTCCT | For tracrRNA-R3L5 synthesis |
| R0L7-F | CTAGAGGAACCATTGGAGGCCAAGTCCAGCTAAGGCTAGTC | For tracrRNA-R3L7 synthesis |
| R0L7-R | AACGGACTAGCCTTAGCTGGACTTGGCCTCCAATGGTTCCT | For tracrRNA-R3L7 synthesis |
| R0L9-F | CTAGAGGAACCATTAGGCCAAGTCCAGCTAAGGCTAGTC | For tracrRNA-R3L9 synthesis |
| R0L9-R | AACGGACTAGCCTTAGCTGGACTTGGCCTAATGGTTCCT | For tracrRNA-R3L9 synthesis |
| R0L11-F | CTAGAGGAACCATTGCCAAGTCCAGCTAAGGCTAGTC | For tracrRNA-R3L11 synthesis |
| R0L11-R | AACGGACTAGCCTTAGCTGGACTTGGCAATGGTTCCT | For tracrRNA-R3L11 synthesis |
| Ar1F | CTAGAGGAACCATTAGCTGCAATGGCGCAAGTACCGGTAAGGCTAGTC | For tracrRNA-Ar1 synthesis |
| Ar1R | AACGGACTAGCCTTACCGGTACTTGCGCCATTGCAGCTAATGGTTCCT | For tracrRNA-Ar1 synthesis |
| Ar2F | CTAGAGGAACCATTAATGGTAATGTTGCAAGTATATCTAAGGCTAGTC | For tracrRNA-Ar2 synthesis |
| Ar2R | AACGGACTAGCCTTAGATATACTTGCAACATTACCATTAATGGTTCCT | For tracrRNA-Ar2 synthesis |

| Name | Sequence | Purpose |
|---|---|---|
| SpeI-wt-u5d3T-cr-R | CGCTACTAGTACAAAACAGCAATCGTCATTTTAGACCGAACTATATAATAACTTCTAGTTTTATTCG | For crRNA-LEA2-U5T synthesis |
| SpeI-wt-u5d3C-cr-R | CGCTACTAGTACAAAACAGCAATCGTCATTTTGGACCGAACTATATAATAACTTCTAGTTTTATTCG | For crRNA-LEA2-U5C synthesis |
| SpeI-wt-u5d3A-cr-R | CGCTACTAGTACAAAACAGCAATCGTCATTTTTGACCGAACTATATAATAACTTCTAGTTTTATTCG | For crRNA-LEA2-U5A synthesis |
| XbaI-wt-tracr-F | CGCTTCTAGAGGAACCATTCAAAACAGCATAGCAAGTTAAAATAAGGCTAGTCCGTTATCAACTTG | For tracrRNA-WT synthesis |
| SpeI-wt-cr-R | CGCTACTAGTACAAAACAGCATAGCTCTAAAACGACCGAACTATATAATAACTTCTAGTTTTATTCG | For crRNA-LEA2-WT synthesis |
| XbaI-wt-d1-tracr-F | CGCTTCTAGAGGAACCATTCAAAACAGCAAAGCAAGTTAAAATAAGGCTAGTCCGTTATCAACTTG | For tracrRNA-d1 synthesis |
| SpeI-wt-d1-cr-R | CGCTACTAGTACAAAACAGCAAAGCTCTAAAACGACCGAACTATATAATAACTTCTAGTTTTATTCG | For crRNA-LEA2-d1 synthesis |
| XbaI-wt-d2-tracr-F | CGCTTCTAGAGGAACCATTCAAAACAGCAATGCAAGTTAAAATAAGGCTAGTCCGTTATCAACTTG | For tracrRNA-d2 synthesis |
| SpeI-wt-d2-cr-R | CGCTACTAGTACAAAACAGCAATGCTCTAAAACGACCGAACTATATAATAACTTCTAGTTTTATTCG | For crRNA-LEA2-d2 synthesis |
| SpeI-wt-d3-cr-R | CGCTACTAGTACAAAACAGCAATCCTCTAAAACGACCGAACTATATAATAACTTCTAGTTTTATTCG | For crRNA-LEA2-d3 synthesis |
| XbaI-wt-d3-tracr-F | CGCTTCTAGAGGAACCATTCAAAACAGCAATCCAAGTTAAAATAAGGCTAGTCCGTTATCAACTTG | For tracrRNA-d3 synthesis |
| SpeI-wt-d4-cr-R | CGCTACTAGTACAAAACAGCAATCGTCTAAAACGACCGAACTATATAATAACTTCTAGTTTTATTCG | For crRNA-LEA2-d4 synthesis |
| XbaI-wt-d4-tracr-F | CGCTTCTAGAGGAACCATTCAAAACAGCAATCGAAGTTAAAATAAGGCTAGTCCGTTATCAACTTG | For tracrRNA-d4 synthesis |
| SpeI-wt-d4U1-cr-R | CGCTACTAGTACAAAACAGCAATCGTCAAAAACGACCGAACTATATAATAACTTCTAGTTTTATTCG | For crRNA-LEA2-U1 synthesis |
| XbaI-wt-d4U1-tracr-F | CGCTTCTAGAGGAACCATTCAAAACAGCAATCGAAGTAAAAATAAGGCTAGTCCGTTATCAACTTG | For tracrRNA-U1 synthesis |
| SpeI-wt-d4U2-cr-R | CGCTACTAGTACAAAACAGCAATCGTCATAAACGACCGAACTATATAATAACTTCTAGTTTTATTCG | For crRNA-LEA2-U2 synthesis |
| XbaI-wt-d4U2-tracr-F | CGCTTCTAGAGGAACCATTCAAAACAGCAATCGAAGTATAAATAAGGCTAGTCCGTTATCAACTTG | For tracrRNA-U2 synthesis |
| SpeI-wt-d4U3-cr-R | CGCTACTAGTACAAAACAGCAATCGTCATTAACGACCGAACTATATAATAACTTCTAGTTTTATTCG | For crRNA-LEA2-U3 synthesis |
| XbaI-wt-d4U3-tracr-F | CGCTTCTAGAGGAACCATTCAAAACAGCAATCGAAGTATTAATAAGGCTAGTCCGTTATCAACTTG | For tracrRNA-U3 synthesis |
| SpeI-wt-d4U4-cr-R | CGCTACTAGTACAAAACAGCAATCGTCATTTACGACCGAACTATATAATAACTTCTAGTTTTATTCG | For crRNA-LEA2-U4 synthesis |
| XbaI-wt-d4U4-tracr-F | CGCTTCTAGAGGAACCATTCAAAACAGCAATCGAAGTATTTATAAGGCTAGTCCGTTATCAACTTG | For tracrRNA-U4 synthesis |
| SpeI-wt-d4U5-cr-R | CGCTACTAGTACAAAACAGCAATCGTCATTTTCGACCGAACTATATAATAACTTCTAGTTTTATTCG | For crRNA-LEA2-U5 synthesis |
| XbaI-wt-d4U5-tracr-F | CGCTTCTAGAGGAACCATTCAAAACAGCAATCGAAGTATTTTTAAGGCTAGTCCGTTATCAACTTG | For tracrRNA-U5 synthesis |
| XbaI-R1-PspA | CGCTTCTAGAGCCCAGGACTCCTCACTTCAGCGGTTAGTGTAATTCGCTAACTCATCCTGGC | For $P_{pspA}$-R1 synthesis |
| XbaI-R2-PspA | CGCTTCTAGAGTAATGCAGAAGAAGACCATGCGGTTAGTGTAATTCGCTAACTCATCCTGGC | For $P_{pspA}$-R2 synthesis |
| XbaI-R3-PspA | CGCTTCTAGAGTGAAGGACGGCGGCCACTACCGGTTAGTGTAATTCGCTAACTCATCCTGGC | For $P_{pspA}$-R3 synthesis |
| XbaI-R1-tra | CGCTTCTAGAGGAACCATTCCTTGTAGATGAACAAGTGCCGTTAAGGCTAGTCCGTTATCAACTTG | For tracrRNA-R1 synthesis |
| XbaI-R2-tra | CGCTTCTAGAGGAACCATTGCTCGGAGGAGGCCAAGTCCAGCTAAGGCTAGTCCGTTATCAACTTG | For tracrRNA-R2 synthesis |
| XbaI-R3-tra | CGCTTCTAGAGGAACCATTAGGTGGTCTTGACCAAGTAGCGTTAAGGCTAGTCCGTTATCAACTTG | For tracrRNA-R3 synthesis |
| XP- L3S3P22-F | CTAGAGCCAATTATTGAAGGCCGCTAACGCGGCCTTTTTTTGTTTCTGGTCTCCCTACTAGTAGCGGCCGCTGCA | For terminator L3S3P22 synthesis |
| XP- L3S3P22-R | GCGGCCGCTACTAGTAGGGAGACCAGAAACAAAAAAAGGCCGCGTTAGCGGCCTTCAATAATTGGCT | For terminator L3S3P22 synthesis |
| XP-L3S2P21-F | CTAGAGCTCGGTACCAAATTCCAGAAAAGAGGCCTCCCGAAAGGGGGGCCTTTTTTCGTTTTGGTCCTACTAGTAGCGGCCGCTGCA | For terminator L3S2P21 synthesis |
| XP-L3S2P21-R | GCGGCCGCTACTAGTAGGACCAAAACGAAAAAAGGCCCCCCTTTCGGGAGGCCTCTTTTCTGGAATTTGGTACCGAGCT | For terminator L3S2P21 synthesis |
| PS1 (2) | AGGGCGGCGGATTTGTCC | For sequencing |
| PS2 (2) | GCGGCAACCGAGCGTTC | For sequencing |

| Name | Sequence | Purpose |
|---|---|---|
| VF2* | TGCCACCTGACGTCTAAGAA | For sequencing |
| VR** | ATTACCGCCTTTGAGTGAGC | For sequencing |
| CF2 | GTTCGTAAGCCATTTCCGCTCGCCGCAGTC | For sequencing |
| CR | AACGGTCTGGTTATAGGTACATTGAGCAAC | For sequencing |

* http://parts.igem.org/wiki/index.php?title=Part:BBa_G00100

** http://parts.igem.org/wiki/index.php?title=Part:BBa_G00101

#### Supplementary Table 5: Sequences for inducible promoters used in this study

| Name | Type and source | Parts | DNA sequence (5'– 3') |
|---|---|---|---|
| $P_{BAD}$ | Inducible promoter<br><br>Composite Module<br>(This study) | Reverse Terminator L3S2P11 | CCTAGTTGTCTTCATGCATGAAGACAAAATTAATACTAGAGGGACCAAAA<br>CGAAAAAAGACGCTCGAAAGCGTCTCTTTTCTGGAATTTGGTACCGAGCT<br>CTAGTA |
| | | Reverse *araC* CDS | TTATGACAACTTGACGGCTACATCATTCACTTTTTCTTCACAACCGGCAC<br>GGAACTCGCTCGGGCTGGCCCCGGTGCATTTTTTAAATACCCGCGAGAAA<br>TAGAGTTGATCGTCAAAACCAACATTGCGACCGACGGTGGCGATAGGCAT<br>CCGGGTGGTGCTCAAAAGCAGCTTCGCCTGGCTGATACGTTGGTCCTCGC<br>GCCAGCTTAAGACGCTAATCCCTAACTGCTGGCGGAAAAGATGTGACAGA<br>CGCGACGGCGACAAGCAAACATGCTGTGCGACGCTGGCGATATCAAAATT<br>GCTGTCTGCCAGGTGATCGCTGATGTACTGACAAGCCTCGCGTACCCGAT<br>TATCCATCGGTGGATGGAGCGACTCGTTAATCGCTTCCATGCGCCGCAGT<br>AACAATTGCTCAAGCAGATTTATCGCCAGCAGCTCCGAATAGCGCCCTTC<br>CCCTTGCCCGGCGTTAATGATTTGCCCAAACAGGTCGCTGAAATGCGGCT<br>GGTGCGCTTCATCCGGGCGAAAGAACCCCGTATTGGCAAATATTGACGGC<br>CAGTTAAGCCATTCATGCCAGTAGGCGCGCGGACGAAAGTAAACCCACTG<br>GTGATACCATTCGCGAGCCTCCGGATGACGACCGTAGTGATGAATCTCTC<br>CTGGCGGGAACAGCAAAATATCACCCGGTCGGCAAACAAATTCTCGTCCC<br>TGATTTTTCACCACCCCCTGACCGCGAATGGTGAGATTGAGAATATAACC<br>TTTCATTCCCAGCGGTCGGTCGATAAAAAAATCGAGATAACCGTTGGCCT<br>CAATCGGCGTTAAACCCGCCACCAGATGGGCATTAAACGAGTATCCCGGC<br>AGCAGGGGATCATTTTGCGCTTCAGCCAT |
| | | $P_{BAD}$ | ACTTTTCATACTCCCGCCATTCAGAGAAGAAACCAATTGTCCATATTGCA<br>TCAGACATTGCCGTCACTGCGTCTTTTACTGGCTCTTCTCGCTAACCAAA<br>CCGGTAACCCCGCTTATTAAAAGCATTCTGTAACAAAGCGGGACCAAAGC<br>CATGACAAAAACGCGTAACAAAAGTGTCTATAATCACGGCAGAAAAGTCC<br>ACATTGATTATTTGCACGGCGTCACACTTTGCTATGCCATAGCATTTTTA<br>TCCATAAGATTAGCGGATCCTACCTGACGCTTTTTATCGCAACTCTCTAC<br>TGTTTCTCCATA<br>(TTS) |
| $P_{lux2}$ | Inducible promoter<br><br>Composite Module<br>(This study) | Reverse Terminator L3S2P55 | CCTAGTTGTCTTCATGCATGAAGACAAAATTAATACTAGAGGGACCAAAA<br>CGAAAAAAGACGCTTTTCAGCGTCTTATTGTTCGTCTTTGGTACCGAGCT<br>CTAGTAGTGATCTACACTAGCACTATCAGTG |
| | | Reverse *luxR* CDS | TTATTAATTTTTAAAGTATGGGCAATCAATTGCTCCTGTTAAAATTGCTT<br>TAGAAATACTTTGGCAGCGGTTTGTTGTATTGAGTTTCATTTGCGCATTG<br>GTTAAATGGAAAGTGACAGTACGCTCACTGCAACCTAATATTTTTGAAAT<br>ATCCCAAGAGCTTTTTCCTTCGCATGCCCACGCTAAACATTCTTTTTCTC<br>TTTTGGTTAAATCGTTGTTTGATTTATTATTTGCTATATTTATTTTTCGA<br>TAATTATCAACTAGAGAAGGAACAATTAATGGTATGTTCATACACGCATG<br>TAAAAATAAACTATCTATATAGTTGTCTTTTTCTGAATGTGCAAAACTAA<br>GCATTCCGAAGCCATTGTTAGCCGTATGAATAGGGAAACTAAACCCAGTG<br>ATAAGACCTGATGTTTTCGCTTCTTTAATTACATTTGGAGATTTTTTATT<br>TACAGCATTGTTTTCAAATATATTCCAATTAATTGGTGAATGATTGGAGT<br>TAGAATAATCTACTATAGGATCATATTTTATTAAATTAGCGTCATCATAA<br>TATTGCCTCCATTTTTTAGGGTAATTATCTAGGATTGAAATATCAGATTT<br>AACCATAGAATGAGGATAAATGATCGCGAGTAAATAATATTCACAATGTA<br>CCATTTTAGTCATATCAGATAAGCATTGATTAATATCATTATTGCTTCTA<br>CAAGCTTTAATTTTATTAATTATTCTGTATGTGTCGTCGGCATTTATGTT<br>TTTCAT |
| | | Reverse RBS B0030 | CTAGTATTTCTCCTCTTTAATCTCTAGTA<br>(Scar/Spacer) |
| | | Reverse J23117 | GCTAGCACAATCCCTAGGACTGAGCTAGCTGTCAATCACACT<br>(Scar/Spacer) |
| | | T7Te Terminator | GGCTCACCTTCGGGTGGGCCTTTCTGCG |

| Name | Type and source | Parts | DNA sequence (5'– 3') |
|---|---|---|---|
| | | $P_{lux2}$ | TTTATATACTAGAGACCTGTAGGATCGTACAGGTTTACGCAAGAAAATGG<br>TTTGTTACTTTCGAATAAAA<br>(TTS, Scar/Spacer) |
| $P_{rhaB}$ | Inducible promoter<br><br>Composite Module<br>(This study) | Reverse Terminator L3S2P55 | CCTAGTTGTCTTCATGCATGAAGACAAAATTAATACTAGAGGGACCAAAA<br>CGAAAAAAGACGCTTTTCAGCGTCTTATTGTTCGTCTTTGGTACCGAGCT<br>AGTA |
| | | Reverse *rhaS* CDS | TTATTGCAGAAAGCCATCCCGTCCCTGGCGAATATCACGCGGTGACCAGT<br>TAAACTCTCGGCGAAAAAGCGTCGAAAAGTGGTTACTGTCGCTGAATCCA<br>CAGCGATAGGCGATGTCAGTAACGCTGGCCTCGCTGTGGCGTAGCAGATG<br>TCGGGCTTTCATCAGTCGCAGGCGGTTCAGGTATCGCTGAGGCGTCAGTC<br>CCGTTTGCTGCTTAAGCTGCCGATGTAGCGTACGCAGTGAAAGAGAAAAT<br>TGATCCGCCACGGCATCCCAATTCACCTCATCGGCAAAATGGTCCTCCAG<br>CCAGGCCAGAAGCAAGTTGAGACGTGATGCGCTGTTTTCCAGGTTCTCCT<br>GCAAACTGCTTTTACGCAGCAAGAGCAGTAATTGCATAAACAAGATCTCG<br>CGACTGGCGGTCGAGGGTAAATCATTTTCCCCTTCCTGCTGTTCCATCTG<br>TGCAACCAGCTGTCGCACCTGCTGCAATACGCTGTGGTTAACGCGCCAGT<br>GAGACGGATACTGCCCATCCAGCTCTTGTGGCAGCAACTGATTCAGCCCG<br>GCGAGAAACTGAAATCGATCCGGCGAGCGATACAGCACATTGGTCAGACA<br>CAGATTATCGGTATGTTCATACAGATGCCGATCATGATCGCGTACGAAAC<br>AGACCGTGCCACCGGTGATGGTATAGGGCTGCCCATTAAACACATGAATA<br>CCCGTGCCATGTTCGACAATCACAATTTCATGAAAATCATGATGATGTTC<br>AGGAAAATCCGCCTGCGGGAGCCGGGGTTCTATCGCCACGGACGCGTTAC<br>CGGACGGAAAAAAATCCACACTATGTAATACGGTCAT |
| | | Reverse J23101-RBS | TTGGGCTCCCTCTAGTAGCTAGCATAATACCTAGGACTGAGCTAGCTGTA<br>AACTCTAGTATCACACT<br>(Scar/Spacer) |
| | | T7Te Terminator | GGCTCACCTTCGGGTGGGCCTTTCTGCGTTTATATACTAGAGAGACCTTT<br>ACGCCGCTGGAGCAGGAATGCGGTGAGCATCACAT<br>(Scar/Spacer) |
| | | $P_{rhaB}$ | CACCACAATTCAGCAAATTGTGAACATCATCACGTTCATCTTTCCCTGGT<br>TGCCAATGGCCCATTTTCCTGTCAGTAACGAGAAGGTCGCGAATTGAGGC<br>GCTTTTTAGACTGGTCGTAA<br>(TTS) |

#### Supplementary Table 6: Sequences for dCas9/dxCas9 generator

| Name | Type and source | Parts | DNA sequence (5'-3') |
|---|---|---|---|
| dCas9 generator | Composite Module (This study) | BioBrick Prefix | GAATTCGCGGCCGCTTCTAGAG |
| | | Reverse Terminator L3S3P00 | GGGAGACCAGAAACAAAAAAAGGGGAGCGGTTTCCCGCTCCCCTTCAATAATT<br>GG |
| | | Reverse Standardized *dCas9* CDS | CTCTAGTATTAGTCACCTCCTAGCTGACTCAAATCAATGCGTGTTTCATAAAG<br>ACCAGTGATGGATTGATGGATAAGAGTGGCATCTAAAACTTCTTTTGTAGACG<br>TATATCGTTTACGATCAATTGTTGTATCAAAATATTTAAAAGCAGCGGGAGCT<br>CCAAGATTCGTCAACGTAAATAAATGAATAATATTTTCTGCTTGTTCACGTAT<br>TGGTTTGTCTCTATGTTTGTTATATGCACTAAGAACTTTATCTAAATTGGCAT<br>CTGCTAAAATAACACGCTTAGAAAATTCACTGATTTGCTCAATAATCTCATCT<br>AAATAATGCTTATGCTGCTCCACAAACAATTGTTTTTGTTCGTTATCTTCTGG<br>ACTACCCTTCAACTTTTCATAATGACTAGCTAAATATAAAAAATTCACATATT<br>TGCTTGGCAGAGCCAGCTCATTTCCTTTTTGTAATTCTCCGGCACTAGCCAGC<br>ATCCGTTTACGACCGTTTTCTAACTCAAAAAGACTATATTTAGGTAGTTTAAT<br>GATTAAGTCTTTTTTAACTTCCTTATATCCTTTAGCTTCTAAAAAGTCAATCG<br>GATTTTTTTCAAAGGAACTTCTTTCCATAATTGTGATCCCTAGTAACTCTTTA<br>ACGGATTTTAACTTCTTCGATTTCCCTTTTTCCACCTTAGCAACCACTAGGAC<br>TGAATAAGCTACCGTTGGACTATCAAAACCACCATATTTTTTTGGATCCCAGT<br>CTTTTTTACGAGCAATAAGCTTGTCCGAATTTCTTTTTGGTAAAATTGACTCC<br>TTGGAGAATCCGCCTGTCTGTACTTCTGTTTTCTTGACAATATTGACTTGGGG<br>CATGGACAATACTTTGCGCACTGTGGCAAAATCTCGCCCTTTATCCCAGACAA<br>TTTCTCCAGTTTCCCCATTAGTTTCGATTAGAGGGCGTTTGCGAATCTCTCCA<br>TTTGCAAGTGTAATTTCTGTTTTGAAGAAGTTCATGATATTAGAGTAAAAGAA<br>ATATTTTGCGGTTGCTTTGCCTATTTCTTGCTCAGACTTAGCAATCATTTTAC<br>GAACATCATAAACTTTATAATCACCATAGACAAACTCCGATTCAAGTTTTGGA<br>TATTTCTTAATCAAAGCAGTTCCAACGACGGCATTTAGATACGCATCATGGGC<br>ATGATGGTAATTGTTAATCTCACGTACTTTATAGAATTGGAAATCTTTTCGGA<br>AGTCAGAAACTAATTTAGATTTTAAGGTAATCACTTTAACCTCTCGAATAAGT<br>TTATCATTTTCATCGTATTTAGTATTCATGCGACTATCCAAAATTTGTGCCAC<br>ATGCTTAGTGATTTGGCGAGTTTCAACCAATTGGCGTTTGATAAAACCAGCTT<br>TATCAAGTTCACTCAAACCTCCACGTTCAGCTTTCGTTAAATTATCAAACTTA<br>CGTTGAGTGATTAACTTGGCGTTTAGAAGTTGTCTCCAATAGTTTTTCATCTT<br>TTTGACTACTTCTTCACTTGGAACGTTATCCGATTTACCACGATTTTTATCAG<br>AACGCGTTAAGACCTTATTGTCTATTGAATCGTCTTTAAGGAAACTTTGTGGA<br>ACAATGGCATCGACATCATAATCACTTAAACGATTAATATCTAATTCTTGGTC<br>CACATACATGTCTCTTCCATTTTGGAGATAATAGAGATAGAGCTTTTCATTTT<br>GCAATTGAGTATTTTCAACAGGATGCTCTTTAAGAATCTGACTTCCTAATTCT<br>TTGATACCTTCTTCGATTCGTTTCATACGCTCTCGCGAATTTTTCTGGCCCTT<br>TTGAGTTGTCTGATTTTCACGTGCCATTTCAATAACGATATTTTCTGGCTTAT<br>GCCGCCCCATTACTTTGACCAATTCATCAACAACTTTTACAGTCTGTAAAATA<br>CCTTTTTTAATAGCAGGGCTACCAGCTAAATTTGCAATATGTTCATGTAAACT<br>ATCGCCTTGTCCAGACACTTGTGCTTTTTGAATGTCTTCTTTAAATGTCAAAC<br>TATCATCATGGATCAGCTGCATAAAATTGCGATTGGCAAAACCATCTGATTTC<br>AAAAAATCTAATATTGTTTTGCCAGATTGCTTATCCCTAATACCATTAATCAA<br>TTTTCGAGACAAACGTCCCCAACCAGTATAACGGCGACGTTTAAGCTGTTTCA<br>TCACCTTATCATCAAAGAGGTGAGCATATGTTTTAAGTCTTTCCTCAATCATC<br>TCCCTATCTTCAAATAAGGTCAATGTTAAAACAATATCCTCTAAGATATCTTC<br>ATTTTCTTCATTATCCAAAAAATCTTTATCTTTAATAATTTTTAGCAAATCAT<br>GGTAGGTACCTAATGAAGCATTAAATCTATCTTCAACTCCTGAAATTTCAACA<br>CTATCAAAACATTCTATTTTTTTGAAATAATCTTCTTTTAATTGCTTAACGGT<br>TACTTTTCGATTTGTTTTGAAGAGTAAATCAACAATGGCTTTCTTCTGTTCAC<br>CTGAAAGAAATGCTGGTTTTCGCATTCCTTCAGTAACATATTTGACCTTTGTC<br>AATTCGTTATAAACCGTAAAATACTCATAAAGCAAACTATGTTTTGGTAGTAC<br>TTTTTCATTTGGAAGATTTTTATCAAAGTTTGTCATGCGTTCAATAAATGATT<br>GAGCTGAAGCACCTTTATCGACAACTTCTTCAAAATTCCATGGGGTAATTGTT<br>TCTTCAGACTTCCGAGTCATCCATGCAAAACGACTATTGCCACGCGCCAATGG<br>ACCAACATAATAAGGAAT**A**CGAAAAGTCAAGATTTTTTCAATCTTCTCACGAT<br>TGTCTTTTAAAAATGGATAAAAGTCTTCTTGTCTTCTCAAAATAGCATGCAGC<br>TCACCCAAGTGAATTTGATGGGGAATAGAGCCGTTGTCAAAGGTCCGTTGCTT<br>GCGCAGCAAATCTTCACGATTTAGTTTCACCAATAATTCCTCAGTACCATCCA<br>TTTTTTCTAAAATTGGTTTGATAAATTTATAAAATTCTTCTTGGCTAGCTCCC |

| Name | Type and source | Parts | DNA sequence (5'-3') |
|---|---|---|---|
| | | | CCATCAATATAACCTGCATATCCGTTTTTTGATTGATCAAAAAAGATTTCTTT<br>ATACTTTTCTGGAAGTTGTTGTCGAACTAAAGCTTTTAAAAGAGTCAAGTCTT<br>GATGATGTTCATCGTAGCGTTTAATCATTGAAGCTGATAGGGGAGCCTTAGTT<br>ATTTCAGTATTTACTCTTAGGATATCTGAAAGTAAAATAGCATCTGATAAATT<br>CTTAGCTGCCAAAAACAAATCAGCATATTGATCTCCAATTTGCGCCAATAAAT<br>TATCTAAATCATCATCGTAAGTATCTTTTGAAAGCTGTAATTTAGCATCTTCT<br>GCCAAATCAAAATTTGATTTAAAATTAGGGGTCAAACCCAATGACAAAGCAAT<br>GAGATTCCCAAATAAGCCATTTTTCTTCTCACCGGGGAGCTGAGCAATGAGAT<br>TTTCTAATCGTCTTGATTTACTCAATCGTGCAGAAAGAATCGCTTTAGCATCT<br>ACTCCACTTGCGTTAATAGGGTTTTCTTCAAATAATTGATTGTAGGTTTGTAC<br>CAACTGGATAAATAGTTTGTCCACATCACTATTATCAGGATTTAAATCTCCCT<br>CAATCAAAAAATGACCACGAAACTTAATCATATGCGCTAAGGCCAAATAGATT<br>AAGCGCAAATCCGCTTTATCAGTAGAATCTACCAATTTTTTTCGCAGATGATA<br>GATAGTTGGATATTTCTCATGATAAGCAACTTCATCTACTATATTTCCAAAAA<br>TAGGATGACGTTCATGCTTCTTGTCTTCTTCCACCAAAAAAGACTCTTCAAGT<br>CGATGAAAGAAACTATCATCTACTTTCGCCATCTCATTTGAAAAAATCTCCTG<br>TAGATAACAAATACGATTCTTCCGACGTGTATACCTTCTACGAGCTGTCCGTT<br>TGAG**G**CGAGTCGCTTCCGCTGTCTCTCCACTGTCAAATAAAAGAGCCCCTATA<br>AGATTTTTTTTGATACTGTGGCGGTCTGTATTTCCCAGAACCTTGAACTTTTT<br>AGACGGAACCTTATATTCATCAGTGATCACCGCCCATCCGACGCTATTTGTGC<br>CGATAGCTAAGCCTATTGAGTATTTCTTATCCAT<br>(Scar/Spacer, **mutation**) |
| | | Reverse RBS B0034 | AGATCCTTTCTCCTCTTT<br><br>(Scar/Spacer) |
| | | *tetR/tetA* promoters | AGATCTTTT**C**AATTCTTTTCTCTATCACTGATAGGGAGTGGTAAAATAACTCT<br>ATCAACGATAGAGTGTCAACAAAAATTAGGAATTAATG<br>(**mutation**) |
| | | *tetR* CDS | ATGTC**A**AGATTAGATAAAAGTAAAGTGATTAACAGCGCATTAGAGCTGCTTAA<br>TGAGGTCGGAATCGAAGGTTTAACAACCCGTAAACTCGCCCAGAAGCTAGGTG<br>TAGAGCAGCCTACATTGTATTGGCATGTAAAAAATAAGCGGGCTTTGCTCGAC<br>GCCTTAGCCATTGAGATGTTAGATAGGCACCATACTCACTTTTGCCCTTTAGA<br>AGGGGAAAGCTGGCAAGATTTTTTACGTAATAACGCTAAAAGTTTTAGATGTG<br>CTTTACTAAGTCATCGCGATGGAGCAAAAGTACATTTAGGTACACGGCCTACA<br>GAAAAACAGTATGAAACTCTCGAAAATCAATTAGCCTTTTTATGCCAACAAGG<br>TTTTTCACTAGAGAATGCATTATATGCACTCAGCGCTGTGGGGCATTTTACTT<br>TAGGTTGCGTATTGGAAGATCAAGAGCATCAAGTCGCTAAAGAAGAAAGGGAA<br>ACACCTACTACTGATAGTATGCCGCCATTATTACGACAAGCTATCGAATTATT<br>TGATCACCAAGGTGCAGAGCCAGCCTTCTTATTCGGCCTTGAATTGATCATAT<br>GCGGATTAGAAAAACAACTTAAATGTGAAAGTGGGTCTTAA<br>(**mutation**) |
| | | Terminator L3S3P22 | TACTAGAGCCAATTATTGAAGGCCGCTAACGCGGCCTTTTTTTGTTTCTGGTC<br>TCCC<br>(Scar/Spacer) |
| | | BioBrick Suffix | TACTAGTAGCGGCCGCTGCAG |
| dxCas9 generator | Composite Module (This study) | BioBrick Prefix | GAATTCGCGGCCGCTTCTAGAG |
| | | Reverse Terminator L3S3P00 | GGGAGACCAGAAACAAAAAAAGGGGAGCGGTTTCCCGCTCCCCTTCAATAATT<br>GG |
| | | Reverse Standardized *dxCas9* CDS | CTCTAGTATTAGTCACCTCCTAGCTGACTCAAATCAATGCGTGTTTCATAAAG<br>ACCAGTGATGGATTGATGGATAAGAGTGGCATCTAAAACTTCTTTTGTAGACG<br>TATATCGTTTACGATCAATTGTTGTATCAAAATATTTAAAAGCAGCGGGAGCT<br>CCAAGATTCGTCAACGTAAATAAATGAATAATATTTTCTGCTTGTTCACGTAT<br>TGGTTTGTCTCTATGTTTGTTATATGCACTAAGAACTTTATCTAAATTGGCAT<br>CTGCTAAAATAACACGCTTAGAAAATTCACTGATTTGCTCAATAATCTCATCT<br>AAATAATGCTTATGCTGCTCCACAAACAATTGTTTTTGTTCGTTATCTTCTGG |

| Name | Type and source | Parts | DNA sequence (5'-3') |
| --- | --- | --- | --- |
| | | | ACTACCCTTCAACTTTTCATAATGACTAGCTAAATATAAAAAATTCACATATT<br>TGCTTGGCAGAGCCAGCTCATTTCCTTTTTGTAA**AA**CTCCGGCACTAGCCAGC<br>ATCCGTTTACGACCGTTTTCTAACTCAAAAAGACTATATTTAGGTAGTTTAAT<br>GATTAAGTCTTTTTTAACTTCCTTATATCCTTTAGCTTCTAAAAAGTCAATCG<br>GATTTTTTTCAAAGGAACTTCTTTCCATAATTGTGATCCCTAGTAACTCTTTA<br>ACGGATTTTAACTTCTTCGATTTCCCTTTTTCCACCTTAGCAACCACTAGGAC<br>TGAATAAGCTACCGTTGGACTATCAAAACCACCATATTTTTTTGGATCCCAGT<br>CTTTTTTACGAGCAATAAGCTTGTCCGAATTTCTTTTTGGTAAAATTGACTCC<br>TTGGAGAATCCGCCTGTCTGTACTTCTGTTTTCTTGACAATATTGACTTGGGG<br>CATGGACAATACTTTGCGCACTGTGGCAAAATCTCGCCCTTTATCCCAGACAA<br>TTTCTCCAGTTTCCCCATTAGTTTCGATTAGAGGGCGTTTGCGAATCTCTCCA<br>TTTGCAAGTGTAATTTCTGTTTTGAAGAAGTTCATGATATTAGAGTAAAAGAA<br>ATATTTTGCGGTTGCTTTGCCTATTTCTTGCTCAGACTTAGCAATCATTTTAC<br>GAACATCATAAACTTTATAATCACCATAGACAAACTCCGATTCAAGTTTTGGA<br>TATTTCTTAATCAAAGCAGTTCCAACGACGGCATTTAGATACGCATCATGGGC<br>ATGATGGTAATTGTTAATCTCACGTACTTTATAGAATTGGAAATCTTTTCGGA<br>AGTCAGAAACTAATTTAGATTTTAAGGTAATCACTTTAACCTCTCGAATAAGT<br>TTATCATTTTCATCGTATTTAGTATTCATGCGACTATCCAAAATTTGTGCCAC<br>ATGCTTAGTGATTTGGCGAGTTTCAACCAATTGGCGTTTGATAAAACCAGCTT<br>TATCAAGTTCACTCAAACCTCCACGTTCAGCTTTCGTTAAATTATCAAACTTA<br>CGTTGAGTGATTAACTTGGCGTTTAGAAGTTGTCTCCAATAGTTTTTCATCTT<br>TTTGACTACTTCTTCACTTGGAACGTTATCCGATTTACCACGATTTTTATCAG<br>AACGCGTTAAGACCTTATTGTCTATTGAATCGTCTTTAAGGAAACTTTGTGGA<br>ACAATGGCATCGACATCATAATCACTTAAACGATTAATATCTAATTCTTGGTC<br>CACATACATGTCTCTTCCATTTTGGAGATAATAGAGATAGAGCTTTTCATTTT<br>GCAATTGAGTATTTTCAACAGGATGCTCTTTAAGAATCTGACTTCCTAATTCT<br>TTGATACCTTCTTCGATTCGTTTCATACGCTCTCGCGAATTTTTCTGGCCCTT<br>TTGAGTTGTCTGATTTTCACGTGCCATTTCAATAACGATATTTTCTGGCTTAT<br>GCCGCCCCATTACTTTGACCAATTCATCAACAACTTTTACAGTCTGTAAAATA<br>CCTTTTTTAATAGCAGGGCTACCAGCTAAATTTGCAATATGTTCATGTAAACT<br>ATCGCCTTGTCCAGACACTTGTGCTTTTTGAATGTCTTCTTTAAATGTCAAAC<br>TATCATCATGGATCAGCTG**A**ATAAAATTGCGATTGGCAAAACCATCTGATTTC<br>AAAAAATCTAATATTGTTTTGCCAGATTGCTTATCCCTAATACCATTAATCAA<br>TTTTCGAGACAAACGTCCCCAACCAGTATAACGGCGACGTTTAAGCTGTTTCA<br>TCACCTTATCATCAAAGAGGTGAGCATATGTTTTAAGTCTTTCCTCAATCATC<br>TCCCTATCTTCAAATAAGGTCAATGTTAAAACAATATCCTCTAAGATATCTTC<br>ATTTTCTTCATTATCCAAAAAATCTTTATCTTTAATAATTTTTAGCAAATCAT<br>GGTAGGTACCTAATGAAGCATTAAATCTATCTTCAACTCCTGAAATTTCAACA<br>CTATCAAAACATTCTATTTTTTTGAAATAATCTTCTTTTAATTGCTTAACGGT<br>TACTTTTCGATTTGTTTTGAAGAGTAAATCAACAATGGCTTTCTTCTG**A**TCAC<br>CTGAAAGAAATGCTGGTTTTCGCATTCCTTCAGTAACATATTTGACCTTTGTC<br>AATTCGTTATAAACCGTAAAATACTCATAAAGCAAACTATGTTTTGGTAGTAC<br>TTTTTCATTTGGAAGATTTTTATCAAAGTTTGTCATGCGTTCAATAAATGATT<br>GAGCTGAAGCACCTTTATCGACAACTT**T**TTCAAAATTCCATGGGGTAATTGTT<br>TCTTCAGACTTCCGAGTCATCCATGCAAAACGACTATTGCCACGCGCCAATGG<br>ACCAACATAATAAGGAATACGAAAAGTCAAGATTTTTTCAATCTTCTCACGAT<br>TGTCTTTTAAAAATGGATAAAAGTCTTCTTGTCTTCTCAAAATAGCATGCAGC<br>TCACCCAAGTGAATTTGATGGGGAATA**AT**GCCGTTGTCAAAGGTCCGTTGCTT<br>GCGCAGCAAATCTTCACGATTTAGTTTCACCAATAATTCCTCAGTACCATCCA<br>TTTTTTCTAAAATTGGTTTGATAAATTTATAAAATTCTTCTTGGCTAGCTCCC<br>CCATCAATATAACCTGCATATCCGTTTTTTGATTGATCAAAAAAGATTTCTTT<br>ATACTTTTCTGGAAGTTGTTGTCGAACTAAAGCTTTTAAAAGAGTCAAGTCTT<br>GATGATGTTCATCGTA**TAA**TTTAATCATTGAAGCTGATAGGGGAGCCTTAGTT<br>ATTTCAGTATTTACTCTTAGGATATCTGAAAGTAAAATAGCATCTGATAAATT<br>CTTAGCTGCCAAAAACAAATCAGCATATTGATCTCCAATTTGCGCCAATAAAT<br>TATCTAAATCATCATCGTAAGTATCTTTTGAAAGCTGTAATTT**G**G**T**ATCTTCT<br>GCCAAATCAAAATTTGATTTAAAATTAGGGGTCAAACCCAATGACAAAGCAAT<br>GAGATTCCCAAATAAGCCATTTTTCTTCTCACCGGGGAGCTGAGCAATGAGAT<br>TTTCTAATCGTCTTGATTTACTCAATCGTGCAGAAAGAATCGCTTTAGCATCT<br>ACTCCACTTGCGTTAATAGGGTTTTCTTCAAATAATTGATTGTAGGTTTGTAC<br>CAACTGGATAAATAGTTTGTCCACATCACTATTATCAGGATTTAAATCTCCCT<br>CAATCAAAAAATGACCACGAAACTTAATCATATGCGCTAAGGCCAAATAGATT<br>AAGCGCAAATCCGCTTTATCAGTAGAATCTACCAATTTTTTTCGCAGATGATA<br>GATAGTTGGATATTTCTCATGATAAGCAACTTCATCTACTATATTTCCAAAAA<br>TAGGATGACGTTCATGCTTCTTGTCTTCTTCCACCAAAAAAGACTCTTCAAGT<br>CGATGAAAGAAACTATCATCTACTTTCGCCATCTCATTTGAAAAAATCTCCTG<br>TAGATAACAAATACGATTCTTCCGACGTGTATACCTTCTACGAGCTGTCCGTT<br>TGAGGCGAGTCGCTTCCGCTGTCTCTCCACTGTCAAATAAAAGAGCCCCTATA |

| Name | Type and source | Parts | DNA sequence (5'-3') |
|---|---|---|---|
| | | | AGATTTTTTTTGATACTGTGGCGGTCTGTATTTCCCAGAACCTTGAACTTTTT<br>AGACGGAACCTTATATTCATCAGTGATCACCGCCCATCCGACGCTATTTGTGC<br>CGATAGCTAAGCCTATTGAGTATTTCTTATCCAT<br>(Scar/Spacer, **mutation**) |
| | | Reverse RBS B0034 | AGATCCTTTCTCCTCTTT<br><br>(Scar/Spacer) |
| | | *tetR/tetA* promoters | AGATCTTTT**C**AATTCTTTTCTCTATCACTGATAGGGAGTGGTAAAATAACTCT<br>ATCAACGATAGAGTGTCAACAAAAATTAGGAATTAATG<br>(**mutation**) |
| | | *tetR* CDS | ATGTC**A**AGATTAGATAAAAGTAAAGTGATTAACAGCGCATTAGAGCTGCTTAA<br>TGAGGTCGGAATCGAAGGTTTAACAACCCGTAAACTCGCCCAGAAGCTAGGTG<br>TAGAGCAGCCTACATTGTATTGGCATGTAAAAAATAAGCGGGCTTTGCTCGAC<br>GCCTTAGCCATTGAGATGTTAGATAGGCACCATACTCACTTTTGCCCTTTAGA<br>AGGGGAAAGCTGGCAAGATTTTTTACGTAATAACGCTAAAAGTTTTAGATGTG<br>CTTTACTAAGTCATCGCGATGGAGCAAAAGTACATTTAGGTACACGGCCTACA<br>GAAAAACAGTATGAAACTCTCGAAAATCAATTAGCCTTTTTATGCCAACAAGG<br>TTTTTCACTAGAGAATGCATTATATGCACTCAGCGCTGTGGGGCATTTTACTT<br>TAGGTTGCGTATTGGAAGATCAAGAGCATCAAGTCGCTAAAGAAGAAAGGGAA<br>ACACCTACTACTGATAGTATGCCGCCATTATTACGACAAGCTATCGAATTATT<br>TGATCACCAAGGTGCAGAGCCAGCCTTCTTATTCGGCCTTGAATTGATCATAT<br>GCGGATTAGAAAAACAACTTAAATGTGAAAGTGGGTCTTAA<br>(**mutation**) |
| | | Terminator L3S3P22 | TACTAGAGCCAATTATTGAAGGCCGCTAACGCGGCCTTTTTTTGTTTCTGGTC<br>TCCC<br>(Scar/Spacer) |
| | | BioBrick Suffix | TACTAGTAGCGGCCGCTGCAG |

#### Supplementary Table 7: Sequences for activator generator and reporter

| Name | Type and source | Parts | DNA sequence (5'-3') |
|---|---|---|---|
| PspFΔHTH::λN22plus | Composite Module (This study) | RBS B0032 | TCACACAGGAAAGTACTAG<br>(Scar/Spacer) |
| | | *pspFΔHTH::λN22plus* CDS | ATGGCAGAATACAAAGATAATTTACTTGGTGAGGCGAACAGCTTTCTCGAA<br>GTGCTGGAACAGGTTTCGCATCTCGCACCGCTGGACAAACCGGTGCTCATC<br>ATCGGCGAACGCGGCACCGGTAAAGAGCTGATTGCCAGCCGCCTGCATTAT<br>CTCTCCTCCCGTTGGCAAGGGCCGTTTATTTCCCTTAACTGCGCGGCGTTA<br>AATGAAAATCTGCTGGATTCCGAACTGTTTGGTCACGAAGCGGGGGCGTTT<br>ACCGGTGCGCAAAAACGTCATCCAGGGAGATTTGAACGTGCCGACGGCGGT<br>ACGCTATTTCTTGATGAACTCGCTACGGCACCGATGATGGTGCAGGAGAAA<br>TTATTGCGCGTGATTGAGTACGGTGAACTGGAGCGCGTTGGCGGCAGCCAA<br>CCATTGCAGGTGAATGTGCGGTTGGTATGCGCGACGAATGCCGATCTCCCG<br>GCGATGGTCAATGAAGGCACTTTTCGCGCTGACCTGCTCGACCGACTGGCT<br>TTTGATGTTGTACAACTGCCACCACTGCGCGAGCGCGAAAGCGACATAATG<br>TTGATGGCAGAATACTTTGCCATCCAGATGTGTCGGGAAATCAAGCTGCCT<br>CTGTTCCCGGGGTTTACGGAGCGCGCCAGAGAAACATTGCTGAATTATCGT<br>TGGCCGGGAAATATTCGTGAATTGAAAAACGTGGTGGAACGTTCAGTGTAT<br>CGCCACGGCACCAGCGATTATCCGCTTGATGACATCATTATTGATCCCTTT<br>AAACGGCGTCCGCCTGAAGACGCTATCGCCGTTTCAGAAACCACCTCGCTT<br>CCAACACTGCCGCTGGATTTACGTGAGTTTCAGATGCAGCAGGAAAAAGAG<br>TTGCTGCAACTCAGTTTGCAAATGAATGCACGCACACGCCGCCGCGAACGT<br>CGCGCAGAGAAACAGGCTCAATGGAAAGCAGCAAATTAA<br>(λN22plus) |
| | | Terminator L3S3P22 | TACTAGAGCCAATTATTGAAGGCCGCTAACGCGGCCTTTTTTTGTTTCTGG<br>TCTCCC<br>(Scar/Spacer) |
| Reporter with sfGFP::ASV tag | Composite Module (This study) | RiboJ | AGCTGTCACCGGATGTGCTTTCCGGTCTGATGAGTCCGTGAGGACGAAACA<br>GCCTCTACAAATAATTTTGTTTAACTAGAG |
| | | RBS B0030 | ATTAAAGAGGAGAAATACCAT<br>(Scar/Spacer) |
| | | *sfgfp* CDS | ATGCGTAAAGGCGAAGAGCTGTTCACTGGTGTCGTCCCTATTCTGGTGGAA<br>CTGGATGGTGATGTCAACGGTCATAAGTTTTCCGTGCGTGGCGAGGGTGAA<br>GGTGACGCAACTAATGGTAAACTGACGCTGAAGTTCATCTGTACTACTGGT<br>AAACTGCCGGTACCTTGGCCGACTCTGGTAACGACGCTGACTTATGGTGTT<br>CAGTGCTTTGCTCGTTATCCGGACCATATGAAGCAGCATGACTTCTTCAAG<br>TCCGCCATGCCGGAAGGCTATGTGCAGGAACGCACGATTTCCTTTAAGGAT<br>GACGGCACGTACAAAACGCGTGCGGAAGTGAAATTTGAAGGCGATACCCTG<br>GTAAACCGCATTGAGCTGAAAGGCATTGACTTTAAAGAAGACGGCAATATC<br>CTGGGCCATAAGCTGGAATACAATTTTAACAGCCACAATGTTTACATCACC<br>GCCGATAAACAAAAAAATGGCATTAAAGCGAATTTTAAAATTCGCCACAAC<br>GTGGAGGATGGCAGCGTGCAGCTGGCTGATCACTACCAGCAAAACACTCCA<br>ATCGGTGATGGTCCTGTTCTGCTGCCAGACAATCACTATCTGAGCACGCAA<br>AGCGTTCTGTCTAAAGATCCGAACGAGAAACGCGATCATATGGTTCTGCTG<br>GAGTTCGTAACCGCAGCGGGCATCACGCATGGTATGGATGAACTGTACAAA<br>**AGGCCTGCTGCAAACGACGAAAACTACGCTGCATCAGTTTAA**<br>(**ASV tag**) |
| | | Terminator L3S3P21 | TAATACTAGAGCCAATTATTGAAGGCCTCCCTAACGGGGGGCCTTTTTTTG<br>TTTCTGGTCTCCC<br>(Scar/Spacer) |
| RiboJ | Insulator | | AGCTGTCACCGGATGTGCTTTCCGGTCTGATGAGTCCGTGAGGACGAAACAGCCTCTACAAATA<br>ATTTTGTTTAA |

#### Supplementary Table 8: Sequences of gRNA, mRNA and related genetic elements

| Part name | Type and source | DNA sequence (5'– 3') |
|---|---|---|
| sgRNA-LEA2 | sgRNA (Previous study) | CTAGAAGTTATTATATAGTTCGGTCGTTTGAGAGCTAGGGCCCTGAAGAAGGGCCCTAGCA<br>AGTTCAAATAAGGCTAGTCCGTTATCAACTTGGGCCCTGAAGAAGGGCCCAAGTGGCACCG<br>AGTCGGTGCTTTTTTTGAAGCTTCTCGGTACCAAATTCCAGAAAAGAGGCCTCCCGAAAGG<br>GGGGCCTTTTTTCGTTTTGGTCC<br>(Scar, Spacer, sgRNA scaffold (Double BoxB), Terminator L3S2P21) |
| sgRNA-LEA2-ex2 | sgRNA (This study) | CTAGAAGTTATTATATAGTTCGGTCGTTTGAGAGCTAGAAATAGCAAGTTCAAATAAGGCT<br>AGTCCGTTATCAACTTGAAAAAGTGGCACCGAGTCGGTGCgggccctgaagaagGGCCCTA<br>GCAAGTTCAAATAAGGCTAGTCCGTTATCAACTTGGGCCCTGAAGAAGGGCCCAAGTGGCA<br>CCGAGTCGGTGCTTTTTTTTACTAGAGCTCGGTACCAAATTCCAGAAAAGAGGCCTCCCGA<br>AAGGGGGGCCTTTTTTCGTTTTGGTCC<br>(Scar, Spacer, sgRNA scaffold (Double BoxB), Terminator L3S2P21) |
| sgRNA-LEA2-ex3 | sgRNA (This study) | CTAGAAGTTATTATATAGTTCGGTCGTTTGAGAGCTAGAAATAGCAAGTTCAAATAAGGCT<br>AGTCCGTTATCAACTTGAAAAAGTGGCACCGAGTCGGTGCgggccctgaagaagGGCCCAA<br>GGCTAGGGCCCTGAAGAAGGGCCCGCAGGGCCCTGAAGAAGGGCCCTTTTTTTTACTAGAG<br>CTCGGTACCAAATTCCAGAAAAGAGGCCTCCCGAAAGGGGGGCCTTTTTTCGTTTTGGTCC<br>(Scar, Spacer, sgRNA scaffold (Double BoxB), Terminator L3S2P21) |
| sgRNA-LEB3-ex2 | sgRNA (This study) | CTAGAGCATAGTTCGTTTCCCATGTTTGAGAGCTAGAAATAGCAAGTTCAAATAAGGCTAG<br>TCCGTTATCAACTTGAAAAAGTGGCACCGAGTCGGTGCgggccctgaagaagGGCCCTAGC<br>AAGTTCAAATAAGGCTAGTCCGTTATCAACTTGGGCCCTGAAGAAGGGCCCAAGTGGCACC<br>GAGTCGGTGCTTTTTTTTACTAGAGCTCGGTACCAAATTCCAGAAAAGAGGCCTCCCGAAA<br>GGGGGGCCTTTTTTCGTTTTGGTCC<br>(Scar, Spacer, sgRNA scaffold (Double BoxB), Terminator L3S2P21) |
| sgRNA-LEB3-ex3 | sgRNA (This study) | CTAGAGCATAGTTCGTTTCCCATGTTTGAGAGCTAGAAATAGCAAGTTCAAATAAGGCTAG<br>TCCGTTATCAACTTGAAAAAGTGGCACCGAGTCGGTGCgggccctgaagaagGGCCCAAGG<br>CTAGGGCCCTGAAGAAGGGCCCGCAGGGCCCTGAAGAAGGGCCCTTTTTTTTACTAGAGCT<br>CGGTACCAAATTCCAGAAAAGAGGCCTCCCGAAAGGGGGGCCTTTTTTCGTTTTGGTCC<br>(Scar, Spacer, sgRNA scaffold (Double BoxB), Terminator L3S2P21) |
| crRNA-LEA2-WT | crRNA (This study) | CTAGAAGTTATTATATAGTTCGGTCGTTTTAGAGCTATGCTGTTTTGTACTAGAGCCAATT<br>ATTGAAGGCCGCTAACGCGGCCTTTTTTTGTTTCTGGTCTCCC<br>(Scar, Spacer, repeat, Terminator L3S3P22) |
| crRNA-LEA2-d1 | crRNA (This study) | CTAGAAGTTATTATATAGTTCGGTCGTTTTAGAGCTTTGCTGTTTTGTACTAGAGCCAATT<br>ATTGAAGGCCGCTAACGCGGCCTTTTTTTGTTTCTGGTCTCCC<br>(Scar, Spacer, repeat, Terminator L3S3P22) |
| crRNA-LEA2-d2 | crRNA (This study) | CTAGAAGTTATTATATAGTTCGGTCGTTTTAGAGCATTGCTGTTTTGTACTAGAGCCAATT<br>ATTGAAGGCCGCTAACGCGGCCTTTTTTTGTTTCTGGTCTCCC<br>(Scar, Spacer, repeat, Terminator L3S3P22) |
| crRNA-LEA2-d3 | crRNA (This study) | CTAGAAGTTATTATATAGTTCGGTCGTTTTAGAGGATTGCTGTTTTGTACTAGAGCCAATT<br>ATTGAAGGCCGCTAACGCGGCCTTTTTTTGTTTCTGGTCTCCC<br>(Scar, Spacer, repeat, Terminator L3S3P22) |
| crRNA-LEA2-d4 | crRNA (This study) | CTAGAAGTTATTATATAGTTCGGTCGTTTTAGACGATTGCTGTTTTGTACTAGAGCCAATT<br>ATTGAAGGCCGCTAACGCGGCCTTTTTTTGTTTCTGGTCTCCC<br>(Scar, Spacer, repeat, Terminator L3S3P22) |
| crRNA-LEA2-U1 | crRNA (This study) | CTAGAAGTTATTATATAGTTCGGTCGTTTTTGACGATTGCTGTTTTGTACTAGAGCCAATT<br>ATTGAAGGCCGCTAACGCGGCCTTTTTTTGTTTCTGGTCTCCC<br>(Scar, Spacer, repeat, Terminator L3S3P22) |
| crRNA-LEA2-U2 | crRNA (This study) | CTAGAAGTTATTATATAGTTCGGTCGTTTATGACGATTGCTGTTTTGTACTAGAGCCAATT<br>ATTGAAGGCCGCTAACGCGGCCTTTTTTTGTTTCTGGTCTCCC<br>(Scar, Spacer, repeat, Terminator L3S3P22) |
| crRNA-LEA2-U3 | crRNA (This study) | CTAGAAGTTATTATATAGTTCGGTCGTTAATGACGATTGCTGTTTTGTACTAGAGCCAATT<br>ATTGAAGGCCGCTAACGCGGCCTTTTTTTGTTTCTGGTCTCCC<br>(Scar, Spacer, repeat, Terminator L3S3P22) |

| Part name | Type and source | DNA sequence (5'– 3') |
|---|---|---|
| crRNA-LEA2-U4 | crRNA (This study) | CTAGAAGTTATTATATAGTTCGGTCGTAAATGACGATTGCTGTTTTGTACTAGAGCCAATT<br>ATTGAAGGCCGCTAACGCGGCCTTTTTTTGTTTCTGGTCTCCC<br>(Scar, Spacer, repeat, Terminator L3S3P22) |
| crRNA-LEA2-U5 | crRNA (This study) | CTAGAAGTTATTATATAGTTCGGTCGAAAATGACGATTGCTGTTTTGTACTAGAGCCAATT<br>ATTGAAGGCCGCTAACGCGGCCTTTTTTTGTTTCTGGTCTCCC<br>(Scar, Spacer, repeat, Terminator L3S3P22) |
| crRNA-LEA2-U5A | crRNA (This study) | CTAGAAGTTATTATATAGTTCGGTCAAAAATGACGATTGCTGTTTTGTACTAGAGCCAATT<br>ATTGAAGGCCGCTAACGCGGCCTTTTTTTGTTTCTGGTCTCCC<br>(Scar, Spacer, repeat, Terminator L3S3P22) |
| crRNA-LEA2-U5C | crRNA (This study) | CTAGAAGTTATTATATAGTTCGGTCCAAAATGACGATTGCTGTTTTGTACTAGAGCCAATT<br>ATTGAAGGCCGCTAACGCGGCCTTTTTTTGTTTCTGGTCTCCC<br>(Scar, Spacer, repeat, Terminator L3S3P22) |
| crRNA-LEA2-U5T | crRNA (This study) | CTAGAAGTTATTATATAGTTCGGTCTAAAATGACGATTGCTGTTTTGTACTAGAGCCAATT<br>ATTGAAGGCCGCTAACGCGGCCTTTTTTTGTTTCTGGTCTCCC<br>(Scar, Spacer, repeat, Terminator L3S3P22) |
| crRNA-LEB3-WT | crRNA (This study) | CTAGAGCATAGTTCGTTTCCCATGTTTTAGAGCTATGCTGTTTTGTACTAGAGCCAATTAT<br>TGAAGGCCGCTAACGCGGCCTTTTTTTGTTTCTGGTCTCCC<br>(Scar, Spacer, repeat, Terminator L3S3P22) |
| tracrRNA-WT | tracrRNA (This study) | CTAGAGGAACCATTCAAAACAGCATAGCAAGTTAAAATAAGGCTAGTCCGTTATCAACTTG<br>AAAAAGTGGCACCGAGTCGGTGCGGGCCCTGAAGAAGGGCCCAAGGCTAGGGCCCTGAAGA<br>AGGGCCCGCAGGGCCCTGAAGAAGGGCCCTTTTTTTTACTAGAGCCAATTATTGAAGGCCG<br>CTAACGCGGCCTTTTTTTGTTTCTGGTCTCCC<br>(Scar, Scaffold, anti-repeat, bulge, Terminator L3S3P22) |
| tracrRNA-d1 | tracrRNA (This study) | CTAGAGGAACCATTCAAAACAGCAAAGCAAGTTAAAATAAGGCTAGTCCGTTATCAACTTG<br>AAAAAGTGGCACCGAGTCGGTGCgggccctgaagaagGGCCCAAGGCTAGGGCCCTGAAGA<br>AGGGCCCGCAGGGCCCTGAAGAAGGGCCCTTTTTTTTACTAGAGCCAATTATTGAAGGCCG<br>CTAACGCGGCCTTTTTTTGTTTCTGGTCTCCC<br>(Scar, Scaffold, anti-repeat, bulge, Terminator L3S3P22) |
| tracrRNA-d2 | tracrRNA (This study) | CTAGAGGAACCATTCAAAACAGCAATGCAAGTTAAAATAAGGCTAGTCCGTTATCAACTTG<br>AAAAAGTGGCACCGAGTCGGTGCgggccctgaagaagGGCCCAAGGCTAGGGCCCTGAAGA<br>AGGGCCCGCAGGGCCCTGAAGAAGGGCCCTTTTTTTTACTAGAGCCAATTATTGAAGGCCG<br>CTAACGCGGCCTTTTTTTGTTTCTGGTCTCCC<br>(Scar, Scaffold, anti-repeat, bulge, Terminator L3S3P22) |
| tracrRNA-d3 | tracrRNA (This study) | CTAGAGGAACCATTCAAAACAGCAATCCAAGTTAAAATAAGGCTAGTCCGTTATCAACTTG<br>AAAAAGTGGCACCGAGTCGGTGCGGGCCCTGAAGAAGGGCCCAAGGCTAGGGCCCTGAAGA<br>AGGGCCCGCAGGGCCCTGAAGAAGGGCCCTTTTTTTTACTAGAGCCAATTATTGAAGGCCG<br>CTAACGCGGCCTTTTTTTGTTTCTGGTCTCCC<br>(Scar, Scaffold, anti-repeat, bulge, Terminator L3S3P22) |
| tracrRNA-d4 | tracrRNA (This study) | CTAGAGGAACCATTCAAAACAGCAATCGAAGTTAAAATAAGGCTAGTCCGTTATCAACTTG<br>AAAAAGTGGCACCGAGTCGGTGCGGGCCCTGAAGAAGGGCCCAAGGCTAGGGCCCTGAAGA<br>AGGGCCCGCAGGGCCCTGAAGAAGGGCCCTTTTTTTTACTAGAGCCAATTATTGAAGGCCG<br>CTAACGCGGCCTTTTTTTGTTTCTGGTCTCCC<br>(Scar, Scaffold, anti-repeat, bulge, Terminator L3S3P22) |
| tracrRNA-U1 | tracrRNA (This study) | CTAGAGGAACCATTCAAAACAGCAATCGAAGTAAAAATAAGGCTAGTCCGTTATCAACTTG<br>AAAAAGTGGCACCGAGTCGGTGCGGGCCCTGAAGAAGGGCCCAAGGCTAGGGCCCTGAAGA<br>AGGGCCCGCAGGGCCCTGAAGAAGGGCCCTTTTTTTTACTAGAGCCAATTATTGAAGGCCG<br>CTAACGCGGCCTTTTTTTGTTTCTGGTCTCCC<br>(Scar, Scaffold, anti-repeat, bulge, Terminator L3S3P22) |
| tracrRNA-U2 | tracrRNA (This study) | CTAGAGGAACCATTCAAAACAGCAATCGAAGTATAAATAAGGCTAGTCCGTTATCAACTTG<br>AAAAAGTGGCACCGAGTCGGTGCGGGCCCTGAAGAAGGGCCCAAGGCTAGGGCCCTGAAGA<br>AGGGCCCGCAGGGCCCTGAAGAAGGGCCCTTTTTTTTACTAGAGCCAATTATTGAAGGCCG<br>CTAACGCGGCCTTTTTTTGTTTCTGGTCTCCC<br>(Scar, Scaffold, anti-repeat, bulge, Terminator L3S3P22) |
| tracrRNA-U3 | tracrRNA (This study) | CTAGAGGAACCATTCAAAACAGCAATCGAAGTATTAATAAGGCTAGTCCGTTATCAACTTG<br>AAAAAGTGGCACCGAGTCGGTGCGGGCCCTGAAGAAGGGCCCAAGGCTAGGGCCCTGAAGA |

| Part name | Type and source | DNA sequence (5'– 3') |
|---|---|---|
| | | AGGGCCCGCAGGGCCCTGAAGAAGGGCCCTTTTTTTTACTAGAGCCAATTATTGAAGGCCG<br>CTAACGCGGCCTTTTTTTGTTTCTGGTCTCCC<br>(Scar, Scaffold, anti-repeat, bulge, Terminator L3S3P22) |
| tracrRNA-U4 | tracrRNA (This study) | CTAGAGGAACCATTCAAAACAGCAATCGAAGTATTTATAAGGCTAGTCCGTTATCAACTTG<br>AAAAAGTGGCACCGAGTCGGTGCGGGCCCTGAAGAAGGGCCCAAGGCTAGGGCCCTGAAGA<br>AGGGCCCGCAGGGCCCTGAAGAAGGGCCCTTTTTTTTACTAGAGCCAATTATTGAAGGCCG<br>CTAACGCGGCCTTTTTTTGTTTCTGGTCTCCC<br>(Scar, Scaffold, anti-repeat, bulge, Terminator L3S3P22) |
| tracrRNA-U5 | tracrRNA (This study) | CTAGAGGAACCATTCAAAACAGCAATCGAAGTATTTTTAAGGCTAGTCCGTTATCAACTTG<br>AAAAAGTGGCACCGAGTCGGTGCGGGCCCTGAAGAAGGGCCCAAGGCTAGGGCCCTGAAGA<br>AGGGCCCGCAGGGCCCTGAAGAAGGGCCCTTTTTTTTACTAGAGCCAATTATTGAAGGCCG<br>CTAACGCGGCCTTTTTTTGTTTCTGGTCTCCC<br>(Scar, Scaffold, anti-repeat, bulge, Terminator L3S3P22) |
| tracrRNA-ESI | tracrRNA (This study) | ACTAGT**GAGACC**CGGAACCATTCAAAGCTTTAAGGCTAGTC**GGTCTC**ACGTTATCAACTTG<br>AAAAAGTGGCACCGAGTCGGTGCGGGCCCTGAAGAAGGGCCCAAGGCTAGGGCCCTGAAGA<br>AGGGCCCGCAGGGCCCTGAAGAAGGGCCCTTTTTTTTACTAGAGCTCGGTACCAAATTCCA<br>GAAAAGAGGCCTCCCGAAAGGGGGGCCTTTTTTCGTTTTGGTCC<br>(Scar, Scaffold, **BsaI site**, Terminator L3S2P21) |
| tracrRNA-R1 | tracrRNA (This study) | CTAGAGGAACCATTCCTTGTAGATGAACAAGTGCCGTTAAGGCTAGTCCGTTATCAACTTG<br>AAAAAGTGGCACCGAGTCGGTGCGGGCCCTGAAGAAGGGCCCAAGGCTAGGGCCCTGAAGA<br>AGGGCCCGCAGGGCCCTGAAGAAGGGCCCTTTTTTTTACTAGAGCTCGGTACCAAATTCCA<br>GAAAAGAGGCCTCCCGAAAGGGGGGCCTTTTTTCGTTTTGGTCC<br>(Scar, Scaffold, anti-repeat, bulge, Terminator L3S2P21) |
| tracrRNA-R2 | tracrRNA (This study) | CTAGAGGAACCATTGCTCGGAGGAGGCCAAGTCCAGCTAAGGCTAGTCCGTTATCAACTTG<br>AAAAAGTGGCACCGAGTCGGTGCGGGCCCTGAAGAAGGGCCCAAGGCTAGGGCCCTGAAGA<br>AGGGCCCGCAGGGCCCTGAAGAAGGGCCCTTTTTTTTACTAGAGCTCGGTACCAAATTCCA<br>GAAAAGAGGCCTCCCGAAAGGGGGGCCTTTTTTCGTTTTGGTCC<br>(Scar, Scaffold, anti-repeat, bulge, Terminator L3S2P21) |
| tracrRNA-R3 | tracrRNA (This study) | CTAGAGGAACCATTAGGTGGTCTTGACCAAGTAGCGTTAAGGCTAGTCCGTTATCAACTTG<br>AAAAAGTGGCACCGAGTCGGTGCGGGCCCTGAAGAAGGGCCCAAGGCTAGGGCCCTGAAGA<br>AGGGCCCGCAGGGCCCTGAAGAAGGGCCCTTTTTTTTACTAGAGCTCGGTACCAAATTCCA<br>GAAAAGAGGCCTCCCGAAAGGGGGGCCTTTTTTCGTTTTGGTCC<br>(Scar, Scaffold, anti-repeat, bulge, Terminator L3S2P21) |
| tracrRNA-R2L1 | tracrRNA (This study) | CTAGAGGAACCATTCTCGGAGGAGGCCAAGTCCAGCTAAGGCTAGTCCGTTATCAACTTGA<br>AAAAGTGGCACCGAGTCGGTGCGGGCCCTGAAGAAGGGCCCAAGGCTAGGGCCCTGAAGAA<br>GGGCCCGCAGGGCCCTGAAGAAGGGCCCTTTTTTTTACTAGAGCTCGGTACCAAATTCCAG<br>AAAAGAGGCCTCCCGAAAGGGGGGCCTTTTTTCGTTTTGGTCC<br>(Scar, Scaffold, anti-repeat, bulge, Terminator L3S2P21) |
| tracrRNA-R2L3 | tracrRNA (This study) | CTAGAGGAACCATTCGGAGGAGGCCAAGTCCAGCTAAGGCTAGTCCGTTATCAACTTGAAA<br>AAGTGGCACCGAGTCGGTGCGGGCCCTGAAGAAGGGCCCAAGGCTAGGGCCCTGAAGAAGG<br>GCCCGCAGGGCCCTGAAGAAGGGCCCTTTTTTTTACTAGAGCTCGGTACCAAATTCCAGAA<br>AAGAGGCCTCCCGAAAGGGGGGCCTTTTTTCGTTTTGGTCC<br>(Scar, Scaffold, anti-repeat, bulge, Terminator L3S2P21) |
| tracrRNA-R2L5 | tracrRNA (This study) | CTAGAGGAACCATTGAGGAGGCCAAGTCCAGCTAAGGCTAGTCCGTTATCAACTTGAAAAA<br>GTGGCACCGAGTCGGTGCGGGCCCTGAAGAAGGGCCCAAGGCTAGGGCCCTGAAGAAGGGC<br>CCGCAGGGCCCTGAAGAAGGGCCCTTTTTTTTACTAGAGCTCGGTACCAAATTCCAGAAAA<br>GAGGCCTCCCGAAAGGGGGGCCTTTTTTCGTTTTGGTCC<br>(Scar, Scaffold, anti-repeat, bulge, Terminator L3S2P21) |
| tracrRNA-R2L7 | tracrRNA (This study) | CTAGAGGAACCATTGGAGGCCAAGTCCAGCTAAGGCTAGTCCGTTATCAACTTGAAAAAGT<br>GGCACCGAGTCGGTGCGGGCCCTGAAGAAGGGCCCAAGGCTAGGGCCCTGAAGAAGGGCCC<br>GCAGGGCCCTGAAGAAGGGCCCTTTTTTTTACTAGAGCTCGGTACCAAATTCCAGAAAAGA<br>GGCCTCCCGAAAGGGGGGCCTTTTTTCGTTTTGGTCC<br>(Scar, Scaffold, anti-repeat, bulge, Terminator L3S2P21) |
| tracrRNA-R2L9 | tracrRNA (This study) | CTAGAGGAACCATTAGGCCAAGTCCAGCTAAGGCTAGTCCGTTATCAACTTGAAAAAGTGG<br>CACCGAGTCGGTGCGGGCCCTGAAGAAGGGCCCAAGGCTAGGGCCCTGAAGAAGGGCCCGC<br>AGGGCCCTGAAGAAGGGCCCTTTTTTTTACTAGAGCTCGGTACCAAATTCCAGAAAAGAGG<br>CCTCCCGAAAGGGGGGCCTTTTTTCGTTTTGGTCC |

| Part name | Type and source | DNA sequence (5'– 3') |
| --- | --- | --- |
| | | (Scar, Scaffold, anti-repeat, bulge, Terminator L3S2P21) |
| tracrRNA-R2L11 | tracrRNA (This study) | CTAGAGGAACCATTGCCAAGTCCAGCTAAGGCTAGTCCGTTATCAACTTGAAAAAGTGGCA<br>CCGAGTCGGTGCGGGCCCTGAAGAAGGGCCCAAGGCTAGGGCCCTGAAGAAGGGCCCGCAG<br>GGCCCTGAAGAAGGGCCCTTTTTTTTACTAGAGCTCGGTACCAAATTCCAGAAAAGAGGCC<br>TCCCGAAAGGGGGGCCTTTTTTCGTTTTGGTCC<br>(Scar, Scaffold, anti-repeat, bulge, Terminator L3S2P21) |
| tracrRNA-R3L1 | tracrRNA (This study) | CTAGAGGAACCATTGGTGGTCTTGACCAAGTAGCGTTAAGGCTAGTCCGTTATCAACTTGA<br>AAAAGTGGCACCGAGTCGGTGCGGGCCCTGAAGAAGGGCCCAAGGCTAGGGCCCTGAAGAA<br>GGGCCCGCAGGGCCCTGAAGAAGGGCCCTTTTTTTTACTAGAGCTCGGTACCAAATTCCAG<br>AAAAGAGGCCTCCCGAAAGGGGGGCCTTTTTTCGTTTTGGTCC<br>(Scar, Scaffold, anti-repeat, bulge, Terminator L3S2P21) |
| tracrRNA-R3L3 | tracrRNA (This study) | CTAGAGGAACCATTTGGTCTTGACCAAGTAGCGTTAAGGCTAGTCCGTTATCAACTTGAAA<br>AAGTGGCACCGAGTCGGTGCGGGCCCTGAAGAAGGGCCCAAGGCTAGGGCCCTGAAGAAGG<br>GCCCGCAGGGCCCTGAAGAAGGGCCCTTTTTTTTACTAGAGCTCGGTACCAAATTCCAGAA<br>AAGAGGCCTCCCGAAAGGGGGGCCTTTTTTCGTTTTGGTCC<br>(Scar, Scaffold, anti-repeat, bulge, Terminator L3S2P21) |
| tracrRNA-R3L5 | tracrRNA (This study) | CTAGAGGAACCATTGTCTTGACCAAGTAGCGTTAAGGCTAGTCCGTTATCAACTTGAAAAA<br>GTGGCACCGAGTCGGTGCGGGCCCTGAAGAAGGGCCCAAGGCTAGGGCCCTGAAGAAGGGC<br>CCGCAGGGCCCTGAAGAAGGGCCCTTTTTTTTACTAGAGCTCGGTACCAAATTCCAGAAAA<br>GAGGCCTCCCGAAAGGGGGGCCTTTTTTCGTTTTGGTCC<br>(Scar, Scaffold, anti-repeat, bulge, Terminator L3S2P21) |
| tracrRNA-R3L7 | tracrRNA (This study) | CTAGAGGAACCATTCTTGACCAAGTAGCGTTAAGGCTAGTCCGTTATCAACTTGAAAAAGT<br>GGCACCGAGTCGGTGCGGGCCCTGAAGAAGGGCCCAAGGCTAGGGCCCTGAAGAAGGGCCC<br>GCAGGGCCCTGAAGAAGGGCCCTTTTTTTTACTAGAGCTCGGTACCAAATTCCAGAAAAGA<br>GGCCTCCCGAAAGGGGGGCCTTTTTTCGTTTTGGTCC<br>(Scar, Scaffold, anti-repeat, bulge, Terminator L3S2P21) |
| tracrRNA-R3L9 | tracrRNA (This study) | CTAGAGGAACCATTTGACCAAGTAGCGTTAAGGCTAGTCCGTTATCAACTTGAAAAAGTGG<br>CACCGAGTCGGTGCGGGCCCTGAAGAAGGGCCCAAGGCTAGGGCCCTGAAGAAGGGCCCGC<br>AGGGCCCTGAAGAAGGGCCCTTTTTTTTACTAGAGCTCGGTACCAAATTCCAGAAAAGAGG<br>CCTCCCGAAAGGGGGGCCTTTTTTCGTTTTGGTCC<br>(Scar, Scaffold, anti-repeat, bulge, Terminator L3S2P21) |
| tracrRNA-R3L11 | tracrRNA (This study) | CTAGAGGAACCATTACCAAGTAGCGTTAAGGCTAGTCCGTTATCAACTTGAAAAAGTGGCA<br>CCGAGTCGGTGCGGGCCCTGAAGAAGGGCCCAAGGCTAGGGCCCTGAAGAAGGGCCCGCAG<br>GGCCCTGAAGAAGGGCCCTTTTTTTTACTAGAGCTCGGTACCAAATTCCAGAAAAGAGGCC<br>TCCCGAAAGGGGGGCCTTTTTTCGTTTTGGTCC<br>(Scar, Scaffold, anti-repeat, bulge, Terminator L3S2P21) |
| tracrRNA-Ar1 | tracrRNA (This study) | CTAGAGGAACCATTAGCTGCAATGGCGCAAGTACCGGTAAGGCTAGTCCGTTATCAACTTG<br>AAAAAGTGGCACCGAGTCGGTGCGGGCCCTGAAGAAGGGCCCAAGGCTAGGGCCCTGAAGA<br>AGGGCCCGCAGGGCCCTGAAGAAGGGCCCTTTTTTTTACTAGAGCTCGGTACCAAATTCCA<br>GAAAAGAGGCCTCCCGAAAGGGGGGCCTTTTTTCGTTTTGGTCC<br>(Scar, Scaffold, anti-repeat, bulge, Terminator L3S2P21) |
| tracrRNA-Ar2 | tracrRNA (This study) | CTAGAGGAACCATTAATGGTAATGTTGCAAGTATATCTAAGGCTAGTCCGTTATCAACTTG<br>AAAAAGTGGCACCGAGTCGGTGCGGGCCCTGAAGAAGGGCCCAAGGCTAGGGCCCTGAAGA<br>AGGGCCCGCAGGGCCCTGAAGAAGGGCCCTTTTTTTTACTAGAGCTCGGTACCAAATTCCA<br>GAAAAGAGGCCTCCCGAAAGGGGGGCCTTTTTTCGTTTTGGTCC<br>(Scar, Scaffold, anti-repeat, bulge, Terminator L3S2P21) |
| mRFP | mRNA (This study) | CTAGAG**ATTAAAGAGGAGAAA**TACTAGATGGTGAGCAAGGGCGAGGAGGATAACATGGCCA<br>TCATCAAGGAGTTCATGCGCTTCAAGGTGCACATGGAGGGCTCCGTGAACGGCCACGAGTT<br>CGAGATCGAGGGCGAGGGCGAGGGCCGCCCCTACGAGGGCACCCAGACCGCCAAGCTGAAG<br>GTGACCAAGGGTGGCCCCCTGCCCTTCGCCTGGGACATCCTGTCCCCTCAGTTCATGTACG<br>GCTCCAAGGCCTACGTGAAGCACCCCGCCGACATCCCCGACTACTTGAAGCTGTCCTTCCC<br>CGAGGGCTTCAAGTGGGAGCGCGTGATGAACTTCGAGGACGGCGGCGTGGTGACCGTGACC<br>CAGGACTCCTCACTTCAGGACGGCGAGTTCATCTACAAGGTGAAGCTGCGCGGCACCAACT<br>TCCCCTCCGACGGCCCCGTAATGCAGAAGAAGACCATGGGCTGGGAGGCCTCCTCCGAGCG<br>GATGTACCCCGAGGACGGCGCCCTGAAGGGCGAGATCAAGCAGAGGCTGAAGCTGAAGGAC<br>GGCGGCCACTACGACGCTGAGGTCAAGACCACCTACAAGGCCAAGAAGCCCGTGCAGCTGC<br>CCGGCGCCTACAACGTCAACATCAAGTTGGACATCACCTCCCACAACGAGGACTACACCAT |

| Part name | Type and source | DNA sequence (5’– 3’) |
|---|---|---|
| | | CGTGGAACAGTACGAACGCGCTGAGGGCCGCCACTCCACCGGCGGCATGGACGAGCTGTAC<br>AAGTAACCAATTATTGAAGGGGAGCGGGAAACCGCTCCCCTTTTTTTGTTTCTGGTCTCCC<br>(Scar, coding, Terminator L3S3P00) |
| mRFP-ΔRBS | mRNA<br>(This study) | CTAGAGGTGAGCAAGGGCGAGGAGGATAACATGGCCATCATCAAGGAGTTCATGCGCTTCA<br>AGGTGCACATGGAGGGCTCCGTGAACGGCCACGAGTTCGAGATCGAGGGCGAGGGCGAGGG<br>CCGCCCCTACGAGGGCACCCAGACCGCCAAGCTGAAGGTGACCAAGGGTGGCCCCCTGCCC<br>TTCGCCTGGGACATCCTGTCCCCTCAGTTCATGTACGGCTCCAAGGCCTACGTGAAGCACC<br>CCGCCGACATCCCCGACTACTTGAAGCTGTCCTTCCCCGAGGGCTTCAAGTGGGAGCGCGT<br>GATGAACTTCGAGGACGGCGGCGTGGTGACCGTGACCCAGGACTCCTCACTTCAGGACGGC<br>GAGTTCATCTACAAGGTGAAGCTGCGCGGCACCAACTTCCCCTCCGACGGCCCCGTAATGC<br>AGAAGAAGACCATGGGCTGGGAGGCCTCCTCCGAGCGGATGTACCCCGAGGACGGCGCCCT<br>GAAGGGCGAGATCAAGCAGAGGCTGAAGCTGAAGGACGGCGGCCACTACGACGCTGAGGTC<br>AAGACCACCTACAAGGCCAAGAAGCCCGTGCAGCTGCCCGGCGCCTACAACGTCAACATCA<br>AGTTGGACATCACCTCCCACAACGAGGACTACACCATCGTGGAACAGTACGAACGCGCTGA<br>GGGCCGCCACTCCACCGGCGGCATGGACGAGCTGTACAAGTAACCAATTATTGAAGGGGAG<br>CGGGAAACCGCTCCCCTTTTTTTGTTTCTGGTCTCCC<br>(Scar, partial coding, Terminator L3S3P00) |
| crRNA-R1 | crRNA<br>(This study) | CTAGACCCAGGACTCCTCACTTCAGGACGGCGAGTTCATCTACAAGGTACTAGAGCCAATT<br>ATTGAAGGGGAGCGGGAAACCGCTCCCCTTTTTTTGTTTCTGGTCTCCC<br>(Scar, Terminator L3S3P00) |
| crRNA-R2 | crRNA<br>(This study) | CTAGATAATGCAGAAGAAGACCATGGGCTGGGAGGCCTCCTCCGAGCTACTAGAGCCAATT<br>ATTGAAGGGGAGCGGGAAACCGCTCCCCTTTTTTTGTTTCTGGTCTCCC<br>(Scar, Terminator L3S3P00) |
| crRNA-R3 | crRNA<br>(This study) | CTAGATGAAGGACGGCGGCCACTACGACGCTGAGGTCAAGACCACCTTACTAGAGCCAATT<br>ATTGAAGGGGAGCGGGAAACCGCTCCCCTTTTTTTGTTTCTGGTCTCCC<br>(Scar, Terminator L3S3P00) |
| mRNA-Ar | mRNA<br>(This study) | CTAGACAATCAGGAGCGCAATATGTCATTTCTGTTACCCATCCAATTGTTCAAAATTCTTG<br>CTGATGAAACCCGTCTGGGCATCGTTTTACTGCTCAGCGAACTGGGAGAGTTATGCGTCTG<br>CGATCTCTGCACTGCTCTCGACCAGTCGCAGCCCAAGATCTCCCGCCACCTGGCATTGCTG<br>CGTGAAAGCGGGCTATTGCTGGACCGCAAGCAAGGTAAGTGGGTTCATTACCGCTTATCAC<br>CGCATATTCCAGCATGGGCGGCGAAAATTATTGATGAGGCCTGGCGATGTGAACAGGAAAA<br>GGTTCAGGCGATTGTCCGCAACCTGGCTCGACAAAACTGTTCCGGGGACAGTAAGAACATT<br>TGCAGTTAAAAATTTAGCTAAACACATATGAATTTTCAGATGTGTTTTATCCGGGAGGCAT<br>TATGTTACTGGCAGGCGCTATCTTTGTCCTGACCATCGTATTGGTTATCTGGCAGCCGAAA<br>GGTTTAGGCATCGGCTGGAGTGCAACGCTCGGCGCAGTACTGGCGTTAGTTACGGGCGTGG<br>TCCATCCGGGTGATATTCCGGTGGTGTGGAATATCGTCTGGAACGCGACGGCTGCGTTTAT<br>CGCCGTCATTATCATCAGCCTGCTGCTGGATGAGTCCGGCTTTTTTGAATGGGCGGCGCTG<br>CACGTCTCACGCTGGGGTAATGGTCGTGGTCGCTTGCTGTTTACCTGGATTGTCCTGCTCG<br>GTGCTGCCGTTGCCGCCCTGTTTGCCAATGATGGCGCGGCGCTTATTTTGACACCGATTGT<br>CATCGCCATGCTGCTGGCTTTAGGGTTCAGTAAAGGCACTACGCTGGCGTTCGTGATGGCG<br>GCCGGATTCATTGCCGATACCGCCAGCCTGCCGCTTATTGTCTCCAACCTGGTGAATATCG<br>TTTCCGCTGATTTCTTTGGCCTCGGCTTTCGCGAATACGCCTCGGTGATGGTGCCGGTGGA<br>TATCGCCGCGATTGTTGCCACGCTGGTGATGTTACATCTCTATTTTCGCAAAGATATTCCG<br>CAGAACTACGATATGGCGCTGCTGAAATCTCCCGCAGAAGCGATCAAAGATCCTGCTACGT<br>TCAAAACTGGCTGGGTTGTTTTACTGCTTCTGCTGGTGGGATTTTTCGTCCTGGAACCGCT<br>CGGCATTCCGGTGAGCGCCATTGCAGCTGTGGGCGCGCTGATATTATTTGTCGTCGCTAAA<br>CGCGGTCATGCGATTAATACGGGTAAAGTCCTGCGCGGTGCCCCCTGGCAGATTGTCATCT<br>TCTCGCTCGGCATGTATCTGGTGGTTTATGGCCTGCGCAATGCCGGATTAACGGAATATCT<br>TTCTGGCGTACTCAACGTGCTGGCGGATAACGGCCTGTGGGCCGCGACGCTCGGCACCGGA<br>TTCCTCACCGCCTTCCTCTCTTCTATTATGAACAATATGCCGACGGTACTGGTTGGCGCGT<br>TGTCCATTGATGGCAGCACGGCATCTGGCGTTATCAAAGAAGCGATGGTTTATGCCAATGT<br>GATTGGCTGCGATTTGGGACCGAAAATTACCCCAATTGGTAGCCTGGCTACGCTACTCTGG<br>CTGCACGTACTTTCGCAGAAGAATATGACTATCAGCTGGGGATATTACTTCCGTACAGGGA<br>TTATCATGACCCTGCCTGTGCTGTTTGTGACGCTGGCTGCGCTGGCGCTACGTCTCTCTTT<br>CACTTTGTAATGAGATACTGATATGAGCAACATTACCATTTATCACAACCCGGCCTGCGGC<br>ACGTCGCGTAATACGCTGGAGATGATCCGCAACAGCGGCACAGAACCGACTATTATCCATT<br>ATCTGGAAACTCCGCCAACGCGCGATGAACTGGTCAAACTCATTGCCGATATGGGGATTTC<br>CGTACGCGCGCTGCTGCGTAAAAACGTCGAACCGTATGAGGAGCTGGGCCTTGCGGAAGAT<br>AAATTTACTGACGATCGGTTAATCGACTTTATGCTTCAGCACCCGATTCTGATTAATCGCC<br>CGATTGTGGTGACGCCGCTGGGAACTCGCCTGTGCCGCCCTTCAGAAGTGGTGCTGGAAAT<br>TCTGCCAGATGCGCAAAAAGGCGCATTCTCCAAGGAAGATGGCGAGAAAGTGGTTGATGAA<br>GCGGGTAAGCGCCTGAAATAATACTAGAGCCAATTATTGAAGGGGAGCGGGAAACCGCTCC<br>CCTTTTTTTGTTTCTGGTCTCCC |

| Part name | Type and source | DNA sequence (5'– 3') |
|---|---|---|
| | | (Scar, Terminator L3S3P00) |
| crRNA-Ar1 | crRNA<br>(This study) | CTAGAGTCCTGGAACCGCTCGGCATTCCGGTGAGCGCCATTGCAGCTTACTAGAGCCAATT<br>ATTGAAGGGGAGCGGGAAACCGCTCCCCTTTTTTTGTTTCTGGTCTCCC<br>(Scar, Terminator L3S3P00) |
| crRNA-Ar2 | crRNA<br>(This study) | CTAGACTAGATTCACTTTGTAATGAGATACTGATATGAGCAACATTACCATTTACTAGAGC<br>CAATTATTGAAGGGGAGCGGGAAACCGCTCCCCTTTTTTTGTTTCTGGTCTCCC<br>(Scar, Terminator L3S3P00) |
| crRNA-Ar3 | crRNA<br>(This study) | CTAGACTAGATGCTGCGTAAAAACGTCGAACCGTATGAGGAGCTGGGCCTTGTACTAGAGC<br>CAATTATTGAAGGGGAGCGGGAAACCGCTCCCCTTTTTTTGTTTCTGGTCTCCC<br>(Scar, Terminator L3S3P00) |
| crRNA-Ar4 | crRNA<br>(This study) | CTAGACTAGAAAGGCGCATTCTCCAAGGAAGATGGCGAGAAAGTGGTTGATGTACTAGAGC<br>CAATTATTGAAGGGGAGCGGGAAACCGCTCCCCTTTTTTTGTTTCTGGTCTCCC<br>(Scar, Terminator L3S3P00) |
| s-crRNA-LEB3 | crRNA<br>(This study) | CTAGACATAGTTCGTTTCCCATGTTTTAGAGCTATGCCCAATTATTGAAGGCCGCTAACGC<br>GGCCTTTTTTTGTTTCTGGTCTCCC<br>(Scar, Terminator L3S3P22) |
| s-crRNA-LEA2-WT | crRNA<br>(This study) | CTAGAAGTTATTATATAGTTCGGTCGTTTTAGAGCTATGCCCAATTATTGAAGGCCGCTAA<br>CGCGGCCTTTTTTTGTTTCTGGTCTCCC<br>(Scar, Terminator L3S3P22) |
| s-crRNA-LEA2-C6 | crRNA<br>(This study) | CTAGAAGTTATTATATAGTTCGGTCTACGTACCGGGTGAACCAATTATTGAAGGCCGCTAA<br>CGCGGCCTTTTTTTGTTTCTGGTCTCCC<br>(Scar, Terminator L3S3P22) |
| s-crRNA-LEA2-C9 | crRNA<br>(This study) | CTAGAAGTTATTATATAGTTCGGTCAGAATCCTCGCTCAGCCAATTATTGAAGGCCGCTAA<br>CGCGGCCTTTTTTTGTTTCTGGTCTCCC<br>(Scar, Terminator L3S3P22) |
| s-crRNA-LEA2-C48 | crRNA<br>(This study) | CTAGAAGTTATTATATAGTTCGGTCATTATTGCCGACACACCAATTATTGAAGGCCGCTAA<br>CGCGGCCTTTTTTTGTTTCTGGTCTCCC<br>(Scar, Terminator L3S3P22) |
| s-crRNA-LEA2-C74 | crRNA<br>(This study) | CTAGAAGTTATTATATAGTTCGGTCAATCGGGAGCTGCCTCCAATTATTGAAGGCCGCTAA<br>CGCGGCCTTTTTTTGTTTCTGGTCTCCC<br>(Scar, Terminator L3S3P22) |
| s-tracrRNA-WT | tracrRNA<br>(This study) | CTAGGCATAGCAAGTTAAAATAAGGCTAGTCCGTTATCAACTTGAAAAAGTGGCACCGAGT<br>CGGTGCGGGCCCTGAAGAAGGGCCCAAGGCTAGGGCCCTGAAGAAGGGCCCGCAGGGCCCT<br>GAAGAAGGGCCCTTTTTTTTACTAGAGCTCGGTACCAAATTCCAGAAAAGAGGCCTCCCGA<br>AAGGGGGGCCTTTTTTCGTTTTGGTCC<br>(Scar,Scaffold,Terminator L3S2P21) |
| s-tracrRNA-C6 | tracrRNA<br>(This study) | CTAGTTCACCCAAGTTACGTTAAGGCTAGTCCGTTATCAACTTGAAAAAGTGGCACCGAGT<br>CGGTGCGGGCCCTGAAGAAGGGCCCAAGGCTAGGGCCCTGAAGAAGGGCCCGCAGGGCCCT<br>GAAGAAGGGCCCTTTTTTTTACTAGAGCTCGGTACCAAATTCCAGAAAAGAGGCCTCCCGA<br>AAGGGGGGCCTTTTTTCGTTTTGGTCC<br>(Scar,Scaffold,Terminator L3S2P21) |
| s-tracrRNA-C9 | tracrRNA<br>(This study) | CTAGCTGAGCGAAGTGATTCTAAGGCTAGTCCGTTATCAACTTGAAAAAGTGGCACCGAGT<br>CGGTGCGGGCCCTGAAGAAGGGCCCAAGGCTAGGGCCCTGAAGAAGGGCCCGCAGGGCCCT<br>GAAGAAGGGCCCTTTTTTTTACTAGAGCTCGGTACCAAATTCCAGAAAAGAGGCCTCCCGA<br>AAGGGGGGCCTTTTTTCGTTTTGGTCC<br>(Scar,Scaffold,Terminator L3S2P21) |
| s-tracrRNA-C48 | tracrRNA<br>(This study) | CTAGTGTGTCGAAGTAATAATAAGGCTAGTCCGTTATCAACTTGAAAAAGTGGCACCGAGT<br>CGGTGCGGGCCCTGAAGAAGGGCCCAAGGCTAGGGCCCTGAAGAAGGGCCCGCAGGGCCCT<br>GAAGAAGGGCCCTTTTTTTTACTAGAGCTCGGTACCAAATTCCAGAAAAGAGGCCTCCCGA<br>AAGGGGGGCCTTTTTTCGTTTTGGTCC<br>(Scar,Scaffold,Terminator L3S2P21) |
| s-tracrRNA-C74 | tracrRNA<br>(This study) | CTAGAGGCAGCAAGTCCGATTAAGGCTAGTCCGTTATCAACTTGAAAAAGTGGCACCGAGT<br>CGGTGCGGGCCCTGAAGAAGGGCCCAAGGCTAGGGCCCTGAAGAAGGGCCCGCAGGGCCCT |

| Part name | Type and source | DNA sequence (5'– 3') |
|---|---|---|
| | | GAAGAAGGGCCCTTTTTTTTACTAGAGCTCGGTACCAAATTCCAGAAAAGAGGCCTCCCGA<br>AAGGGGGGCCTTTTTTCGTTTTGGTCC<br>(Scar,Scaffold,Terminator L3S2P21) |
| s-tracrRNA-R1 | tracrRNA (This study) | CCTTGTAGATGAACAAGTGCCGTTAAGGCTAGTCCGTTATCAACTTGAAAAAGTGGCACCG<br>AGTCGGTGCGGGCCCTGAAGAAGGGCCCAAGGCTAGGGCCCTGAAGAAGGGCCCGCAGGGC<br>CCTGAAGAAGGGCCCTTTTTTTTACTAGAGCTCGGTACCAAATTCCAGAAAAGAGGCCTCC<br>CGAAAGGGGGGCCTTTTTTCGTTTTGGTCC<br>(Scar,Scaffold,Terminator L3S2P21) |
| s-tracrRNA-R2 | tracrRNA (This study) | GCTCGGAGGAGGCCAAGTCCAGCTAAGGCTAGTCCGTTATCAACTTGAAAAAGTGGCACCG<br>AGTCGGTGCGGGCCCTGAAGAAGGGCCCAAGGCTAGGGCCCTGAAGAAGGGCCCGCAGGGC<br>CCTGAAGAAGGGCCCTTTTTTTTACTAGAGCTCGGTACCAAATTCCAGAAAAGAGGCCTCC<br>CGAAAGGGGGGCCTTTTTTCGTTTTGGTCC<br>(Scar,Scaffold,Terminator L3S2P21) |
| s-tracrRNA-R3 | tracrRNA (This study) | AGGTGGTCTTGACCAAGTAGCGTTAAGGCTAGTCCGTTATCAACTTGAAAAAGTGGCACCG<br>AGTCGGTGCGGGCCCTGAAGAAGGGCCCAAGGCTAGGGCCCTGAAGAAGGGCCCGCAGGGC<br>CCTGAAGAAGGGCCCTTTTTTTTACTAGAGCTCGGTACCAAATTCCAGAAAAGAGGCCTCC<br>CGAAAGGGGGGCCTTTTTTCGTTTTGGTCC<br>(Scar,Scaffold,Terminator L3S2P21) |
| CONAN | Sensor RNA | TTGCCATGTGTATGTGGGAGACGGTCGGGTCCAGATATTCGTATCTGTCGAGTAGAGTGTG<br>GGCTCCCACATACTCTGATGATCCTTCGGGATCATTCATGGCAACTGCTGCTCTTCAACCT<br>CGGTGACGAGGTTGAAGAGCAGCAGTTGCCATGTGTATGTGGGAGCCCACACTCTACTCGA<br>CAGATACGTAGCATAACCCCTTGGGGCCTCTAAACGGGTCTTGAGGGGTTTTTTG<br>(Target, broccoli RNA ,PAM ,Terminator T7) |
| s-crRNA-13 | crRNA (This study) | AGTTATTATATAGTTCGGTCGTTTTAGAGCTATGCCAATTATTGAAGGCCGCTAACGCGGC<br>CTTTTTTTGTTTCTGGTCTCCC<br>(Spacer ,Terminator L3S3P22) |
| s-crRNA-12 | crRNA (This study) | AGTTATTATATAGTTCGGTCGTTTTAGAGCTATCCAATTATTGAAGGCCGCTAACGCGGCC<br>TTTTTTTGTTTCTGGTCTCCC<br>(Spacer ,Terminator L3S3P22) |
| s-crRNA-11 | crRNA (This study) | AGTTATTATATAGTTCGGTCGTTTTAGAGCTACCAATTATTGAAGGCCGCTAACGCGGCCT<br>TTTTTTGTTTCTGGTCTCCC<br>(Spacer ,Terminator L3S3P22) |
| s-crRNA-10 | crRNA (This study) | AGTTATTATATAGTTCGGTCGTTTTAGAGCTCCAATTATTGAAGGCCGCTAACGCGGCCTT<br>TTTTTGTTTCTGGTCTCCC<br>(Spacer ,Terminator L3S3P22) |
| s-tracrRNA-13 | tracrRNA (This study) | CATAGCAAGTTAAAATAAGGCTAGTCCGTTATCAACTTGAAAAAGTGGCACCGAGTCGGTG<br>CGGGCCCTGAAGAAGGGCCCAAGGCTAGGGCCCTGAAGAAGGGCCCGCAGGGCCCTGAAGA<br>AGGGCCCTTTTTTTTACTAGAGCTCGGTACCAAATTCCAGAAAAGAGGCCTCCCGAAAGGG<br>GGGCCTTTTTTCGTTTTGGTCC<br>(Scar,Scaffold,Terminator L3S2P21) |
| s-tracrRNA-12 | tracrRNA (This study) | ATAGCAAGTTAAAATAAGGCTAGTCCGTTATCAACTTGAAAAAGTGGCACCGAGTCGGTGC<br>GGGCCCTGAAGAAGGGCCCAAGGCTAGGGCCCTGAAGAAGGGCCCGCAGGGCCCTGAAGAA<br>GGGCCCTTTTTTTTACTAGAGCTCGGTACCAAATTCCAGAAAAGAGGCCTCCCGAAAGGGG<br>GGCCTTTTTTCGTTTTGGTCC<br>(Scar,Scaffold,Terminator L3S2P21) |
| s-tracrRNA-11 | tracrRNA (This study) | TAGCAAGTTAAAATAAGGCTAGTCCGTTATCAACTTGAAAAAGTGGCACCGAGTCGGTGCG<br>GGCCCTGAAGAAGGGCCCAAGGCTAGGGCCCTGAAGAAGGGCCCGCAGGGCCCTGAAGAAG<br>GGCCCTTTTTTTTACTAGAGCTCGGTACCAAATTCCAGAAAAGAGGCCTCCCGAAAGGGGG<br>GCCTTTTTTCGTTTTGGTCC<br>(Scar,Scaffold,Terminator L3S2P21) |
| s-tracrRNA-10 | tracrRNA (This study) | AGCAAGTTAAAATAAGGCTAGTCCGTTATCAACTTGAAAAAGTGGCACCGAGTCGGTGCGG<br>GCCCTGAAGAAGGGCCCAAGGCTAGGGCCCTGAAGAAGGGCCCGCAGGGCCCTGAAGAAGG<br>GCCCTTTTTTTTACTAGAGCTCGGTACCAAATTCCAGAAAAGAGGCCTCCCGAAAGGGGGG<br>CCTTTTTTCGTTTTGGTCC<br>(Scar,Scaffold,Terminator L3S2P21) |

| Part name | Type and source | DNA sequence (5'– 3') |
|---|---|---|
| s-tracrRNA-Ar2 | tracrRNA (This study) | AATGGTAATGTTGCAAGTATATCTAAGGCTAGTCCGTTATCAACTTGAAAAAGTGGCACCG AGTCGGTGCGGGCCCTGAAGAAGGGCCCAAGGCTAGGGCCCTGAAGAAGGGCCCGCAGGGC CCTGAAGAAGGGCCCTTTTTTTTACTAGAGCTCGGTACCAAATTCCAGAAAAGAGGCCTCC CGAAAGGGGGGCCTTTTTTCGTTTTGGTCC (Scar,Scaffold,Terminator L3S2P21) |
| tracrRNA- Ar3 | tracrRNA (This study) | CTAGAGGAACCATTCAAGGCCCAGCTCCAAGTATACGTAAGGCTAGTCCGTTATCAACTTG AAAAAGTGGCACCGAGTCGGTGCGGGCCCTGAAGAAGGGCCCAAGGCTAGGGCCCTGAAGA AGGGCCCGCAGGGCCCTGAAGAAGGGCCCTTTTTTTTACTAGAGCTCGGTACCAAATTCCA GAAAAGAGGCCTCCCGAAAGGGGGGCCTTTTTTCGTTTTGGTCC (Scar,Scaffold,Terminator L3S2P21) |
| tracrRNA- Ar4 | tracrRNA (This study) | CTAGAGGAACCATTCATCAACCACTTTCAAGTGCCATTAAGGCTAGTCCGTTATCAACTTG AAAAAGTGGCACCGAGTCGGTGCGGGCCCTGAAGAAGGGCCCAAGGCTAGGGCCCTGAAGA AGGGCCCGCAGGGCCCTGAAGAAGGGCCCTTTTTTTTACTAGAGCTCGGTACCAAATTCCA GAAAAGAGGCCTCCCGAAAGGGGGGCCTTTTTTCGTTTTGGTCC (Scar,Scaffold,Terminator L3S2P21) |
| tracrRNA- WT-21T | tracrRNA (This study) | CTAGAGGAACCATTCAAAACAGCATAGCAAGTTAAAATAAGGCTAGTCCGTTATCAACTTG AAAAAGTGGCACCGAGTCGGTGCGGGCCCTGAAGAAGGGCCCAAGGCTAGGGCCCTGAAGA AGGGCCCGCAGGGCCCTGAAGAAGGGCCCTTTTTTTTACTAGAGCTCGGTACCAAATTCCA GAAAAGAGGCCTCCCGAAAGGGGGGCCTTTTTTCGTTTTGGTCC (Scar,Scaffold,Terminator L3S2P21) |

#### Supplementary Table 9: Sequences of promoters

| Part name | Type and source | DNA sequence (5'– 3') |
|---|---|---|
| $P_{pspA}$-LEA2B2 | $\sigma^{54}$-dependent promoter<br>(This study) | AGTTATTATATAGTTCGGTCCGGTTTGAGACGTTGTTTTGCGGTTAGTGTAATTCGCTAAC<br>TCATCCTGGCATGTTGCTGTTGATTCTTCAATCAGATCTTTATAAATCAAAAAGATAAAAA<br>ATTGGCACGCAAATTGTATTAACAGTTCAGCAGGACAATCCTGAACGCAAA<br>(-24 Box, -12 Box, TTS, UAS) |
| $P_{pspA}$-R1 | $\sigma^{54}$-dependent promoter<br>(This study) | CCCAGGACTCCTCACTTCAGCGGTTAGTGTAATTCGCTAACTCATCCTGGCATGTTGCTGT<br>TGATTCTTCAATCAGATCTTTATAAATCAAAAAGATAAAAAAATTGGCACGCAAATTGTATT<br>AACAGTTCAGCAGGACAATCCTGAACGCAAA<br>(-24 Box, -12 Box, TTS, UAS) |
| $P_{pspA}$-R2 | $\sigma^{54}$-dependent promoter<br>(This study) | TAATGCAGAAGAAGACCATGCGGTTAGTGTAATTCGCTAACTCATCCTGGCATGTTGCTGT<br>TGATTCTTCAATCAGATCTTTATAAATCAAAAAGATAAAAAAATTGGCACGCAAATTGTATT<br>AACAGTTCAGCAGGACAATCCTGAACGCAAA<br>(-24 Box, -12 Box, TTS, UAS) |
| $P_{pspA}$-R3 | $\sigma^{54}$-dependent promoter<br>(This study) | TGAAGGACGGCGGCCACTACCGGTTAGTGTAATTCGCTAACTCATCCTGGCATGTTGCTGT<br>TGATTCTTCAATCAGATCTTTATAAATCAAAAAGATAAAAAAATTGGCACGCAAATTGTATT<br>AACAGTTCAGCAGGACAATCCTGAACGCAAA<br>(-24 Box, -12 Box, TTS, UAS) |
| $P_{pspA}$-Ar1 | $\sigma^{54}$-dependent promoter<br>(This study) | GTCCTGGAACCGCTCGGCATCGGTTAGTGTAATTCGCTAACTCATCCTGGCATGTTGCTGT<br>TGATTCTTCAATCAGATCTTTATAAATCAAAAAGATAAAAAAATTGGCACGCAAATTGTATT<br>AACAGTTCAGCAGGACAATCCTGAACGCAAA<br>(-24 Box, -12 Box, TTS, UAS) |
| $P_{pspA}$-Ar2 | $\sigma^{54}$-dependent promoter<br>(This study) | TTCACTTTGTAATGAGATACCGGTTAGTGTAATTCGCTAACTCATCCTGGCATGTTGCTGT<br>TGATTCTTCAATCAGATCTTTATAAATCAAAAAGATAAAAAAATTGGCACGCAAATTGTATT<br>AACAGTTCAGCAGGACAATCCTGAACGCAAA<br>(-24 Box, -12 Box, TTS, UAS) |
| $P_{pspA}$-Ar3 | $\sigma^{54}$-dependent promoter<br>(This study) | AGTGCTGCGTAAAAACGTCGAACGGTTAGTGTAATTCGCTAACTCATCCTGGCATGTTGCT<br>GTTGATTCTTCAATCAGATCTTTATAAATCAAAAAGATAAAAAAATTGGCACGCAAATTGTA<br>TTAACAGTTCAGCAGGACAATCCTGAACGCAAA<br>(-24 Box, -12 Box, TTS, UAS) |
| $P_{pspA}$-Ar4 | $\sigma^{54}$-dependent promoter<br>(This study) | AGAAGGCGCATTCTCCAAGGAACGGTTAGTGTAATTCGCTAACTCATCCTGGCATGTTGCT<br>GTTGATTCTTCAATCAGATCTTTATAAATCAAAAAGATAAAAAAATTGGCACGCAAATTGTA<br>TTAACAGTTCAGCAGGACAATCCTGAACGCAAA<br>(-24 Box, -12 Box, TTS, UAS) |
| J23106 | Anderson promoter | TTTACGGCTAGCTCAGTCCTAGGTATAGTGCTAGC |
| $P_{T7}$ | T7 promoter | TAATACGACTCACTATAGG |
